# The G protein-coupled receptor GPR31 governs inflammatory signaling in the pancreatic islet microenvironment

**DOI:** 10.1101/2025.10.02.680021

**Authors:** Kerim B. Kaylan, Xia S. Saavedra, Christian Checkcinco, Jacob R. Enriquez, Titli Nargis, Emily Elliott, Armando A. Puente, Jiayi E. Wang, Melissa A. Walsh, Jennifer B. Nelson, Abhishek Kulkarni, Charanya Muralidharan, Sarah C. May, Ryan M. Anderson, Raghavendra G. Mirmira, Sarah A. Tersey

## Abstract

In type 1 diabetes (T1D), immune-mediated destruction of the insulin-secreting pancreatic β cells is amplified by inflammatory signaling within the β-cell itself. The 12-lipoxygenase pathway exemplifies the effects of β-cell-intrinsic signaling, and its product, 12-hydroxyeicosatetraenoic acid (12-HETE) induces cellular stress in β cells and promotes macrophage migration and cytokine release. The G protein-coupled receptor GPR31 is a putative 12-HETE receptor, but its role in islet inflammation is undefined. Here we interrogate GPR31 using germline *Gpr31b* knockout mice, siRNA knockdown, and a small-molecule GPR31 inhibitor. Loss of GPR31 decreases inflammatory and oxidative stress transcriptional programs in islets and reduces migration and interferon signaling in macrophages. Across multiple mouse models of T1D, disrupting GPR31 improves glucose tolerance, decreases insulitis and macrophage infiltration, and reduces islet oxidative stress. Loss of GPR31 also perturbs alternative splicing, regulating differential expression of isoforms involved in the cell type-specific functions of GPR31. In human single-cell sequencing data, *GPR31* marks conventional type 1 dendritic cells of the pancreatic lymph node that carry the same inflammatory, interferon, and splicing signature as murine *Gpr31b*^+^ macrophages. Collectively, these findings establish GPR31 as a key inflammatory node in both β cells and macrophages in T1D.

## INTRODUCTION

In the classical perspective of type 1 diabetes (T1D), loss of immune tolerance triggers a T cell-mediated immune response that destroys the insulin-producing pancreatic β cells. This immune-centric perspective has been challenged in recent years, and emerging evidence shows that other cell types within the pancreas, such as macrophages and β cells, actively participate in the pathogenesis of T1D (1–3). For example, blockade of stress-related or inflammatory signaling cascades within β cells alone prevents the emergence of adaptive immunity and β-cell loss in non-obese diabetic (NOD) mice (1,4). Similarly, blockade of inflammatory responses in early infiltrating macrophages in NOD mice prevents adaptive immunity and overt T1D (5–7). These findings are not limited to mouse models. Clinical studies employing the repurposed drugs verapamil and eflornithine suggest that reductions in β-cell oxidative and endoplasmic reticulum (ER) stress appear to preserve β-cell function in early T1D (8,9). These and other studies emphasize the need to identify pathways and potential targets in β cells and macrophages that may contribute to the initiation or propagation of T1D.

Lipoxygenase (LOX) enzymes are critical regulators of inflammation that catalyze the oxygenation of polyunsaturated fatty acids to form both pro– and anti-inflammatory mediators (10,11). Among enzymes present in both β cells and macrophages, 12/15-LOX converts arachidonic acid (C20:4) to the pro-inflammatory eicosanoid 12(*S*)-hydroxyeicosatetraenoic acid (12(*S*)-HETE or simply 12-HETE) (12,13). In individuals with early stage or established T1D, 12/15-LOX production is notably elevated in pancreatic islets (14), and levels of 12-HETE are also elevated in the sera of individuals with new-onset T1D (15). Chemical inhibition of 12/15-LOX decreases 12-HETE production in cytokine-exposed human islets and improves β-cell function (16). In NOD mice, tissue-specific deletion of the gene encoding 12/15-LOX (*Alox15*) in either β cells or macrophages results in reduced islet inflammation (insulitis) and protection from the development of diabetes in both sexes (4,6). In NOD mice with the human ortholog replacing murine 12/15-LOX, inhibition with the selective inhibitor VLX-1005 delays diabetes onset through reduction in both β-cell stress and inflammatory signaling in macrophages (17). Although the signals transducing 12/15-LOX action have not been fully elucidated, it is known that its primary product, 12-HETE, can activate various pro-inflammatory signaling pathways. These include p38 mitogen-activated protein kinase (MAPK) and c-JUN N-terminal kinase (JNK) (18,19), and the action of these kinases can induce or exacerbate oxidative and ER stress (20).

As a lipid peroxide, 12-HETE may exert pro-oxidative effects in β cells and macrophages as it accumulates to critical levels intracellularly. However, since 12-HETE can be measured in circulation, its effects may not be limited to cell-autonomous functions, raising the possibility that an eicosanoid receptor may mediate systemic effects. The orphan G protein-coupled receptor GPR31 has been shown to function as a receptor for 12-HETE (21). GPR31 displays high affinity (K_d_ ∼4 nM) for 12-HETE and exhibits stereospecificity for its biologically relevant *S-*enantiomer. GPR31 is coupled to G_αq_, and its effects are mediated by phosphatidylinositol signaling and subsequent activation of MAPK, extracellular signal-regulated kinases 1/2 (ERK1/2), and nuclear factor kappa-light-chain-enhancer of activated B cells (NFκB) (21). Consistent with the idea that GPR31 functions as a receptor for 12-HETE, knockdown of its encoding gene in human hepatocytes diminished 12-HETE-induced activation of protein kinase C (PKC)–JNK signaling (22). However, the role of GPR31 in macrophages and β cells in the pathogenesis of T1D remains speculative. In one study, the gene encoding GPR31 (*Gpr31b*) was found to be induced in islet-resident macrophages following high-fat diet feeding of mice (23). Yet, its direct role in macrophage-mediated inflammation was not explored. Another study in zebrafish suggested that GPR31 lies downstream of 12/15-LOX in islets based on the developmental phenotype (24), but its role in β-cell inflammation was not examined. In this study, we tested the hypothesis that GPR31 promotes β-cell and macrophage inflammatory signaling and contributes to β-cell dysfunction in diabetes, using complementary genetic, pharmacologic, and transcriptomic approaches across mouse, zebrafish, and human data.

## RESULTS

### Mice with whole-body Gpr31b knockout exhibit normal body growth and glycemic control

Prior work identified GPR31 as a receptor for 12-HETE (21). Based on previous data showing 12-HETE is produced by pancreatic islet β cells and macrophages, we sought to determine if GPR31 is also expressed in islets and macrophages. In mice, GPR31 is encoded by the gene *Gpr31b*, but there are two non-functional *Gpr31b* pseudogenes (*Gpr31a* and *Gpr31c*) that do not encode protein. By quantitative PCR of tissues from wildtype (WT) *C57BL/6J* mice, an assay that does not distinguish *Gpr31b* from its pseudogenes, we found *Gpr31a/b/c* mRNA to be expressed predominantly in the islets with lower expression in spleen, bone marrow-derived macrophages (BMDMs), peritoneal macrophages, liver, and fat **(Figure 1A)**.

**Figure 1:**
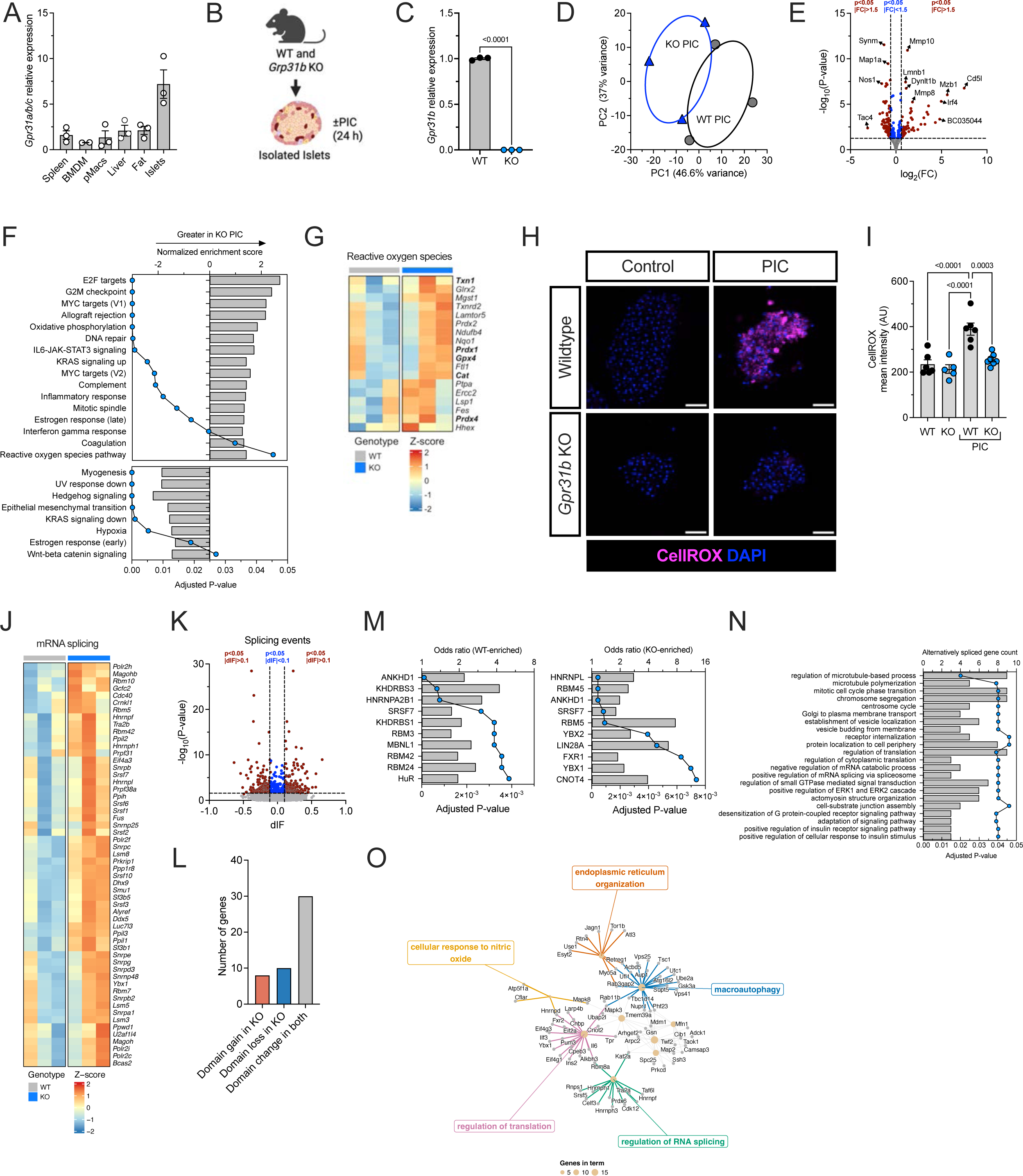
GPR31 regulates mRNA transcription and splicing of genes involved in β-cell stress and responses to reactive oxygen species. **(A)** Relative expression of Gpr31b in tissues from C57BL/6J mice by quantitative PCR. BMDM: bone marrow-derived macrophages; pMacs: peritoneal macrophages. **(B)** Schematic of experimental approach for RNA sequencing of islets treated with vehicle or pro-inflammatory cytokines (PIC). **(C)** RNA sequencing data showing relative expression of Gpr31b in WT and KO islets treated with PIC. **(D)** Principal components analysis (PCA) of WT and Gpr31b KO islets treated with PIC. **(E)** Volcano plot of differentially expressed genes in WT and Gpr31b KO islets treated with PIC. **(F)** Gene set enrichment analysis (GSEA) of WT and Gpr31b KO islets treated with PIC using hallmark gene sets. Both activated (top panel) and suppressed gene sets (bottom panel) are shown. All pathways meeting significance criteria of adjusted P-value <0.05 are shown. **(G)** Heatmap from WT and Gpr31b KO islets treated with PIC showing Z-scores for genes in the leading edge for the ROS hallmark gene set. **(H)** Representative fluorescence images of WT and Gpr31b KO islets treated with vehicle or PIC. Imaging shows nuclei stained by 4’,6-diamidino-2-phenylindole (DAPI; blue) and CellROX (magenta). Scale bars are 50 µm. **(I)** Quantification of CellROX mean intensity. **(J)** Heatmap from WT and Gpr31b KO islets treated with PIC showing Z-scores for genes in the leading edge for the mRNA splicing GO gene set. **(K)** Volcano plot of differentially spliced genes in WT and Gpr31b KO islets treated with PIC. **(L)** Quantification of the number of isoform switches resulting in protein domain gain or loss. **(M)** Motif enrichment analysis of the switching isoforms. **(N)** GO overrepresentation of alternatively spliced genes; alternatively spliced gene count (bars) and adjusted P-value (blue dots). **(O)** GO overrepresentation network of spliced genes (gray dots) and GO terms (gold dots) for which size represents genes per term. Data presented as mean with SEM. Statistical significance determined by Student’s t-test or one-way analysis of variance (ANOVA). Single data points represent individual mice unless otherwise noted.

To study the role of GPR31 in islet inflammation, we generated a whole-body knockout (KO) of *Gpr31b* on the *C57BL/6J* background. Because of the extensive homology between *Gpr31b* and its pseudogenes, we generated conditional *Gpr31b^Loxp/+^* mice using long-arm bacterial artificial chromosome (BAC) technology. Chimeric mice were then crossed to *Sox2*-Cre mice to generate somatic germline whole-body *Gpr31b*^-/-^ (herein referred to as *Gpr31b* KO) mice **(Figure S1A)** and backcrossed more than 10 generations onto the *C57BL/6J* background. WT and *Gpr31b* KO mouse genotypes were confirmed by PCR genotyping **(Figure S1B)**. To verify the deletion of *Gpr31b* in the *Gpr31b* KO mice, we performed quantitative PCR for *Gpr31a/b/c* mRNA in isolated pancreatic islets. *Gpr31a/b/c* mRNA expression was reduced by 40% in *Gpr31b* KO islets compared to islets isolated from WT littermates **(Figure S1C)**. Because TaqMan-based quantitative PCR analysis does not distinguish between *Gpr31b* and its two pseudogenes, we sequenced the PCR products to identify each gene individually. We found that WT islets express 70% *Gpr31a* and 30% *Gpr31b*, while *Gpr31b* KO islets only express *Gpr31a*, confirming that the appropriate gene has been deleted **(Figure S1D)**. *Gpr31b* KO mice exhibit normal Mendelian inheritance and appear healthy. Both male and female KO mice have a comparable body weight to WT littermates and have normal glucose homeostasis at 8 and 15 weeks of age **(Figure S1E-J)**. These data are similar to *Alox15^-/-^* mice, which also appear normal and healthy with normal glucose homeostasis (25).

### GPR31 mediates inflammatory and reactive oxygen species signaling in islets

Because 12-LOX inhibition reduces pro-inflammatory islet responses (26,27), we next wanted to investigate how GPR31 affects the transcriptome of pancreatic islets under normal and pro-inflammatory conditions. Therefore, we isolated islets from WT and *Gpr31b* KO mice treated with vehicle or pro-inflammatory cytokines (PIC; TNF-α, IFN-γ, and IL-1β), extracted RNA, and performed RNA sequencing **(Figure 1B)**. Of note, *Gpr31b* expression in this RNA sequencing analysis was only identified in WT islets treated with PIC **(Figure 1C)**. Principal components analysis (PCA) revealed two distinct clusters between vehicle-treated WT and *Gpr31b* KO islets **(Figure S2A)**. When comparing vehicle-treated islets from WT and *Gpr31b* KO mice, 64 out of 15,212 identified genes reached the differential expression thresholds of absolute fold change to ≥1.5 and adjusted P-value <0.05 **(Figure S2B and Data S1)**. Using GSEA with hallmark gene sets, 16 pathways were significantly enriched (adjusted P-value <0.05). Hallmark gene sets with increased enrichment notably included oxidative phosphorylation, whereas gene sets with decreased enrichment included epithelial-mesenchymal transition (EMT), proliferation (G2M checkpoint), protein secretion, inflammatory response, and TGFβ signaling **(Figure S2C)**.

Upon PIC treatment, PCA revealed two distinct clusters **(Figure 1D)**. When comparing PIC-treated islets from WT and *Gpr31b* KO mice, 150 genes out of 15,813 identified reached the differential expression thresholds of absolute fold change ≥1.5 and adjusted P-value <0.05 **(Figure 1E and Data S1)**. The top gene upregulated in the *Gpr31b* KO islets was *Cd5l*, which encodes the scavenger receptor CD5L with known roles in anti-inflammatory responses and lipid metabolism (28,29). Given that CD5L is a secreted protein, we confirmed by enzyme-linked immunosorbent assay (ELISA) that CD5L protein levels are greater in the supernatant of *Gpr31b* KO islets treated with PIC compared to WT islets **(Figure S2D)**. Using GSEA with hallmark gene sets to analyze PIC-treated islets, 24 pathways were significantly enriched (adjusted P-value <0.05) **(Figure 1F)**. Hallmark gene sets with increased enrichment included transcription (E2F targets), proliferation (G2M checkpoint), oxidative phosphorylation, inflammatory responses, and reactive oxygen species (ROS), whereas EMT was decreased, consistent with the findings for the vehicle-treated KO islets **(Figure 1F)**. We visualized a subset of the differentially expressed genes in this data set meeting more stringent criteria (absolute fold change ≥2 and adjusted P-value <0.05) **(Figure S2E)**. These differentially expressed genes included those related to proliferation (*Lmnb1*, *Kif2c*, *Hmmr*), immunity (*Cd5l*, *Irf4*, *Mzb1*), and neuronal identity (*Nos1*, *Phox2b*, *Adcyap1*). For a more granular analysis of pathway enrichment, we performed GSEA with gene ontology (GO) annotations, which showed suppression of multiple pathways related to neurotransmitter or synapse function, as well as activation in pathways related to immunity **(Figure S2F)**. Using the pathlinkR package (30), we visualized groups of ≥5 Reactome gene sets, which revealed clusters related to insulin secretion and metabolic signaling, neuronal signaling and synaptic transmission, and cell cycle DNA replication **(Figure S2G)**, consistent with our analysis of differential gene expression. Given the known importance of ROS in islet inflammation, we examined expression of the gene transcripts found in the leading edge of the hallmark ROS gene set, which showed upregulation of multiple antioxidant genes including catalase (*Cat*), glutathione peroxidase 4 (*Gpx4*), peroxide scavengers (*Prdx1*, *Prdx4*), and protein disulfide reduction (*Txn1*) **(Figure 1G)**.

Given this enrichment of transcripts encoding antioxidant genes, we next directly asked whether GPR31 regulates cytokine-induced oxidative stress in islets. To answer this question with a functional readout, we used a fluorescent dye-based free radical sensor (CellROX). We treated WT islets with PIC for 18 h which resulted in an increase in ROS production **(Figure 1H-I)**. By contrast, PIC-induced ROS production was dramatically reduced in *Gpr31b* KO islets, consistent with the results of our RNA sequencing and what we have previously observed with *Alox15^-/-^* mice or inhibition of 12-LOX (20,26).

### GPR31 regulates islet mRNA splicing of genes involved in cell stress and responses to nitric oxide

Given the prominent regulation of transcriptional processes in the *Gpr31b* KO islets **(Figure S2G)** and importance of mRNA splicing in cytokine-induced β-cell stress (31,32), we next examined mRNA splicing in our RNA sequencing data. Reactome GSEA of our sequencing data showed an enrichment of transcripts in mRNA splicing in islets **(Data S2)**. Visualization of the leading edge genes in the GO gene set for mRNA splicing, which showed increases in key categories of RNA splicing proteins including heterogenous ribonuclear proteins and small nuclear ribonucleoproteins **(Figure 1J)**. Using DEXSeq and the isoformSwitchAnalyzeR packages (33,34), we performed analysis of differential isoform fraction (dIF) in the *Gpr31b* KO islet RNA sequencing data. There were no significant changes in specific types of alternative splicing with *Gpr31b* KO **(Figure S2H)**. However, 380 isoforms from 253 genes out of 15,136 isoforms tested met both significance cutoffs of adjusted P-value <0.05 and dIF >0.1 **(Figure 1K)**. These splicing events resulted in multiple functional differences in transcript coding, topology, subcellular location, open reading frame length or gain/loss, nonsense mediated decay, and intron retention **(Figure S2I)**. Importantly, in the *Gpr31b* KO islets, there were 8 isoform switches resulting in protein domain gain and 10 isoform switches resulting in protein domain loss **(Figure 1L and Figure S2J)**. Motif enrichment analysis of the switching isoforms identified overrepresentation of RNA-binding protein motifs relative to non-switching isoforms, with KHDRBS3 and HNRNPA2B1 ranked highest among isoforms preferred in WT islets and hnRNP L and RBM45 among those preferred in *Gpr31b* KO islets **(Figure 1M)**. Overrepresentation analysis showed select GO terms associated with the significantly spliced genes demonstrated a network centered around the cellular response to nitric oxide (NOS), regulation of translation and ER organization, autophagy, and RNA splicing **(Figure 1N-O)**. Examining specific splice events revealed isoform switches concentrated in the ER, secretory, and autophagy pathways highlighted by the GO network. *Gpr31b* KO islets favored longer coding isoforms of the ER and secretory regulators *Tor1b*, *Fkbp2*, *Jagn1*, and *Stxbp2* and of the autophagy and endolysosomal regulators *Tbc1d14* and *Vps25*, alongside a switch in *Mdm2*, the E3 ubiquitin ligase that targets p53 for proteasomal degradation **(Figure S2K-L)**. These data demonstrate that *Gpr31b* KO in islets alters mRNA splicing of transcripts related to response to NOS, β-cell stress pathways, and RNA splicing, contributing to protection from islet inflammation.

### Gpr31b KO mice are protected against multiple low-dose streptozotocin-induced diabetes

To test whether *Gpr31b* KO mice are resistant to cytokine-induced oxidative stress and β-cell dysfunction *in vivo*, we subjected male WT and *Gpr31b* KO mice to multiple low-dose streptozotocin (mldSTZ) injections (55 mg/kg daily for 5 days) **(Figure 2A)**. In this model, STZ selectively damages β cells, leading to the recruitment of macrophages and dendritic cells into the vicinity of the islets and the local release of pro-inflammatory cytokines. This leads to oxidative stress, islet dysfunction, and eventual β-cell death (35–38). In response to mldSTZ injections, WT mice exhibited worsening blood glucose levels as early as day 5 of injections, whereas, through day 12, *Gpr31b* KO mice remained euglycemic **(Figure 2B)**. Likewise, *Gpr31b* KO mice had improved glucose tolerance compared to WT controls at day 10 **(Figure 2C)**. Additionally, β-cell mass was significantly higher in *Gpr31b* KO mice than WT controls **(Figure 2D-E)**. Additionally, *Gpr31b* KO mice had fewer macrophages within their islets compared to WT mice, though this did not meet statistical significance **(Figure 2D and F)**. We next examined both oxidative stress levels and macrophage recruitment into the islet based on our previous results. WT mice exhibited increased levels of 4-HNE (a marker of oxidative stress) while islets of *Gpr31b* KO mice had reduced levels of 4-HNE (**Figure 2G-H**). These data are consistent with prior studies showing that global and β-cell–specific *Alox15^-/-^* mice are protected from mldSTZ-induced diabetes (20,39).

**Figure 2:**
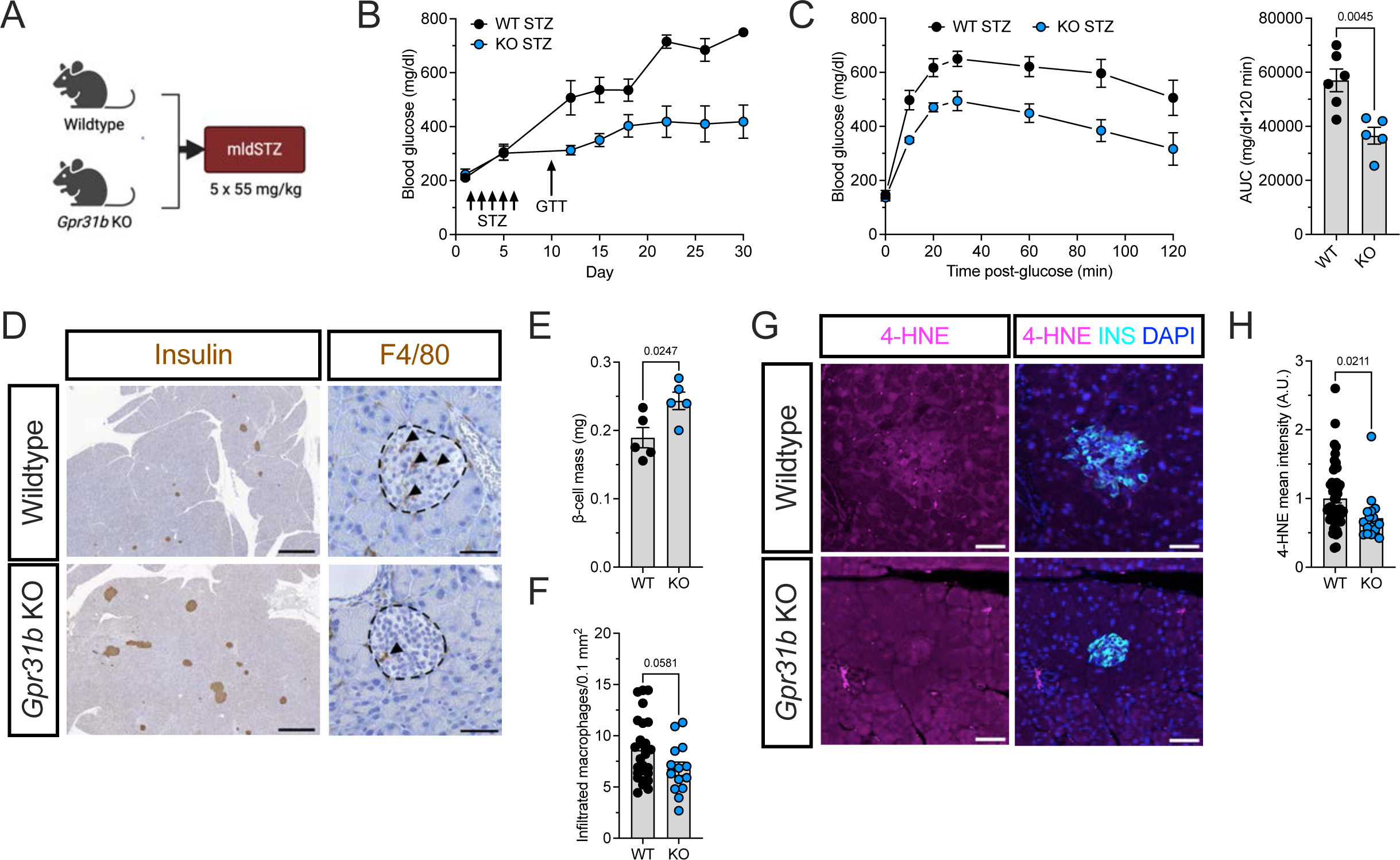
Gpr31b KO mice are protected against multiple low-dose streptozotocin (mldSTZ)-induced diabetes. (**A)** Schematic of mldSTZ experimental design. **(B)** Blood glucose measurements for WT and *Gpr31b* KO mice after mldSTZ administration (n=5–6 mice per condition). **(C)** GTT at 10 days after start of mldSTZ administration (*left panel*) with AUC (*right panel*). **(D)** Representative images of pancreata immunostained for insulin (left panel; scale bars = 500 µm) and macrophage marker F4/80 (right panel; scale bars = 150 µm). Islets are marked with a dotted line and infiltrated (intra-islet) macrophages are marked with arrowheads). **(E)** Quantification of β-cell mass. **(F)** Quantification of infiltrated (intra-islet) macrophages per area (n=14–28 islets per condition). **(G)** Representative immunofluorescence labeling for nuclei (DAPI; blue), insulin (teal), and the oxidative stress marker 4-HNE (magenta). Scale bars = 50 µm. **(H)** Quantification of 4-HNE mean intensity (n=17–51 islets per condition). Data presented as mean with SEM. Statistical significance determined by Student’s t-test or one-way analysis of variance (ANOVA).

### siRNA knockdown of Gpr31b reduces insulitis, macrophage migration, and oxidative stress in NOD mice

We next sought to study the role of GPR31 in the context of a T1D mouse model. To do this, we utilized small interfering RNA (siRNA) against *Gpr31b* (si*Gpr31b*) and a non-targeting control pool siRNA (siNTC or NTC). We first confirmed that si*Gpr31b* produced the expected knockdown of *Gpr31b* in β cells. In MIN6 murine β cells, we stimulated *Gpr31b* expression using PIC and found that *Gpr31a/b/c* levels are reduced upon siRNA knockdown **(Figure 3A)** to the approximate predicted levels based on gene expression of the *Gpr31b* pseudogenes, which should not be altered **(Figure S1C-D)**. We administered siRNA by intraperitoneal (IP) injection into prediabetic 6-week-old female NOD mice, a timeframe characterized by immune invasion during which the islets exhibit ER and oxidative stress (2). Mice received three injections of 1.7 mg/kg each of either si*Gpr31b* or siNTC spaced one day apart. The study ended 2 weeks after the initial dose of siRNA at which point we harvested pancreatic tissue and performed flow cytometry on pancreatic lymph nodes (pLN) **(Figure 3B)**.

**Figure 3:**
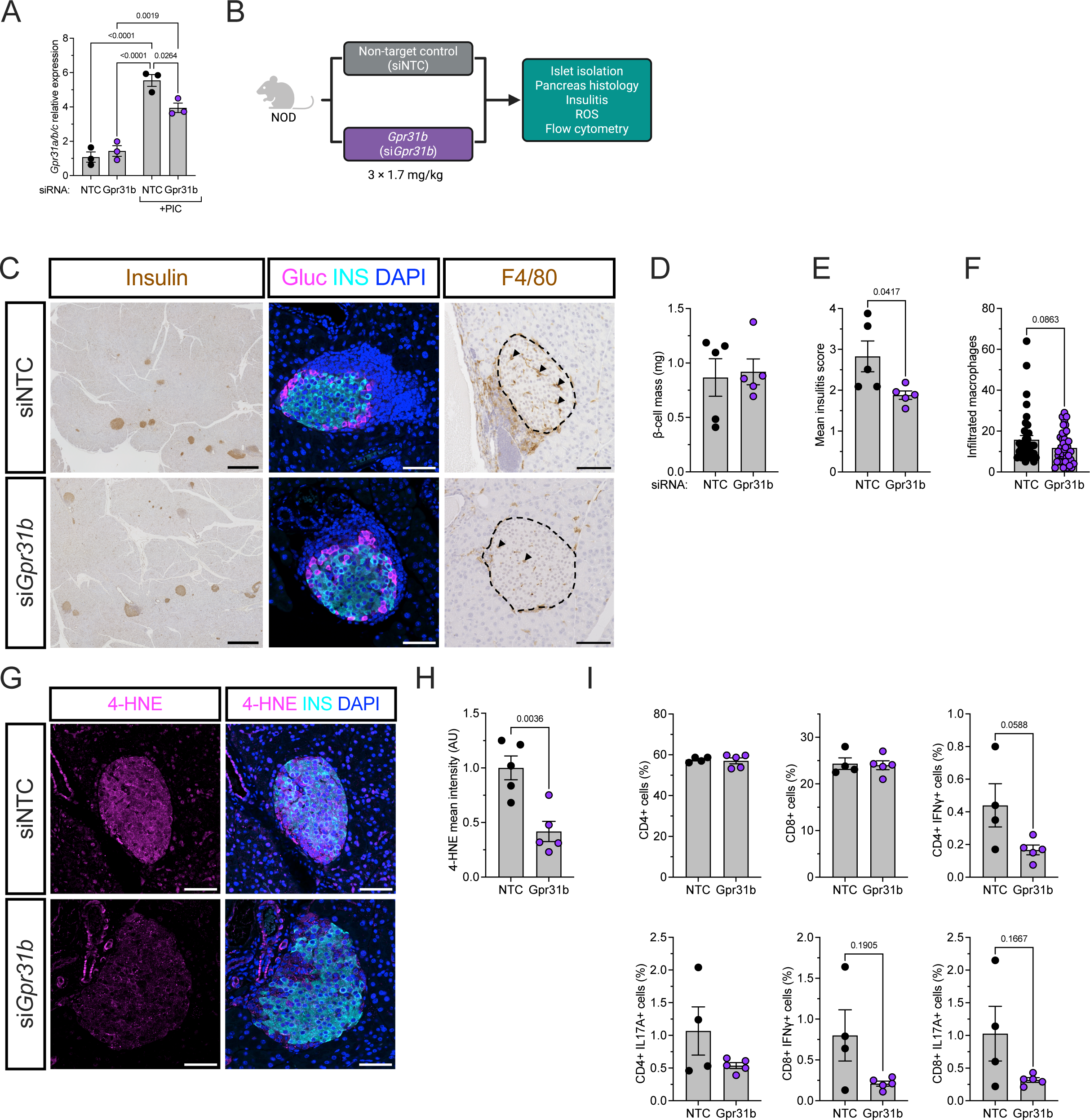
Small interfering RNA (siRNA) knockdown of Gpr31b reduces insulitis, macrophage migration, and oxidative stress in NOD mice. **(A)** Relative Gpr31a/b/c mRNA expression in MIN6 murine β cells treated with vehicle or PIC as well as non-targeting control pool siRNA (siNTC or NTC) and siRNA against Gpr31b (siGpr31b). **(B)** Schematic of siRNA injection experiment in female prediabetic NOD mice. **(C)** Representative images of pancreata immunostained by IHC for insulin (left panel; scale bars = 500 µm); by immunofluorescence for insulin (cyan), glucagon (magenta), and nuclei (DAPI; blue) (middle panel; scale bars = 100 µm); or by IHC for the macrophage marker F4/80 (right panel; scale bars = 200 µm). Islets are marked with a dotted line and infiltrated (intra-islet) macrophages are marked with arrowheads. **(D)** Quantification of β-cell mass. **(E)** Mean insulitis score. **(F)** Quantification of infiltrated (intra-islet) macrophages per area (n=39–42 islets per condition). **(G)** Representative immunofluorescence labeling in pancreatic tissue showing nuclei (DAPI; blue), insulin (green), and the oxidative marker 4-HNE (magenta). Scale bars = 100 µm; islets are marked with a dotted line. **(H)** Quantification of 4-HNE mean intensity. **(I)** Flow cytometry of CD4^+^ or CD8^+^ T cells from pancreatic lymph nodes (pLN). Data presented as mean with SEM. Statistical significance determined by Student’s t-test. Single data points represent individual mice.

We first examined the islets by immunohistochemistry and found that si*Gpr31b* did not alter the islet architecture or β-cell mass **(Figure 3C-D)**. However, the amount of insulitis was reduced in the NOD mice treated with si*Gpr31b* compared to control-injected mice **(Figure 3C and E)**. We next analyzed macrophage migration by immunohistochemical staining for F4/80 in the pancreas. We observed a reduced number of macrophages in the islet area of NOD mice treated with si*Gpr31b* compared to control-injected mice **(Figure 3C and F)**, although these data were not statistically significant. Additionally, these islets exhibited less oxidative stress as shown by 4-HNE staining **(Figure 3G-H)**. Lastly, to assess for an immune cell phenotype in lymphocytes with *Gpr31b* knockdown, we performed flow cytometry from both splenic tissue and pLN. We labeled for CD19 (B cell), CD4 and CD8 (T cells), FoxP3 (T regulatory cells), IFN-γ (Th1 cells), and IL17A (Th17 cells). Flow cytometry of splenic tissue showed no differences between siNTC and si*Gpr31b* injected mice **(Figure S3A)**. In contrast, flow cytometry of pLN showed a trend toward decreased IFN-γ+ and IL-17A+ T cells; however, this did not meet statistical significance **(Figure 3I and Figure S3B)**. Collectively, these findings demonstrate that GPR31 promotes insulitis, macrophage infiltration, and oxidative stress in prediabetic NOD mice.

### GPR31 shapes the islet inflammatory microenvironment

Using G4451, a recently described small molecule inhibitor of GPR31 (40), we designed an experiment to ask whether pharmacologic inhibition of GPR31 impacts disease severity in NOD mice at a prediabetic stage **(Figure S4A)**. NOD mice were treated with 60 mg/kg/day of G4451 by oral gavage or vehicle (carboxymethylcellulose-Na [CMC-Na]) from 6 weeks to 10 weeks of age. Pancreatic tissues from G4451-treated mice at 10 weeks of age showed no differences in mean insulitis score, β-cell mass, or infiltrating peri-islet macrophages compared to vehicle-treated mice **(Figure S4B–D)**. However, there was decreased 4-HNE fluorescence intensity in G4451-treated mice compared to vehicle-treated mice, consistent with both the mldSTZ-treated *Gpr31b* KO mice and si*Gpr31b* mice **(Figure S4E–F)**.

We next performed bulk RNA sequencing of islets isolated from both NOD treatment groups: 1) si*Gpr31b*– and NTC-injected mice; and 2) G4451– and vehicle-treated mice. Samples from si*Gpr31b*-injected mice were clearly separated from NTC control by PCA **(Figure 4A)**. Similarly, there was clear separation between G4451– and vehicle-treated islets **(Figure 4A)**. When comparing si*Gpr31b* islets to NTC control, 41 genes out of 22,924 identified genes reached the differential expression threshold of absolute fold change ≥1.5 and adjusted P-value <0.05 **(Figure 4B and D; Data S3)**. When comparing G4451-treated islets to vehicle-treated mice, 286 genes out of 24,148 identified reached the differential expression thresholds of absolute fold change ≥1.5 and adjusted P-value <0.05 **(Figure 4B and 4F, Data S4)**. For si*Gpr31b*, GSEA and pathlinkR network analysis of groups with ≥5 Reactome gene sets showed clusters representing increased cytokine-receptor and immune signaling, cell cycle, DNA replication, GPCR and chemokine signaling, and extracellular matrix related gene sets, among others **(Figure 4C and Figure S5A–B)**. For G4451, pathlinkR network analysis of networks with ≥5 Reactome gene sets showed enrichment of innate immune and cytokine-receptor signaling, translation and viral RNA processing, IFN signaling, antigen presentation, DNA replication, and cell cycle, among others **(Figure 4C and Figure S5C–D)**. Collectively, these data demonstrate that GPR31 knockdown or inhibition results in changes in the inflammatory microenvironment.

**Figure 4:**
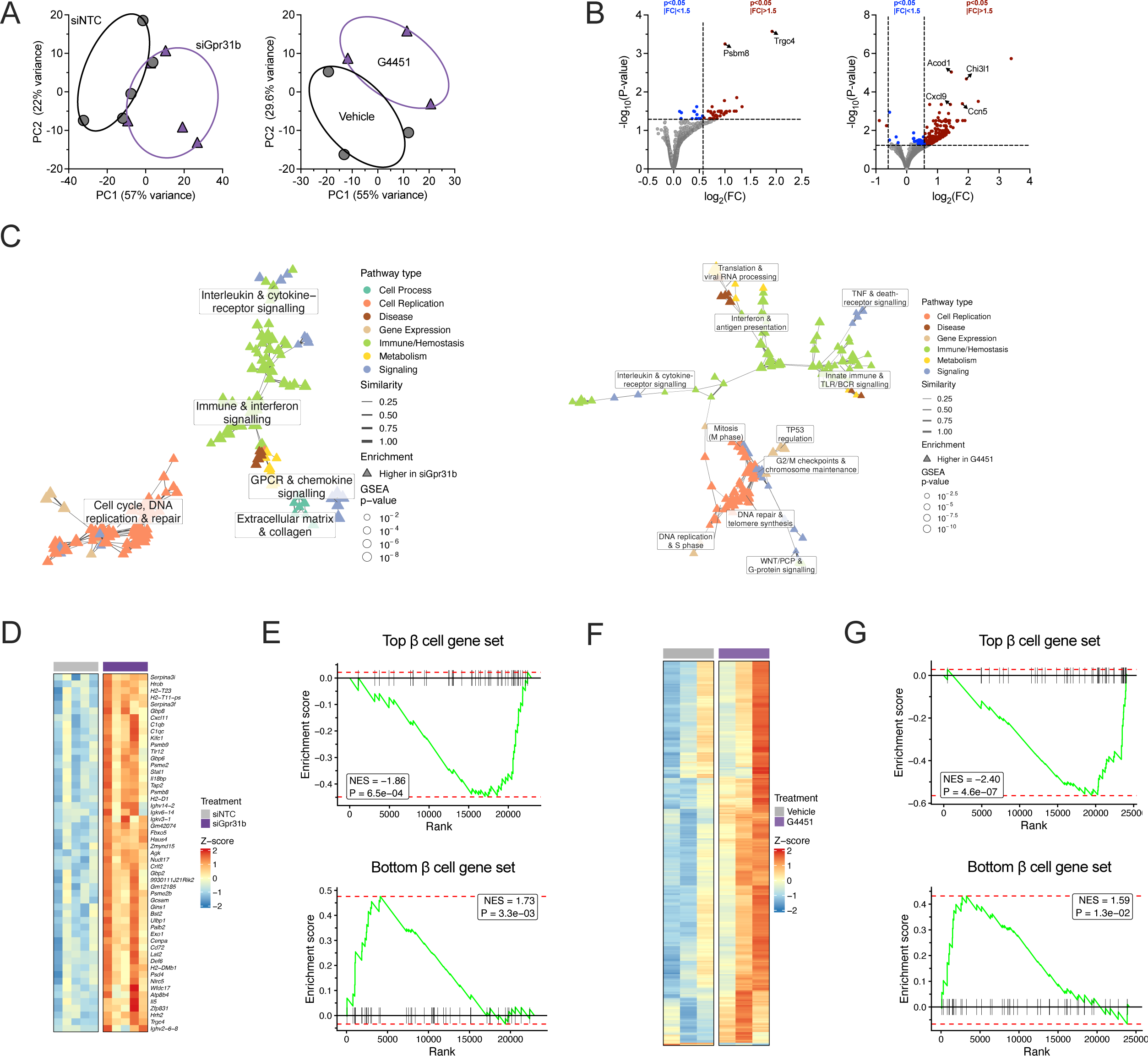
GPR31 knockdown or inhibition shapes the islet inflammatory microenvironment and shifts islets toward a bottom β cell phenotype. Islets from NOD mice injected with siNTC or siGpr31b (n=5 per group) and from NOD mice treated with vehicle or G4451 (n=3 per group). **(A)** PCA, siRNA (left) and G4451 (right). **(B)** Volcanoes, siRNA (left) and G4451 (right). **(C)** pathlinkR Reactome networks, siRNA (left) and G4451 (right). **(D)** Differentially expressed genes, siRNA. **(E)** GSEA of top (upper) and bottom (lower) β-cell signatures, siRNA; P-values are medians across random seeds. **(F)** Differentially expressed genes, G4451. **(G)** GSEA of top (upper) and bottom (lower) β-cell signatures, G4451. Data presented as mean with SEM. Statistical significance determined by Student’s t-test. Single data points represent individual mice.

### Loss of GPR31 results in an increase in “bottom” β cells and an islet-protective immune microenvironment

Given the findings of *Gpr31b* KO in the mldSTZ model and si*Gpr31b*-injected or G4451-treated NOD mice, we sought to address the apparent contradiction between increased islet inflammation and improvement in diabetes severity. Prior work has established a population of β cells that have deficits in insulin secretory capacity and increased PD-L1 expression and markers of stemness (41). These “bottom” β cells are relatively resistant to immunologic attack compared to more functional, “top” β cells. Using publicly available RNA-seq data (GSE94382) (41), we performed reanalysis of flow-sorted top versus bottom β cells, demonstrating separation by PCA **(Figure S6A)** and 1,315 DEGs **(Figure S6B-C and Data S5)**. GSEA of Hallmark and Reactome gene sets showed the expected findings, namely decreased β-cell identity, protein secretion and increased inflammatory signaling, including IFN-α and IFN-γ, in the bottom β cells **(Figure S6D-E)**. Using Boruta random forest machine learning, we established two separate gene signatures which reliably distinguished between top and bottom β cells **(Figure S6F)**. Application of the top and bottom β-cell gene signatures using GSEA showed decreased enrichment of the top gene signature and increased enrichment of the bottom gene signature in both the si*Gpr31b*-treated islets and G4451-treated islets **(Figure 4E and G)**.

Finally, given that siRNA and small molecule inhibitors have off-target effects, we performed a concordance analysis of the islet RNA sequencing from the bottom β cells, the G4451-treated, and the si*Gpr31b*-injected mice using comparison of the normalized enrichment scores from GSEA. Comparison of gene sets significantly enriched in at least one of the three data sets showed a core group of gene sets related to inflammation, including IFN-α and IFN-γ, TNFA signaling via NFKB, and others **(Figure 5A and Figure S7)**. To identify gene expression changes shared between the two islet RNA sequencing data sets, we performed a meta-analysis across experiments. For each gene detected in both data sets, the DESeq2 Wald statistics from the G4451 and si*Gpr31b* experiments were combined using Stouffer’s method and the resulting P-values were corrected for multiple testing by the Benjamini–Hochberg procedure. By this analysis, 1,645 genes were significantly upregulated and 418 genes were significantly downregulated across both the G4451 and si*Gpr31b* RNA sequencing experiments **(Figure 5B)**. Among the genes that were increased were *Acod1*, which encodes the enzyme cis-aconitate decarboxylase, and *Cd274*, encoding PD-L1, a key marker of bottom β cells and known to be increased in islets from donors with T1D (42). Immunolabeling of pancreatic tissues from G4451-treated mice showed increased ACOD1 and PD-L1 without changes in MHC-I **(Figure 5C–H)**. Examination of PanKbase data demonstrated that *ACOD1* expression is lost in β cells from donors with T1D whereas *CD274* is increased, as expected **(Figure 5I–J)**. These data demonstrate that loss of GPR31 results in an islet-protective immune microenvironment, potentially mediated in part by increased expression of PD-L1 and ACOD1.

**Figure 5:**
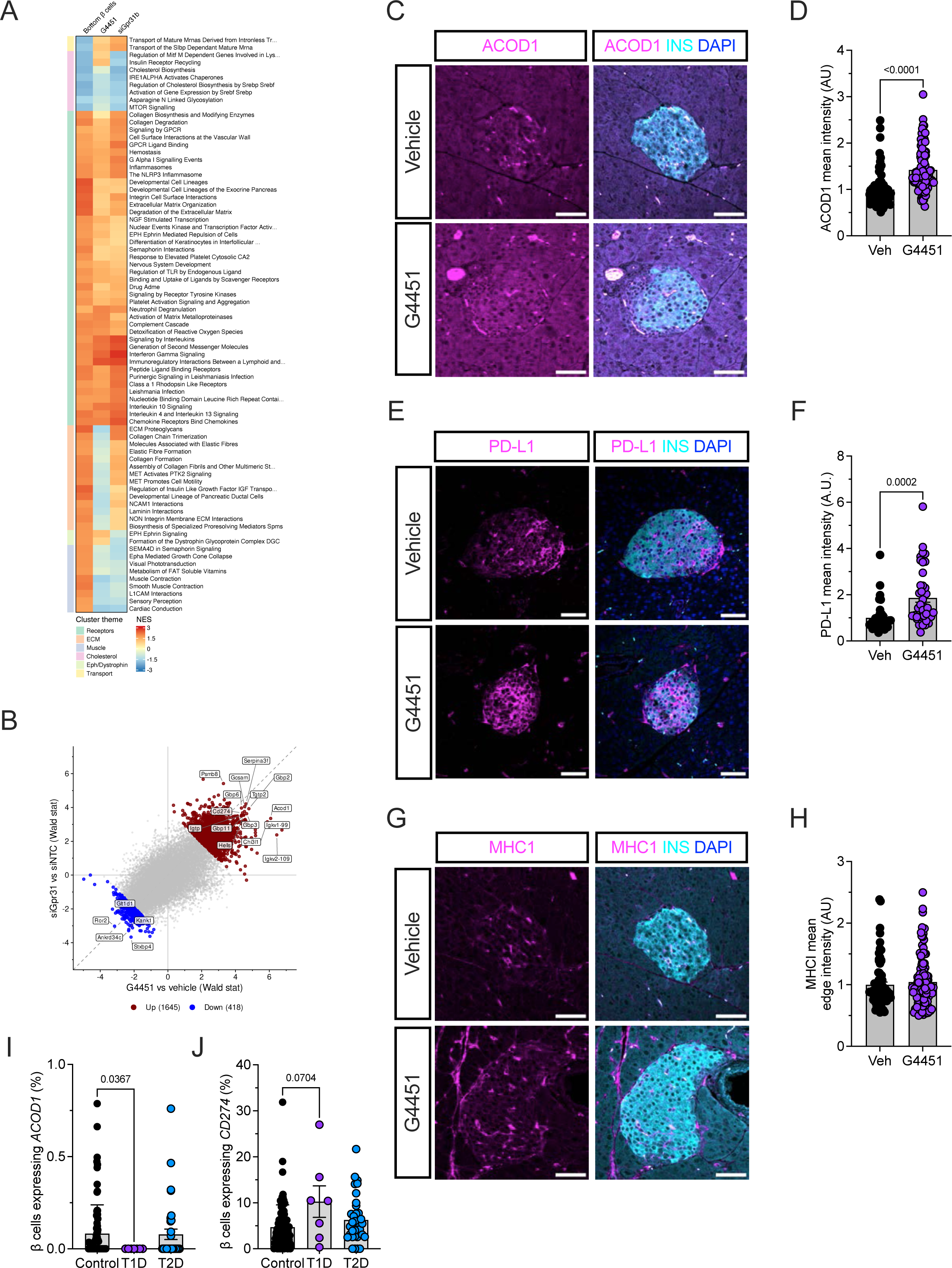
GPR31 knockdown and inhibition converge on an islet-protective immune signature marked by ACOD1 and PD-L1. (**A)** Normalized enrichment scores for Reactome gene sets from RNA-sequencing of sorted bottom β cells, si*Gpr31b*, and G4451. **(B)** DESeq2 Wald statistics for G4451 versus vehicle and si*Gpr31b* versus siNTC combined by Stouffer’s method with Benjamini–Hochberg correction. **(C-H)** Pancreata from vehicle– and G4451-treated NOD mice stained for ACOD1 **(C)**, PD-L1 **(E)**, or MHC class I **(G)** in magenta with insulin (cyan) and DAPI (blue), with quantification **(D, F, H)**. Scale bars, 50 µm. n=37–104 islets per condition. **(I, J)** β cells expressing *ACOD1* **(I)** or *CD274* **(J)** in PanKbase human islet donors without diabetes (n=86), with type 1 diabetes (n=7), or with type 2 diabetes (n=34–35). Data presented as mean with SEM. Statistical significance determined by Student’s t-test or Kruskal–Wallis test.

### GPR31 is expressed in pro-inflammatory macrophages

Given our data showing decreased macrophage infiltration in both *Gpr31b* KO mice treated with mldSTZ **(Figure 2F)** and NOD mice injected with si*Gpr31b* **(Figure 3F)**, we next asked whether GPR31 was expressed in macrophages. Using publicly available RNA sequencing datasets of mouse immune cells and intra-islet macrophages (23,43), we examined the distribution of *Gpr31b* expression across immune cell populations. Analysis of these datasets showed that intra-islet macrophages and other myeloid cell lineages including monocytes and classical dendritic cells had the highest expression of *Gpr31b* **(Figure 6A)**. To further investigate the *Gpr31b* expression of intra-islet macrophages, we used prior, pre-annotated single-cell RNA sequencing data from non-obese diabetic (NOD) mice at multiple ages (44). These data demonstrated that islet macrophages had the greatest number of *Gpr31b^+^*cells **(Figure 6B-C)**. Using existing subclustering of these data (44), we found relative enrichment of *Gpr31b*^+^ cells in a subcluster of resident macrophages within the islet **(Figure S8A–C)**. Gene set enrichment analysis (GSEA) with Reactome gene sets of *Gpr31b*^+^ macrophages showed enrichment in major histocompatibility complex (MHC) I and II, antigen processing and presentation, interferon (IFN) signaling, mRNA splicing, and G*_⍺_*_i_ signaling events **(Figure 6D)**. GSEA with hallmark gene sets similarly showed enrichment of the IFN-α and IFN-γ response **(Figure S8D)**. Collectively, these data show expression of *Gpr31b* in pancreatic islets and intra-islet macrophages, and that *Gpr31b*^+^ macrophages mount a stronger inflammatory response through antigen processing, IFN signaling, and related pathways.

**Figure 6:**
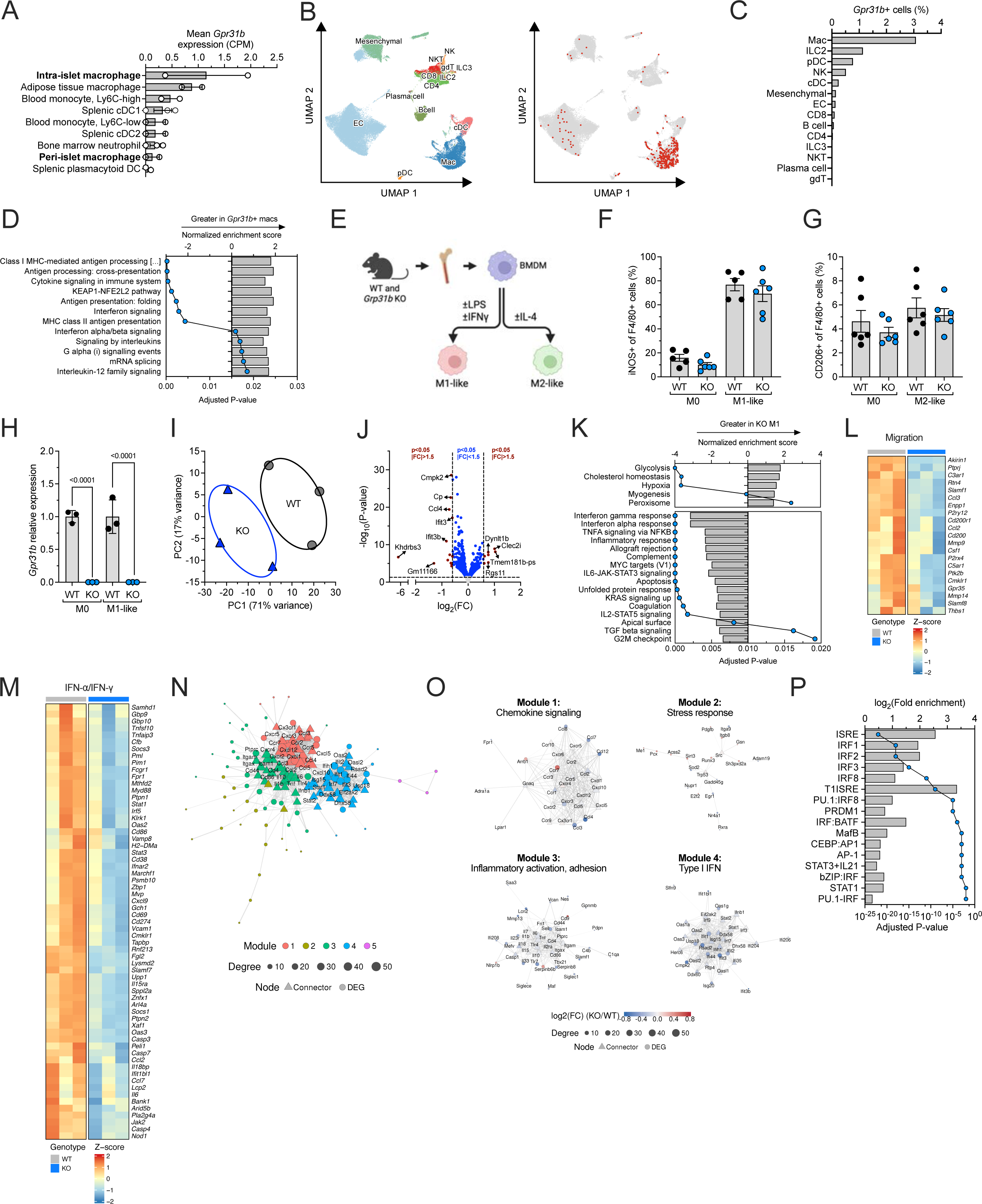
GPR31 is expressed in pro-inflammatory macrophages and drives interferon signaling. (**A)** Relative expression of *Gpr31b* by RNA sequencing from publicly available RNA sequencing datasets of mouse immune cells and intra-islet macrophages. **(B)** UMAP of pre-annotated single-cell RNA sequencing data from NOD mouse islets showing cell type annotation (*left panel*) and *Gpr31b*^+^ cells in red (*right panel*). **(C)** Quantification of *Gpr31b+* cell types. **(D)** GSEA from Reactome gene sets of *Gpr31b*^+^ macrophages. **(E)** Schematic of experimental approach for polarization of macrophages. **(F)** Flow cytometry for the M1 marker iNOS as a percentage of F4/80+ cells in M0 and M1-polarized macrophages. **(G)** Flow cytometry for the M2 marker CD206 as a percentage of F4/80+ cells in M0 and M2-polarized macrophages. **(H)** Relative expression of *Gpr31b* by RNA sequencing. **(I)** PCA of M1-polarized macrophages. **(J)** Volcano plot of M1-like macrophages. **(K)** GSEA of M1-polarized macrophages using hallmark gene sets. All significant pathways are shown. **(L)** Leading edge genes, GO cell migration gene set. **(M)** Leading-edge genes, hallmark IFN-α and IFN-γ response. **(N)** STRINGdb network of differentially expressed genes colored by module and **(O)** individual modules colored by log_2_ fold change, KO versus WT; node size, degree; triangles, connector nodes. **(P)** HOMER promoter motif enrichment for genes downregulated in *Gpr31b* KO; bars, log_2_ fold enrichment. ISRE, interferon-stimulated response element. Data presented as mean with SEM. Statistical significance determined by Student’s t-test or one-way analysis of variance (ANOVA).

### GPR31 promotes macrophage inflammatory signaling

We next sought to establish the function of GPR31 in macrophages. To determine if GPR31 affects macrophage polarization, we isolated peritoneal macrophages from WT and *Gpr31b* KO mice. We polarized these macrophages into classically activated (M1-like) macrophages using LPS and IFN-γ or alternatively activated (M2-like) macrophages using IL-4 and initially analyzed their phenotypes by flow cytometry **(Figure 6E)**. The macrophages from both WT and *Gpr31b* KO mice showed an indistinguishable propensity to polarize to an M1-like state, as assessed by flow cytometry of the M1 marker inducible nitric oxide synthase (iNOS) **(Figure 6F)**. Similarly, upon alternative polarization, macrophages from both WT and *Gpr31b* KO mice showed no differences by flow cytometry in the M2 marker CD206 **(Figure 6G)**.

We next performed RNA sequencing on both M0 and M1-like BMDMs isolated as previously described (3) and polarized as above. Normalized counts from RNA sequencing confirmed *Gpr31b* KO in both M0 and M1-like macrophages **(Figure 6H)**. PCA revealed two distinct clusters between M0 macrophages from WT and *Gpr31b* KO mice **(Figure S9A)**. When comparing M0 macrophages from WT and *Gpr31b* KO mice, 71 genes out of a total 13,882 identified genes reaching the differential expression thresholds of absolute fold change ≥1.5 and adjusted P-value <0.05 **(Figure S9B and Data S5)**. PCA revealed two distinct clusters between M1-like macrophages from WT and *Gpr31b* KO mice **(Figure 6I)**. When comparing M1-like macrophages from WT and *Gpr31b* KO mice, only 17 genes out of a total 12,914 identified genes identified genes reached the differential expression thresholds of absolute fold change ≥1.5 and adjusted P-value <0.05 **(Figure 6J and Data S5)**. Using GSEA with hallmark gene sets revealed decreased enrichment of multiple inflammatory signaling pathways in the *Gpr31b* KO macrophages, including IFN-γ and IFN-*α* responses, as well as increased enrichment of metabolic pathways including glycolysis and cholesterol homeostasis **(Figure 6K)**. Given the known roles of 12-LOX in migration, we used the leading edge genes from the GO migration gene set and visualized expression by heatmap, which showed consistent downregulation of genes in the leading edge **(Figure 6L)**. Similarly, using the leading edge genes from the Hallmark IFN-α and IFN-γ gene sets, we visualized expression by heatmap, which showed consistent downregulation of genes in each leading edge, including genes shared between both pathways **(Figure 6M)**, with the full leading edge of each interferon gene set shown separately **(Figure S9C)**. We similarly visualized the leading edge genes from the Hallmark glycolysis and fatty acid metabolism gene sets, which showed coordinated changes in metabolic gene expression in the Gpr31b KO macrophages **(Figure S9D)**. PathlinkR network analysis with ≥5 Reactome gene sets per cluster showed decreased enrichment of IFN signaling, antigen presentation, GPCR processes, and mRNA splicing, among others **(Figure S9E)**. Complementary analysis using Reactome gene sets showed 77 significant pathways, including decreased enrichment of GPCR ligand binding, signaling by GPCR, G*_α_*_1_ signaling events, antigen processing and cross presentation, fatty acid metabolism, and carbohydrate metabolism **(Data S6)**. Using the protein-protein interaction database STRINGdb (45), we analyzed the DEGs and identified modules that differentiated WT from *Gpr31b* KO macrophages, including chemokine signaling, stress response, inflammatory activation and adhesion, and type I IFN signaling **(Figure 6N-O)**. HOMER analysis showed enrichment of multiple motifs involved in IFN signaling, including the interferon-stimulated response element (ISRE) **(Figure 6P and Figure S9F)**. Collectively, these data show that GPR31 drives pro-inflammatory signaling in M1-polarized macrophages through IFN and alters the transcriptional machinery of mRNA splicing and antigen processing.

### GPR31 alters macrophage mRNA splicing in genes related to GPCR signaling and actomyosin contractility

Given the decreased enrichment of mRNA splicing genes in the M1-like *Gpr31b* KO macrophages **(Data S7)**, we next investigated whether there were changes in alternative splicing due to *Gpr31b* KO in BMDMs. Using the same bulk RNA sequencing data set, we visualized the leading edge genes in the GO gene set for mRNA splicing, which showed decreases in key mRNA splicing genes including the heterogenous ribonuclear proteins *Hnrnph2* and *Hnrnpa2b1* as well as splicing activators and couplers *Snw1*, *Fus*, and *U2af1* **(Figure 7A)**. Using DEXSeq and the isoformSwitchAnalyzeR packages, we performed analysis of differential isoform usage (dIF) in the *Gpr31b* KO RNA sequencing data. There were no significant changes in specific types of alternative splicing with *Gpr31b* KO **(Figure 7B)**. However, 111 isoforms in 86 genes out of 15,426 isoforms tested met both significance cutoffs of adjusted P-value <0.05 and dIF >0.1 **(Figure 7C)**. These splicing events resulted in multiple functional differences in transcript coding, topology, subcellular location, open reading frame length or gain/loss, nonsense mediated decay, and intron retention **(Figure 7D)**. Importantly, in the *Gpr31b* KO macrophages, there were 94 isoform switches resulting in protein domain gain and 87 isoform switches resulting in protein domain loss **(Figure 7E and Figure S10A)**. Using analysis of motif enrichment, there was overrepresentation of several key motifs related to transcriptional regulation **(Figure 7F)**. Overrepresentation analysis of the GO terms associated with the significantly spliced genes demonstrated a network centered around GPCR signaling and cell migration **(Figure 7G and S10B)**. Examining specific splice events in genes revealed key changes related to the macrophage phenotype. Several spliced genes related to microtubules and actomyosin contractility (*Rapgef2* and *Camsap2*), whereas others were key members of canonical GPCR signal transduction pathways (*Arrb1* and *Gnai2*) **(Figure S10C–D)**. These data demonstrate that *Gpr31b* KO in macrophages alters mRNA splicing of transcripts related to GPCR signaling and actomyosin contractility, contributing to the phenotype of dampened IFN signaling and macrophage migration.

**Figure 7:**
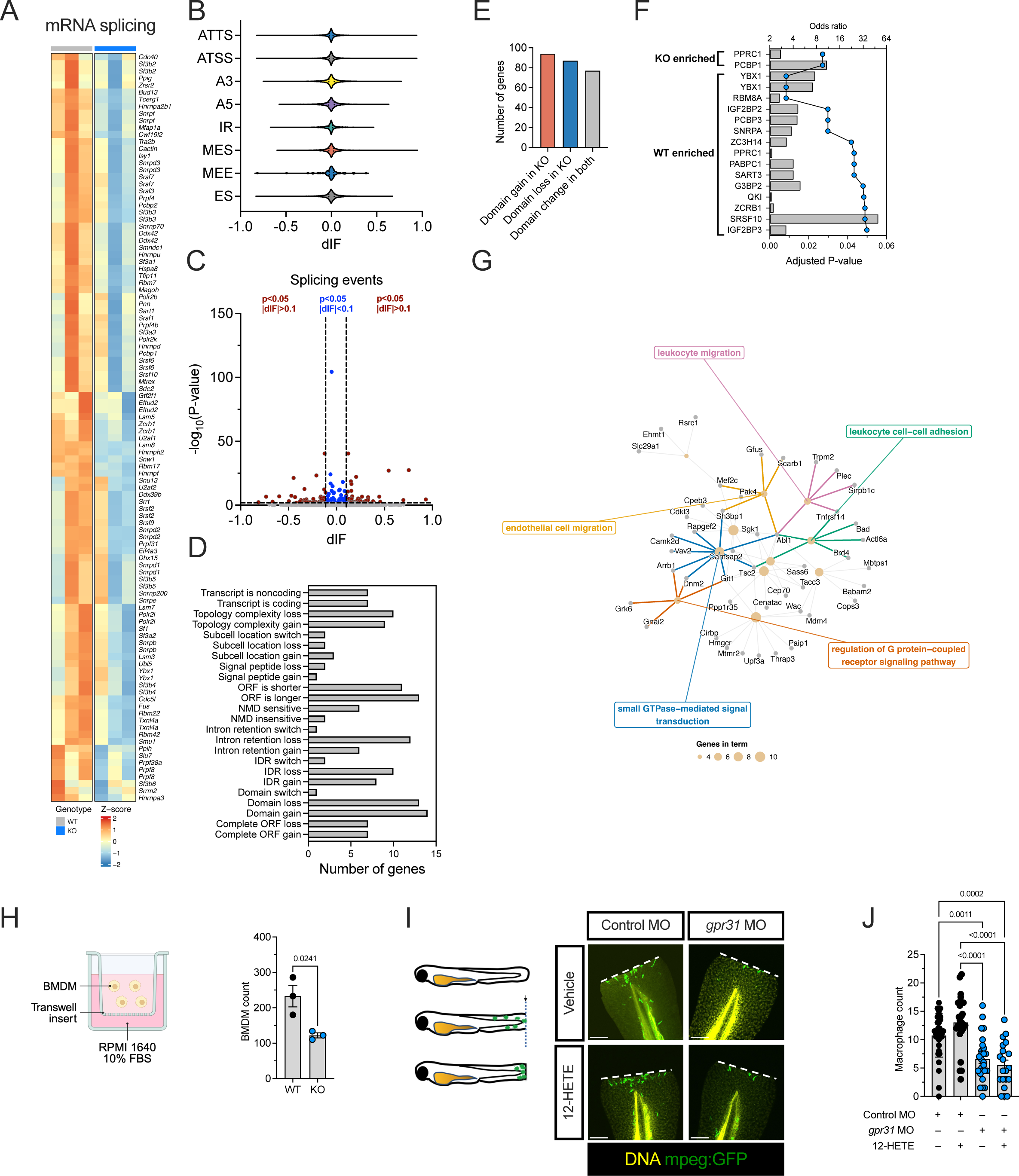
GPR31 regulates splicing of inflammatory genes and promotes macrophage migration. (**A)** Leading-edge genes for RNA splicing from M1-polarized macrophages. **(B)** Quantification of types of RNA splicing events. **(C)** Volcano plot. **(D)** Functional consequences of isoform switches. **(E)** Genes with protein domain gain, loss, or domain changes in both WT and KO. **(F)** RNA-binding protein motif enrichment in isoforms preferred in WT and in *Gpr31b* KO; odds ratio (bars) and adjusted P-value (dots). **(G)** GO overrepresentation network of spliced genes (gray dots) and GO terms (gold dots) for which size represents genes per term. **(H)** Schematic (*left panel*) and quantification (*right panel*) of BMDM transwell migration assay. **(I)** Schematic (*left panel*) and representative images (*right panel*) of zebrafish tailfin cut assay showing DNA (TO-PRO; yellow) and macrophages (GFP; green). Scale bars = 100 µm. **(J)** Quantification of assay results as macrophage count. Data presented as mean with SEM. Statistical significance determined by Student’s t-test or one-way ANOVA.

### GPR31 promotes macrophage migration

To determine if GPR31 affects cell migration, another major macrophage function, we isolated BMDMs from WT and *Gpr31b* KO mice and performed an *in vitro* migration assay using transwell chambers **(Figure 7H)**. The number of *Gpr31b* KO macrophages that migrated through the transwell membrane was significantly fewer than the number of WT macrophages **(Figure 7H)**. To verify these results and quantify macrophage migration *in vivo*, we used the tailfin transection model in *Tg(mpeg:eGFP)* transgenic zebrafish in which macrophages are fluorescently labeled with green fluorescent protein (GFP) **(Figure 7I)**. We performed knockdown of the zebrafish ortholog, *gpr31*, using a morpholino (MO) as previously described (24). Both *gpr31* MO knockdown and control zebrafish underwent tailfin transection at 3 days post fertilization (dpf), and the number of macrophages migrating to the injury site after 6 h was quantified **(Figure 7I–J)**. In control zebrafish, we observed the expected migration of macrophages (10.7±3.8 macrophages) to the injury site, whereas *gpr31* MO zebrafish showed a significant reduction in macrophage recruitment (6.60±3.95 macrophages) **(Figure 7I–J)**. Given that the pro-inflammatory eicosanoid 12-HETE is a ligand for GPR31 (21), we also tested whether exogenous 12-HETE could rescue the migration defect by including 12-HETE treatment (3 µM) or vehicle in the same experiment. Treatment with 12-HETE did not significantly alter migration in either control MO (13.0±4.7 macrophages) or *gpr31* MO injected zebrafish (5.5±3.94 macrophages), failing to rescue the knockdown phenotype. In contrast, zebrafish injected with an MO against the 12-LOX zebrafish ortholog *alox12* showed a similar reduction in migration, but this defect was rescued by the addition of 12-HETE **(Supplemental Figure S10E)**. These results are consistent with GPR31 functioning downstream of 12-HETE. Taken together, these findings indicate that the 12-HETE–GPR31 axis is required for macrophage migration but not for cytokine-induced polarization of macrophages.

### GPR31 is expressed in dendritic cells in human pancreatic lymph node

Finally, we used an existing cellular indexing of transcriptomes and epitopes by sequencing (CITE-seq) dataset of the spleen, pancreatic lymph node, mesenteric lymph nodes, and superior mesenteric artery lymph nodes from donors without diabetes, with autoantibody positivity, and with T1D (46). Reanalysis of these data demonstrated that cDC1 cells expressed *GPR31* and were found primarily in the spleen and to a lesser degree in the pancreatic lymph node **(Figure S11A–B)**. GPR31 expression in cDC1 was approximately 3-fold higher in AAb+ or T1D donors than in non-diabetic donors, although this did not reach significance given the limited number of donors **(Figure S11C)**. Like murine *Gpr31b^+^* macrophages, these *GPR31*^+^ cDC1 cells are enriched in the inflammatory response, IFN responses, and GPCR signaling and show altered mRNA splicing and antigen presentation **(Figure S11D–E)**.

## DISCUSSION

We identify the orphan G protein-coupled receptor GPR31 as a key mediator of pancreatic islet inflammation and autoimmune diabetes development. Our findings place GPR31 at the intersection of β-cell stress and macrophage activation, paralleling but also extending the established role of 12/15-LOX and its lipid metabolite 12-HETE in T1D pathogenesis. Specifically, our data show that (i) genetic deletion, knockdown, or inhibition of GPR31 reduces islet oxidative stress, macrophage inflammatory signaling, and macrophage infiltration of the islet; (ii) loss of GPR31 alters mRNA splicing in both β cells and macrophages; and (iii) loss of GPR31 shifts islets toward a bottom-like β-cell state. We discuss here the role of GPR31 in the 12-LOX–12-HETE axis, cell type-specific roles in β cells and macrophages, and potential consequences of GPR31 expression in human myeloid cells.

12-LOX and 12-HETE are established mediators of T1D pathogenesis in both β cells and macrophages. 12-HETE impairs human β-cell function, and reduction of 12-HETE with 12-LOX inhibition protects human islets from cytokine-induced injury (16,27). Whole-body 12-LOX deletion protects from STZ-induced and autoimmune diabetes (39,47,48), as does conditional knockout in either β cells or the myeloid lineage (4,6). These converging roles underscore the centrality of the 12-LOX pathway in β-cell dysfunction and macrophage activation. Given that GPR31 is a high-affinity receptor for 12-HETE (21), we asked whether GPR31 deficiency would recapitulate the protective effects of 12-LOX loss. Although 12-LOX is most abundant in macrophages (17), we found GPR31 expressed in pancreatic islets. Whole-body *Gpr31b* KO mice, like whole-body 12-LOX KO mice (39), were protected from hyperglycemia and loss of β-cell mass after STZ-induced diabetes. Our data showed that *Gpr31b* KO protected islets from cytokine-induced injury, most notably by reducing cytokine-induced oxidative stress. β cells are highly susceptible to oxidative damage, which is amplified by cytokines in T1D. Because 12-LOX activity promotes oxidative stress in β cells and its inhibition increases antioxidant defenses (17,20,26), our findings support a model in which blockade of 12-HETE signaling via GPR31 deletion reduces oxidative stress and thereby sustains β-cell health.

Cytokine-treated *Gpr31b* KO islets also showed loss of synaptic and neurotransmitter-related genes. Because β cells express a subset of neuronal genes and share developmental machinery (49–51), we interpret this signature as loss of mature β-cell identity rather than evidence of a neural GPR31 pool, consistent with the shift toward a low-identity, immune-resistant “bottom” β-cell state described below and potentially protective under the pro-inflammatory conditions of T1D (41,52). Islets from si*Gpr31b*-injected and G4451-treated NOD mice showed increased inflammatory gene set enrichment despite reduced insulitis and oxidative stress. Bottom β cells are a subpopulation with low secretory output, reduced autoantigen expression, and increased surface PD-L1 that resist immune– and cytokine-mediated killing (41), and similar cells may be enriched among surviving β cells in human T1D (53). Both knockdown and inhibition shifted islets toward this state across two unrelated perturbations, and we read the inflammatory signature as accompanying a β-cell state that trades maturity for immune resistance. One key molecular feature of this program may be PD-L1 expression on β cells. PD-L1 is IFN-inducible through IRF1 (42), rises with T cell infiltration in NOD islets and delays diabetes when overexpressed in β cells (54,55). Notably, β-cell–specific *Alox15* deletion on the NOD background raises β-cell PD-L1 and anti-PD-L1 reverses the resulting protection (4), placing this checkpoint downstream of the same 12-LOX axis in which GPR31 may operate. ACOD1, induced alongside CD274, is a separate candidate whose itaconate product restrains cytokine production by innate immune cells (56,57), though endogenous ACOD1 has not been examined systematically in T1D. That β-cell ACOD1 falls in human T1D while CD274 rises may reflect loss of an early protective program or selection among surviving β cells.

A surprising consequence of GPR31 loss was the perturbation of mRNA splicing. Cytokines remodel splicing networks in primary β cells and human islets (31,32). For example, cytokine-associated loss of spliceosomal regulators shifts β cells toward apoptosis-linked isoforms (58). Splice variants also supply epitopes for autoreactive CD4 T cells (59,60). Our findings implicate GPR31 as a key regulator of alternative splicing in islets and macrophages. *Gpr31b* KO redirected isoform usage in genes relating to ER and secretory function, autophagy, and the cellular response to nitric oxide in islets, and in genes relating to GPCR signal transduction and motility in macrophages. Our motif analysis indicated a shift in the RNA-binding proteins governing splice-site selection but the specific mechanism by which the GPCR activity of GPR31 relates to alternative splicing remains unresolved. Like many GPCRs, GPR31 recruits β-arrestin (61), which has been recently shown to form condensates that compartmentalize GPCR signaling and may lend selectivity to its regulation of mRNA splicing (62). Resolving these questions will require careful GPCR-focused experiments, including ligand-stimulated and rescue controls, GPR31-versus β-arrestin–focused perturbations, and better resolution of resulting splice isoforms using long-read sequencing.

GPR31 also exerts distinct effects in macrophages. Both si*Gpr31b*-injected NOD mice and STZ-treated *Gpr31b* KO mice showed fewer intra-islet macrophages. These findings mirror the phenotype of whole-body and myeloid 12-LOX knockout mice, which also show reduced islet immune-cell invasion and protection from diabetes (6,47,48). Islet macrophages play a pivotal role in initiating and amplifying the immune response in early T1D, in part through local secretion of cytokines including TNF-α and IL-1β, and through presentation of β-cell antigen to infiltrating lymphocytes (63,64). GPR31 loss impaired macrophage migration in both BMDMs and the zebrafish tailfin transection assay. Migration loss in *gpr31*-deficient zebrafish was not rescued by 12-HETE, whereas migration in alox12-deficient zebrafish was. In macrophages, the function of GPR31 appears to drive inflammation, in particular through IFN-*α* and IFN-γ signaling. This suggests an early role in T1D, as circulating IFN-α precedes autoantibody conversion (65–67). IFN-α is thought to derive from innate immune cells, and signaling downstream of IFN-α drives β-cell apoptosis, ER stress, and ROS production (68–70). These data also raise the possibility that the islet-specific effects of *Gpr31b* KO are in part secondary to activated IFN-α signaling by GPR31 in macrophages. Our transcriptomic data also showed decreased pathway enrichment of antigen presentation in M1-like *Gpr31b* KO macrophages. Together, these findings place GPR31 downstream of 12-HETE in the regulation of macrophage motility and indicate that it sustains an IFN-driven, antigen-presenting program, both of which may contribute to early islet infiltration.

In human lymphoid tissue, *GPR31* expression was largely restricted to cDC1 cells. These *GPR31*^+^ cDC1 cells appear to carry part of the transcriptional programs of murine *Gpr31b*^+^ macrophages, including enriched IFN and antigen presentation signatures. In NOD mice, resident macrophages sample β-cell granules and present antigen locally (7,71), whereas BATF3-dependent cDC1s are required for cross-presentation to autoreactive CD8 T cells and for progression to destructive insulitis (72,73). In humans, imaging of pancreata identifies cDC-like cells near islets peaking at onset of stage 3 T1D (74). Notably, GPR31 is selectively expressed on human intestinal cDC1s, where the pyruvate–GPR31 axis promotes transepithelial dendrite formation (75), whereas the analogous mouse function was shown in CX3CR1^+^ myeloid cells (76). These findings suggest that GPR31 supports antigen presentation through different myeloid populations across species. Whether GPR31 expressed in human pancreatic cDC1s contributes to islet autoimmunity remains to be established.

This study has several limitations. Conditional models targeting β cells and myeloid lineages are needed to define cell-specific function of GPR31. The STZ model engages innate rather than adaptive immunity, and additional models will be required to test broader relevance in T1D. Further pharmacologic work with G4451 and other GPR31 inhibitors (40,61) will be required to define the full impact of GPR31 blockade in T1D. Our mRNA splicing findings are computational and not orthogonally validated. Finally, *GPR31*^+^ cell numbers in the reanalyzed mouse and human single-cell RNA sequencing datasets were too low for comprehensive comparison across disease states.

In summary, our data establish GPR31 as a critical inflammatory mediator in the pathogenesis of T1D. By linking 12-LOX–12-HETE signaling to β-cell oxidative stress, macrophage migration and IFN responses, and the splicing programs underlying both, GPR31 integrates cell type-specific mechanisms that converge on the islet microenvironment. Cell type-specific approaches will be required to resolve how GPR31 signaling in β cells and in myeloid cells each contributes to the early inflammatory events of T1D.

## RESEARCH DESIGN AND METHODS

### Mouse studies

All animal experiments were approved by the University of Chicago Institutional Animal Care and Use Committee. All mice used in this study were housed under specific pathogen-free conditions with a standard 12 h light:dark cycle. *Gpr31b*^loxP/+^ mice were generated using conditional long-arm bacterial artificial chromosome (BAC) technology to insert loxP sites around the single exon 1 and a Neo cassette for selection with flippase recombinase (FLP) sites for removal (Ingenious Targeting Labs, Ronkonkoma, NY). The Neo cassette was deleted via FLP recombination in vitro using FLP-expressing embryonic stem cells. Chimeric mice were then crossed to *Sox2*-Cre mice to generate somatic germline whole-body *Gpr31b*^-/-^ (herein referred to as *Gpr31b* KO) mice **(Figure S1A)**. *Gpr31b* KO mice were backcrossed more than 10 generations onto the *C57BL/6J* background using remote speed congenics services (Jackson Labs, Bar Harbor, ME). Allele-specific genotyping was performed by polymerase chain reaction (PCR) using the following primers: 5’–GCA GGA CGT GTC TTG TAA ACA GC–3’ and 5’–TTC CCT GGG GAC TCT ATG GG–3’; 5’–CAA ATT CTG CTG GCT ACT TGG–3’ and 5’–GGC TTC CAT ACT TTG AGA CG–3’. Using these primers, the wildtype (WT) *Gpr31b* allele generates an amplicon of 335 bp and the mutant allele generates an amplicon of 699 bp. Additionally, there is a WT band that includes pseudogenes with an amplicon of 3,100 bp **(Figure S1B).**

For glucose tolerance testing at day 10 after the initial dose of STZ, mice were fasted for 16 h. Mice were IP injected with glucose (2 g/kg) and blood glucose was measured at 0, 10, 20, 30, 60, 90, and 120 min. Blood glucose was monitored via tail vein using a glucometer (AlphaTrak, Zoetis, Parsippany-Troy Hills, NJ). For multiple low-dose streptozotocin (mldSTZ) studies, mice were IP injected with 55 mg/kg STZ (Sigma, St. Louis, MO) for 5 consecutive days.

For small interfering RNA (siRNA) studies, 6-week-old female NOD mice were IP injected with 1.7 mg/kg of siRNA against a non-targeting control pool (siNTC) or *Gpr31b* (si*Gpr31b*) (Accell, Horizon Discovery, Cambridge, UK). Mice received three injections each spaced one day apart. The study ended 2 weeks after the initial dose of siRNA. For studies involving treatment with the GPR31 inhibitor G4451 (40), 6-week-old female NOD mice were administered either vehicle (0.5% CMC-Na in PBS) or 60 mg/kg of G4451 daily by oral gavage for 4 weeks.

At the end of each study, mice were euthanized and tissue and blood were collected. To isolate islets, collagenase was injected into the pancreatic bile duct to inflate the pancreas prior to removal as previously described (77). Briefly, a Histopaque–Hank’s balanced salt solution gradient was applied to the dissociated pancreas and followed by centrifugation at 900 rcf for 18 min. The mouse islets were then removed from the center of the gradient and cultured in RPMI 1640 medium supplemented with 10% fetal bovine serum (FBS) and 1% penicillin/streptomycin. Islets were handpicked and allowed to recover overnight before experimentation. Pro-inflammatory cytokines (PIC) were used at 10 ng/ml TNF-α, 100 ng/ml IFN-γ, and 5 ng/ml IL-1β throughout, for 24 h for RNA sequencing and 18 h for oxidative stress measurement.

### Zebrafish husbandry and maintenance

Zebrafish were maintained in 2 L tanks at 28.5°C in a recirculating aquaculture system, in accordance with University of Chicago Institutional Animal Care and Use Committee regulations. The transgenic line used for experiments was *Tg(mpeg:eGFP)^gl22^*(78). Embryos were collected at spawning and placed into egg water (0.1% Instant Ocean salts, 0.0075% calcium sulfate, and 0.1% methylene blue). At 24 h post-fertilization, the medium was replaced by egg water supplemented with 0.003% 1-phenyl-2-thiourea (PTU) to prevent pigmentation.

Morpholinos (MO) were purchased from GeneTools (Philomath, OR) with the following sequences: *gpr31* MO sequence: 5’–gtgca gattc ccagt ccgag atgac–3’; *alox12* MO sequence: 5’–ccact gtcac tttgt actcc atctt–3’ (24). For microinjections, MO were diluted with nuclease-free H_2_O and mixed with 0.1% phenol red marker. The injection volume was calibrated by determining the diameter of the injection droplet on mineral oil to inject 2 ng of the MO. Zygotes were collected at spawning and injected at the 1-cell stage. The embryos marked with phenol red were sorted for use in later experiments.

### Tailfin transection assay

For the tailfin transection assay, larvae at 3 days post-fertilization (dpf) were paralyzed with 0.01% tricaine to prevent movement. The distal tips of the tailfins were cut with a sharp scalpel as previously described (6). Following injury, embryos were placed in a 28.5°C incubator for 6 h with or without 12-HETE (3 µM; Cayman Chemicals, Ann Arbor, MI). At the end of the incubation period, the embryos were fixed with 3% formaldehyde in PEM buffer at 4°C for 18 h, washed, deyolked, and counterstained for nuclei with 4’,6-diamidino-2-phenylindole (DAPI; Thermo Fisher, Waltham, MA). The embryos were mounted on glass slides with antifade medium (Vector Labs, Newark, CA) and confocal imaging was performed with a Nikon A1 microscope (Nikon USA, Melville, NY). Macrophage number was determined by counting macrophages between the cut site and the distal tip of the notochord rod.

### Macrophage isolation and polarization

Bone marrow-derived macrophages (BMDMs) were isolated as described previously (6) and cultured for 7 days in complete medium: RPMI containing 10% FBS, 10 mM 4-(2-hydroxyethyl)-1-piperazineethanesulfonic acid (HEPES) buffer, 100 U/ml penicillin/streptomycin, and 10 ng/ml M-CSF. On day 7, BMDMs were stimulated for 16 h with 10 ng/ml lipopolysaccharide (LPS) and 25 ng/ml IFN-γ for M1-like polarization and 10 ng/ml IL-4 for M2-like polarization. Polarized BMDMs were subsequently utilized for *in vitro* migration studies and RNA sequencing.

Peritoneal macrophages were isolated from 8-10-week-old mice by injecting 10 ml ice cold phosphate-buffered saline (PBS) containing 3% FBS into the peritoneal cavity and removed using a 20-gauge needle as described previously (79). Cells were lysed with red blood cell (RBC) lysis buffer to remove RBCs and washed with PBS. Isolated peritoneal macrophages were used for flow cytometry.

### In vitro macrophage migration studies

For the migration assays, bone marrow-derived macrophages (2×10^5^) were placed in the upper chamber of an 8 µm transparent polyethylene terephthalate (PET) membrane (Corning, Corning, NY) and RPMI 1640 medium with 10% FBS was loaded in the bottom chamber. Cells were incubated for 4 h at 37°C, fixed in formalin and stained with crystal violet. Migrated cells were mounted on a slide and imaged using a BZ-X810 fluorescence microscope (Keyence, Elmwood, NJ). The number of macrophages was quantified by manual counting.

### Flow cytometry

To stain for surface antigens, peritoneal macrophages were incubated with F4/80 (BioLegend, San Diego, CA) and CD11b (BioLegend, San Diego, CA) antibodies for 20 min on ice. For intracellular staining, cells were permeabilized with Cytofix/Cytoperm (BD Pharmingen, Franklin Lakes, NJ) and incubated with CD206 (141706; BioLegend, San Diego, CA), iNOS (56-5920-82; Invitrogen, Carlsbad, CA), CD4 (100510; BioLegend), CD8 (100734; BioLegend), CD19 (152414; BioLegend), FoxP3 (560401, BD Biosciences), IFN-γ (554412, BD Biosciences), or IL17A (560821, BD Biosciences) antibodies for 20 min on ice. All antibodies were used at 1:100 dilution. Cells were analyzed on the Attune NxT Flow Cytometer (Thermo Fisher Scientific, Waltham, MA). Data were analyzed by FlowJo software (FlowJo, BD, Ashland, OR).

### Immunofluorescence and immunohistochemistry

Pancreata were fixed in 4% paraformaldehyde, paraffin embedded, and sectioned. Tissues were immunostained with anti-insulin (1:1000; 15848-1-AP, ProteinTech, Rosemont, IL) or anti-F4/80 (1:150; D2S9R, Cell Signaling, Danvers, MA) followed by conjugated anti-rabbit Ig (Vector Laboratories, Newark, CA), 3,3’-diaminobenzidine (DAB) peroxidase substrate kit (Vector Laboratories, Newark, CA), and counterstained with hematoxylin (Sigma, St. Louis, MO). Images were acquired using a BZ-X810 fluorescence microscope (Keyence, Elmwood, NJ). β-cell mass was calculated as previously described (80). For fluorescence labeling, tissues were immunostained with anti-4-HNE (1:200; ab65545, Abcam, Waltham, MA), anti-insulin (1:10, IR002, Dako, Carpinteria, CA), anti-glucagon (1:200, ab55092-100, Abcam, Waltham, MA), anti-ACOD1 (1:200, M13250, Boster Bio, Pleasanton, CA), anti-PD-L1 (1:200, AF1019, R&D Systems, Minneapolis, MN), or anti-MHC class I (1:100, 114602, BioLegend, San Diego, CA) followed by conjugated anti-rabbit Ig and anti-guinea pig Ig (1:500; Thermo Fisher, Waltham, MA) and DAPI. Imaging was performed using a Nikon A1 confocal microscope (Nikon, Melville, NY). Fluorescence intensity was quantified using Fiji (81) and CellProfiler (82). Intra-islet macrophages stained with F4/80 were counted manually.

We quantified β-cell mass and scored insulitis as previously described (35,83). Briefly, the insulitis score reflects the degree of immune cell infiltration within pancreatic islets. The scoring system we used is as follows: 1, no insulitis; 2, infiltrate <50% circumference; 3, infiltrate >50% circumference; 4, infiltration within islet.

### Enzyme-linked immunosorbent assay

CD5L protein levels in islet supernatants were quantified using a mouse CD5L enzyme-linked immunosorbent assay (ELISA) kit (orb565622; Biorbyt, Cambridge, UK) according to the manufacturer’s instructions. Briefly, islet culture supernatant was added to pre-coated wells and incubated for 90 min at 37°C, followed by sequential incubations with biotin-labeled detection antibody and horseradish peroxidase (HRP)-streptavidin conjugate. After incubation with 3,3′,5,5′-tetramethylbenzidine (TMB) substrate for no more than 30 minutes, optical density was measured at 450 nm and CD5L concentrations were calculated from a standard curve.

### Quantitative PCR

RNA was isolated from mouse tissues and macrophages using an RNeasy Mini kit (Qiagen, Germantown, MD) and cDNA synthesis was performed using a High-Capacity cDNA Reverse Transcription kit (Applied Biosystems, Waltham, MA) according to the manufacturer’s instructions. For siRNA studies, MIN6 murine β cells were cultured as previously described (84) and treated with and without pro-inflammatory cytokines (PIC; 10 ng/ml TNF-α, 100 ng/ml IFN-γ, 5 ng/ml IL-1β) and either 1 µM siNTC or si*Gpr31b* (Horizon Discovery, Cambridge, UK) for 24 h followed by RNA isolation. Quantitative PCR was performed using Bio-Rad CFX Opus with a predesigned Taqman assay probe for mouse genes *Gpr31b* (Mm02391728_g1) and *Actb* (Mm01205647_m1) (Invitrogen, Carlsbad, CA). Relative gene expression was calculated using the comparative threshold cycle value (C_t_) normalized to *Actb*.

### In vitro oxidative stress measurement

For assessment of oxidative stress in pancreatic islets isolated from WT and *Gpr31b* KO mice, we used a fluorescent dye-based free radical sensor (CellROX; Thermo Fisher Scientific, Waltham, MA). We followed the manufacturer’s instructions; briefly, islets were treated with PIC for 18 h. For the final 30 minutes of treatment with PIC, 5 µM CellROX reagent was added to the media. At endpoint, the islets were fixed and imaged as above. Intensity of CellROX was quantified using Fiji (81).

### Bioinformatics analyses

For bulk RNA sequencing, islets and BMDMs were isolated as described above. RNA was isolated using an RNeasy Mini kit (Qiagen, Germantown, MD). Reads were trimmed using trimmomatic (85) or cutadapt (86). Quality scores were assessed using FastQC. Reads were aligned to the *Mus musculus* genome build GRCm38 or GRCm39 using STAR (87). Individual sample reads were quantified using HTSeq (88) or featureCounts (89). Genes with fewer than 5 counts in fewer than 2 samples per group were filtered out. Raw counts were normalized via relative log expression with DESeq2 (90) using R Statistical Software (version 4.5–4.6; R Core Team 2025). Prior to differential expression analysis, batch effects between experimental replicates were corrected using ComBat-Seq (91). DESeq2 was used to calculate fold changes, adjust P-values for false discovery rate, and perform covariate correction. Gene set enrichment analysis was carried out using clusterProfiler (92) and the MSigDB hallmark database (93), Gene Ontology knowledgebase (94), or Reactome (95) as indicated. Reactome enrichment results were displayed as pathway networks with pathlinkR. Protein–protein interaction networks were built from STRING (45). Promoter motif enrichment was performed separately with HOMER (96) in gene-list mode against the mouse RefSeq promoter set.

Differential isoform usage was assessed with isoformSwitchAnalyzeR (33). Transcripts were quantified with StringTie in reference-guided mode, imported at the fragment level, filtered for expression, and tested with DEXSeq (34). Isoform switches were called at adjusted *P* <0.05 with absolute dIF >0.1, where dIF > 0 indicates greater use of an isoform in *Gpr31b* KO. Functional consequences of each switch were annotated from coding potential, protein domains, signal peptides, transmembrane topology, subcellular localization, and intrinsically disordered regions using CPC2, PfamScan, SignalP, DeepTMHMM, DeepLoc, and NetSurfP, respectively, and summarized with isoformSwitchAnalyzeR’s consequence analysis. Sequences of switching isoforms were scanned for RNA-binding protein motifs using universalmotif with one-sided Fisher testing against the Ray2013 collection (97). Enrichment against all non-switching isoforms was tested separately for isoforms preferred in *Gpr31b* KO and in WT.

This study reanalyzed the following publicly available datasets: GSE112002 (mouse islet-resident macrophage RNA sequencing) (23), GSE109125 (ImmGen ULI RNA sequencing of sorted mouse immune populations) (43), GSE107011 (RNA sequencing of sorted human immune cell types) (98), GSE141784 (single-cell RNA sequencing of NOD mouse islet-infiltrating immune cells) (44), GSE269540 (HPAP single-nucleus multiome of human pancreatic lymph node and spleen) (46), GSE221787 (HPAP CITE-seq of human pancreatic and non-pancreatic lymph nodes and spleen) (46), and PanKbase (99). For bulk data, differential expression used DESeq2 on counts (GSE112002, GSE109125, GSE94382) or limma-trend on log_2_(TPM+1) with donor as a blocking factor (GSE107011). To compare *Gpr31b* abundance across cell populations, mean expression in counts per million was computed for each sorted immune population and for intra– and peri-islet macrophages. GSE112002 was additionally re-quantified with salmon v1.10.2 against the GENCODE vM25 index. Single-cell data were processed in Seurat v5 with Harmony batch correction and depth-controlled (GSE141784 and GSE269540).

Top and bottom β cell signatures were derived from a published RNA-seq dataset (GSE94382) (41). Feature importance was computed with the Boruta algorithm in R (100), and all pre-filtered genes were ranked by mean importance and signed by the direction of their log_2_ fold change to yield “top-enriched” and “bottom-enriched” arms. The Boruta signature comprises the 100 highest-importance genes, which split 50/50 between arms. After gene-level filtering, 48 top-enriched and 47 bottom-enriched genes were retained, and the retained sets were identical across both test datasets.

For concordance analysis, results from the G4451 and si*Gpr31b* islet experiments were combined by Stouffer’s method, treating DESeq2 Wald statistics as Z scores with equal weights and correcting by the Benjamini–Hochberg procedure. Normalized enrichment scores were also compared directly across the bottom β cell, G4451, and si*Gpr31b* data sets, retaining gene sets significant in at least one and grouping them by theme for display. The two arms of the Boruta-derived top and bottom β cell signature were scored against the si*Gpr31b* and G4451 data by gene set enrichment analysis with fgsea (101) repeated across random seeds and reporting median P-values.

### Statistical analysis

Data analyses were performed using the GraphPad Prism 11 software package (GraphPad, Boston, MA). Unless otherwise noted, statistical testing to determine significant differences was performed using two-tailed, unpaired Student’s t-test for comparing 2 means and one-way ANOVA followed by post-hoc Tukey’s test for comparing ≥3 means. P-values that were <0.05 were considered statistically significant.

### Data and resource availability

RNA sequencing datasets have been deposited to the NIH Gene Expression Omnibus (GEO) with the accession numbers GSE284693 (*Gpr31b* KO islets), GSE285039 (*Gpr31b* KO macrophages), GSE349714 (siGpr31b islets), and GSE349713 (G4451 islets).

## ACKNOWLEDGMENTS

The authors wish to acknowledge Svetlana Navitskaya, Advaita Chakraborty, Kayla Figatner, Hailey Boyle, Maya Gauhar, and James Tersey at the University of Chicago for their technical assistance in these studies.

This work was supported in part by National Institutes of Health grants R03 TR003381 (SAT and RGM), R01 DK124906 (SAT), R01 DK105588 (RGM), R01 DK060581 (RGM), P30 DK020595 (Diabetes Research and Training Center grant to the University of Chicago), and K12 DK133995 (KBK; multi-center program directors David Maahs and Linda DiMeglio). Additional support was provided by the Pollak family through the Kovler Diabetes Center of the University of Chicago (KBK). The content is solely the responsibility of the authors and does not necessarily represent the official views of the National Institutes of Health.

This manuscript used data acquired from the database (https://hpap.pmacs.upenn.edu/) of the Human Pancreas Analysis Program (HPAP; RRID:SCR_016202) (102–105). HPAP is part of a Human Islet Research Network (RRID:SCR_014393) consortium (UC4 DK112217, U01 DK123594, UC4 DK112232, and U01 DK123716). Some experimental results were derived from human pancreatic islets or other resources provided by the NIDDK-funded Integrated Islet Distribution Program (IIDP) (RRID:SCR_014387) at City of Hope (UC4 DK098085).

## AUTHOR CONTRIBUTIONS

KBK, RGM, and SAT conceived and designed the research, provided funding, and wrote the manuscript. KBK, CC, JRE, TN, EE, AAP, JEW, MAW, CM, XSS, JBN, AK, SCM RMA, and SAT performed the research and acquired the data. All authors reviewed and edited the manuscript.

## Disclosure

The authors have nothing to disclose.

## SUPPLEMENTAL FIGURE LEGENDS

***Supplemental Figure S1: Mice with whole-body Gpr31b knockout (KO) exhibit normal body growth and glycemic control*. (A)** Schematic of approach to whole-body KO of *Gpr31b*. **(B)** Representative genotypes of wildtype (WT), heterozygous, and *Gpr31b* KO mice by PCR. The WT *Gpr31b* allele generates an amplicon of 335 bp and the mutant allele generates an amplicon of 699 bp. Additionally, there is a WT band that includes pseudogenes with an amplicon of 3,100 bp. **(C)** Relative expression of *Gpr31a*/*b*/*c* in WT, heterozygous, and *Gpr31b* KO mice by quantitative PCR. **(D)** Sequencing of PCR products of amplification of *Gpr31b* and its pseudogenes. **(E)** Body weight of 8-week-old males. **(F)** GTT of 8-week-old males (*left panel*) with area under the curve (AUC, *right panel*). **(G)** GTT of 15-week-old males (*left panel*) with AUC (right panel). **(H)** Body weight of 8-week-old females. **(I)** GTT of 8-week-old females (*left panel*) with AUC (*right panel*). **(J)** GTT of 15-week-old females (*left panel*) with AUC (*right panel*). Data presented as mean with SEM. Statistical significance determined by Student’s t-test or one-way analysis of variance (ANOVA).

***Supplemental Figure S2: Additional analyses of Gpr31b KO islet RNA sequencing.*** (**A**) PCA of WT and *Gpr31b* KO islets treated with vehicle. (**B**) Volcano plot of differentially expressed genes in islets treated with vehicle. **(C)** GSEA of islets treated with vehicle (n=3 mice per condition) using hallmark gene sets. All pathways meeting significance criteria of adjusted P-value <0.05 are shown. (**D**) CD5L concentration by enzyme-linked immunosorbent assay (ELISA) in the supernatant of islets treated with vehicle or PIC. **(E)** Heatmap from islets treated with PIC. **(F)** GSEA of islets treated with cytokines using GO annotations. **(G)** pathlinkR Reactome network. **(H)** Quantification of types of RNA splicing events. **(I)** Functional consequences of isoform switches. **(J)** Protein domains affected by alternative splicing per gene, for domain gain in KO (*left*), domain loss in KO (*middle*), and domains affected in both WT and KO (*right*). **(K)** Gene-level expression in transcripts per million (top) and isoform usage (bottom) for *Tor1b*, *Stxbp2*, *Fkbp2*, *Jagn1*, *Tbc1d14*, *Vps25*, and *Mdm2*. **(L)** Isoform structures for the genes in **(K)**, annotated for protein domains and predicted topology. Data presented as mean with SEM. Single data points represent individual mice. Statistical significance determined by Student’s t-test or one-way analysis of variance (ANOVA).

***Supplemental Figure S3: Flow cytometry from NOD mice after siRNA knockdown of Gpr31b.* (A)** Flow cytometry from splenic tissue of NOD mice injected with siNTC or si*Gpr31b*. **(B)** Flow cytometry from pancreatic lymph nodes (pLN) of NOD mice injected with siNTC or si*Gpr31b*.

***Supplemental Figure S4:* Inhibition of GPR31 reduces islet oxidative stress in NOD mice. (A)** Schematic of GPR31 inhibitor experiment in female prediabetic NOD mice. **(B)** Mean insulitis score. **(C)** Quantification of β-cell mass. **(D)** Quantification of infiltrated (intra-islet) macrophages per area. **(E)** Representative immunofluorescence labeling in pancreatic tissue showing nuclei (DAPI; blue), insulin (green), and the oxidative marker 4-HNE (magenta). Scale bars = 50 µm. **(F)** Quantification of 4-HNE mean intensity. Data presented as mean with SEM. Statistical significance determined by Student’s t-test.

***Supplemental Figure S5: GSEA from siRNA and inhibitor.* (A, B)** si*Gpr31b* versus siNTC islets, hallmark (**A**) and Reactome (**B**) gene sets (n=5 per group). **(C, D)** G4451-versus vehicle-treated islets, hallmark (**C**) and Reactome (**D**) gene sets (n=3 per group).

***Supplemental Figure S6: Segregation of top and bottom islet β cells.*** Public RNA-seq data (GSE94382) (n=4 per group). **(A)** PCA. **(B)** Volcano. **(C)** Differentially expressed genes. **(D)** hallmark GSEA. **(E)** Reactome GSEA. **(F)** Top-enriched (*upper*) and bottom-enriched (*lower*) gene signatures derived by Boruta random forest.

*Supplemental Figure S7: Concordance of hallmark gene set enrichment across bottom β cells, G4451-treated islets, and siGpr31b islets.* Normalized enrichment scores for hallmark gene sets significant in at least one of bottom β cells, G4451-treated islets, and si*Gpr31b* islets; rows grouped by cluster theme.

***Supplemental Figure S8: Gpr31b^+^ islet macrophage subsets.* (A)** UMAP of NOD islet macrophage subclusters: annotation (*left*), *Gpr31b*^+^ cells in red (*middle*), and the monocyte-derived Mac-3(Cxcl9)-2 and resident-derived Mac-3(Cxcl9)-3 subclusters (*right*). **(B)** *Gpr31b*^+^ cells per subcluster. **(C)** Marker gene expression by subcluster; dot color, mean expression; dot size, percentage of cells detected. **(D)** hallmark GSEA, *Gpr31b*^+^ versus *Gpr31b*^−^ islet macrophages.

***Supplemental Figure S9: Additional analysis of Gpr31b KO macrophage RNA sequencing.* (A)** PCA of M0 macrophages. **(B)** Volcano plot of differentially expressed genes in M0-like macrophages from WT and *Gpr31b* KO mice. Genes that meet the threshold of absolute fold change ≥1.5 and adjusted P-value <0.05 are in red; genes that only meet the adjusted P-value threshold are in blue. **(C)** Leading-edge genes, hallmark IFN-γ (*left*) and IFN-α (*right*) response. **(D)** Leading-edge genes, hallmark glycolysis (*left*) and fatty acid metabolism (*right*). **(E)** pathlinkR Reactome network, M1-like. **(F)** Sequence logos for motifs identified by HOMER. IRF, interferon regulatory factor.

Supplemental Figure S10: Domain and gene ontology consequences of alternative splicing in Gpr31b KO macrophages and effects alox12 morpholino in zebrafish. (A) Protein domains affected by alternative splicing per gene, for domain gain in KO (*top left*), domain loss in KO (*top right*), and domains affected in both (*bottom*). **(B)** GO overrepresentation of spliced genes; bars, alternatively spliced gene count; blue dots, adjusted P-value. **(C)** Gene-level expression in transcripts per million (*top*) and isoform usage (*bottom*) for *Rapgef2*, *Arrb1*, *Gnai2*, *Camsap2*, *Mtmr2*, *Babam2*, and *Tnfrsf14*. **(D)** Isoform structures for the genes in **(C)**, annotated for protein domains and predicted topology. Data presented as mean with SEM. Single data points represent individual mice. **(E)** Macrophage count from the tailfin transection assay in zebrafish injected with control or *alox12* MO and treated with vehicle or 12-HETE (3 µM). Single data points represent individual zebrafish. Data presented as mean with SEM. Statistical significance determined by one-way ANOVA.

***Supplemental Figure S11: GPR31 is expressed in cDC1 cells in human spleen and pancreatic lymph node.*** CITE-seq of human spleen, pancreatic lymph node, mesenteric lymph node, and superior mesenteric artery lymph node. **(A)** UMAP: annotation (*left*) and *GPR31*^+^ cells in red (*right*). **(B)** *GPR31*^+^ cells by population and tissue; dot color, percentage *GPR31*^+^; dot size, cells assayed; empty circles indicate no *GPR31*^+^ cells. **(C)** *GPR31* expression in cDC1, spleen and pancreatic lymph node, by donor. Non-diabetic (ND; n=5) versus AAb+ or T1D (n=13). P-values are from exact donor-label permutation of a Poisson mixed model. **(D)** hallmark GSEA, *GPR31*^+^ versus *GPR31*^−^ cDC1. **(E)** Reactome GSEA, *GPR31*^+^ versus *GPR31*^−^ cDC1.

## SUPPLEMENTAL TABLES

***Data File S1***: Differentially expressed genes for RNA sequencing of *Gpr31b* KO islets.

***Data File S2***: Significant Reactome GSEA results for RNA-sequencing of PIC-treated *Gpr31b* KO islets.

***Data File S3***: Differentially expressed genes for RNA sequencing of si*Gpr31b* islets.

***Data File S4***: Differentially expressed genes for RNA sequencing of G4451 islets.

***Data File S5***: Top and bottom gene signatures.

***Data File S6***: Differentially expressed genes for RNA sequencing of *Gpr31b* KO BMDMs.

***Data File S7***: Significant Reactome GSEA results for RNA-sequencing of *Gpr31b* KO M1-like BMDMs.

## Notes

### Competing Interest Statement

The authors have declared no competing interest.

### Summary of Updates

Substantial changes have been made throughout the manuscript. All figures have been revised. Added new bioinformatics experiments, and NOD mice treated with siRNA and inhibitor.

## REFERENCES

1. Lee H, Lee YS, Harenda Q, Pietrzak S, Oktay HZ, Schreiber S, et al. Beta Cell Dedifferentiation Induced by IRE1α Deletion Prevents Type 1 Diabetes. Cell Metabolism. 2020 Apr 7;31(4):822–836.e5. doi:10.1016/j.cmet.2020.03.002 PubMed PMID: 32220307.

2. Tersey SA, Nishiki Y, Templin AT, Cabrera SM, Stull ND, Colvin SC, et al. Islet β-cell endoplasmic reticulum stress precedes the onset of type 1 diabetes in the nonobese diabetic mouse model. Diabetes. 2012 Apr;61(4):818–27. doi:10.2337/db11-1293 PubMed PMID: 22442300; PubMed Central PMCID: PMC3314371.

3. Kulkarni A, Pineros AR, Walsh MA, Casimiro I, Ibrahim S, Hernandez-Perez M, et al. 12-Lipoxygenase governs the innate immune pathogenesis of islet inflammation and autoimmune diabetes. JCI Insight. 2021 Jul 22;6(14):e147812. doi:10.1172/jci.insight.147812 PubMed PMID: 34128835; PubMed Central PMCID: PMC8410073.

4. Piñeros AR, Kulkarni A, Gao H, Orr KS, Glenn L, Huang F, et al. Proinflammatory signaling in islet β cells propagates invasion of pathogenic immune cells in autoimmune diabetes. Cell Rep. 2022 Jun 28;39(13):111011. doi:10.1016/j.celrep.2022.111011 PubMed PMID: 35767947; PubMed Central PMCID: PMC9297711.

5. Nikolic T, Geutskens SB, van Rooijen N, Drexhage HA, Leenen PJM. Dendritic cells and macrophages are essential for the retention of lymphocytes in (peri)-insulitis of the nonobese diabetic mouse: a phagocyte depletion study. Lab Invest. 2005 Apr;85(4):487– 501. doi:10.1038/labinvest.3700238 PubMed PMID: 15654358.

6. Kulkarni A, Pineros AR, Walsh MA, Casimiro I, Ibrahim S, Hernandez-Perez M, et al. 12-Lipoxygenase governs the innate immune pathogenesis of islet inflammation and autoimmune diabetes. JCI Insight. 2021 Jun 15;147812. doi:10.1172/jci.insight.147812 PubMed PMID: 34128835.

7. Carrero JA, McCarthy DP, Ferris ST, Wan X, Hu H, Zinselmeyer BH, et al. Resident macrophages of pancreatic islets have a seminal role in the initiation of autoimmune diabetes of NOD mice. Proceedings of the National Academy of Sciences. 2017 Nov 28;114(48):E10418–27. doi:10.1073/pnas.1713543114

8. Forlenza GP, McVean J, Beck RW, Bauza C, Bailey R, Buckingham B, et al. Effect of Verapamil on Pancreatic Beta Cell Function in Newly Diagnosed Pediatric Type 1 Diabetes: A Randomized Clinical Trial. JAMA. 2023 Mar 28;329(12):990–9. doi:10.1001/jama.2023.2064 PubMed PMID: 36826844; PubMed Central PMCID: PMC9960020.

9. Sims EK, Kulkarni A, Hull A, Woerner SE, Cabrera S, Mastrandrea LD, et al. Inhibition of polyamine biosynthesis preserves β cell function in type 1 diabetes. Cell Reports Medicine. 2023 Nov 1;101261. doi:10.1016/j.xcrm.2023.101261

10. Dobrian AD, Lieb DC, Cole BK, Taylor-Fishwick DA, Chakrabarti SK, Nadler JL. Functional and pathological roles of the 12– and 15-lipoxygenases. Prog Lipid Res. 2011 Jan;50(1):115–31. doi:10.1016/j.plipres.2010.10.005 PubMed PMID: 20970452; PubMed Central PMCID: PMC3012140.

11. Kulkarni A, Nadler JL, Mirmira RG, Casimiro I. Regulation of Tissue Inflammation by 12-Lipoxygenases. Biomolecules. 2021 May 11;11(5):717. doi:10.3390/biom11050717 PubMed PMID: 34064822; PubMed Central PMCID: PMC8150372.

12. Brash AR. Lipoxygenases: occurrence, functions, catalysis, and acquisition of substrate. J Biol Chem. 1999 Aug 20;274(34):23679–82. PubMed PMID: 10446122.

13. Yamamoto S, Suzuki H, Ueda N. Arachidonate 12-lipoxygenases. Prog Lipid Res. 1997 Mar;36(1):23–41. PubMed PMID: 9373619.

14. Grzesik WJ, Nadler JL, Machida Y, Nadler JL, Imai Y, Morris MA. Expression pattern of 12-lipoxygenase in human islets with type 1 diabetes and type 2 diabetes. J Clin Endocrinol Metab. 2015 Mar;100(3):E387–395. doi:10.1210/jc.2014-3630 PubMed PMID: 25532042.

15. Hennessy E, Rakovac Tisdall A, Murphy N, Carroll A, O’Gorman D, Breen L, et al. Elevated 12-hydroxyeicosatetraenoic acid (12-HETE) levels in serum of individuals with newly diagnosed Type 1 diabetes. Diabet Med. 2017 Feb;34(2):292–4. doi:10.1111/dme.13177 PubMed PMID: 27353008.

16. Ma K, Xiao A, Park SH, Glenn L, Jackson L, Barot T, et al. 12-Lipoxygenase Inhibitor Improves Functions of Cytokine-Treated Human Islets and Type 2 Diabetic Islets. J Clin Endocrinol Metab. 2017 01;102(8):2789–97. doi:10.1210/jc.2017-00267 PubMed PMID: 28609824; PubMed Central PMCID: PMC5546865.

17. Nargis T, Muralidharan C, Enriquez JR, Wang JE, Kaylan KB, Chakraborty A, et al. 12-Lipoxygenase inhibition delays onset of autoimmune diabetes in human gene replacement mice. JCI Insight. 2024 Nov 12;9(24):e185299. doi:10.1172/jci.insight.185299 PubMed PMID: 39531315.

18. Natarajan R, Nadler JL. Lipid inflammatory mediators in diabetic vascular disease. Arterioscler Thromb Vasc Biol. 2004 Sep;24(9):1542–8. doi:10.1161/01.ATV.0000133606.69732.4c PubMed PMID: 15166011.

19. Wen Y, Gu J, Chakrabarti SK, Aylor K, Marshall J, Takahashi Y, et al. The role of 12/15-lipoxygenase in the expression of interleukin-6 and tumor necrosis factor-alpha in macrophages. Endocrinology. 2007 Mar;148(3):1313–22. doi:10.1210/en.2006-0665 PubMed PMID: 17170102.

20. Tersey SA, Maier B, Nishiki Y, Maganti AV, Nadler JL, Mirmira RG. 12-Lipoxygenase Promotes Obesity-Induced Oxidative Stress in Pancreatic Islets. Mol Cell Biol. 2014 Jul 28. doi:10.1128/MCB.00157-14 PubMed PMID: 25071151.

21. Guo Y, Zhang W, Giroux C, Cai Y, Ekambaram P, Dilly AK, et al. Identification of the orphan G protein-coupled receptor GPR31 as a receptor for 12-(S)-hydroxyeicosatetraenoic acid. J Biol Chem. 2011 Sep 30;286(39):33832–40. doi:10.1074/jbc.M110.216564 PubMed PMID: 21712392; PubMed Central PMCID: PMC3190773.

22. Zhang XJ, Cheng X, Yan ZZ, Fang J, Wang X, Wang W, et al. An ALOX12-12-HETE-GPR31 signaling axis is a key mediator of hepatic ischemia-reperfusion injury. Nat Med. 2018;24(1):73–83. doi:10.1038/nm.4451 PubMed PMID: 29227475.

23. Ying W, Lee YS, Dong Y, Seidman JS, Yang M, Isaac R, et al. Expansion of Islet-Resident Macrophages Leads to Inflammation Affecting β Cell Proliferation and Function in Obesity. Cell Metab. 2019 Feb 5;29(2):457–474.e5. doi:10.1016/j.cmet.2018.12.003 PubMed PMID: 30595478; PubMed Central PMCID: PMC6701710.

24. Hernandez-Perez M, Kulkarni A, Samala N, Sorrell C, El K, Haider I, et al. A 12-lipoxygenase-Gpr31 signaling axis is required for pancreatic organogenesis in the zebrafish. The FASEB Journal. 2020;34(11):14850–62. doi:10.1096/fj.201902308RR

25. Nunemaker CS, Chen M, Pei H, Kimble SD, Keller SR, Carter JD, et al. 12-Lipoxygenase-knockout mice are resistant to inflammatory effects of obesity induced by Western diet. Am J Physiol Endocrinol Metab. 2008 Nov;295(5):E1065–1075. doi:10.1152/ajpendo.90371.2008 PubMed PMID: 18780776; PubMed Central PMCID: PMC2584815.

26. Hernandez-Perez M, Chopra G, Fine J, Conteh AM, Anderson RM, Linnemann AK, et al. Inhibition of 12/15-Lipoxygenase Protects Against β-Cell Oxidative Stress and Glycemic Deterioration in Mouse Models of Type 1 Diabetes. Diabetes. 2017;66(11):2875–87. doi:10.2337/db17-0215 PubMed PMID: 28842399; PubMed Central PMCID: PMC5652601.

27. Ma K, Nunemaker CS, Wu R, Chakrabarti SK, Taylor-Fishwick DA, Nadler JL. 12-Lipoxygenase Products Reduce Insulin Secretion and {beta}-Cell Viability in Human Islets. J Clin Endocrinol Metab. 2010 Feb;95(2):887–93. doi:10.1210/jc.2009-1102 PubMed PMID: 20089617; PubMed Central PMCID: PMC2840856.

28. Wang C, Yosef N, Gaublomme J, Wu C, Lee Y, Clish CB, et al. CD5L/AIM Regulates Lipid Biosynthesis and Restrains Th17 Cell Pathogenicity. Cell. 2015 Dec;163(6):1413–27. doi:10.1016/j.cell.2015.10.068

29. Sanjurjo L, Aran G, Roher N, Valledor AF, Sarrias MR. AIM/CD5L: a key protein in the control of immune homeostasis and inflammatory disease. Journal of Leukocyte Biology. 2015 Aug 1;98(2):173–84. doi:10.1189/jlb.3RU0215-074R

30. Blimkie TM, An A, Hancock REW. Facilitating pathway and network based analysis of RNA-Seq data with pathlinkR. PLOS Computational Biology. 2024 Sep 16;20(9):e1012422. doi:10.1371/journal.pcbi.1012422

31. Ortis F, Naamane N, Flamez D, Ladrière L, Moore F, Cunha DA, et al. Cytokines interleukin-1beta and tumor necrosis factor-alpha regulate different transcriptional and alternative splicing networks in primary beta-cells. Diabetes. 2010 Feb;59(2):358–74. doi:10.2337/db09-1159 PubMed PMID: 19934004; PubMed Central PMCID: PMC2809955.

32. Wu W, Syed F, Simpson E, Lee CC, Liu J, Chang G, et al. Impact of Proinflammatory Cytokines on Alternative Splicing Patterns in Human Islets. Diabetes. 2021 Oct 25;71(1):116–27. doi:10.2337/db20-0847

33. Han C, Gilis J, Delgado EI, Clement L, Vitting-Seerup K. IsoformSwitchAnalyzeR v2: Analysis of Functional Isoform Changes in Long-read and Single-cell Sequencing Data [Internet]. bioRxiv; 2025 [cited 2026 Feb 2]. p. 2025.12.08.693027. Available from: https://www.biorxiv.org/content/10.64898/2025.12.08.693027v1 doi:10.64898/2025.12.08.693027

34. Reyes A, Anders S, Weatheritt RJ, Gibson TJ, Steinmetz LM, Huber W. Drift and conservation of differential exon usage across tissues in primate species. Proc Natl Acad Sci USA. 2013 Sep 17;110(38):15377–82. doi:10.1073/pnas.1307202110

35. Maier B, Ogihara T, Trace AP, Tersey SA, Robbins RD, Chakrabarti SK, et al. The unique hypusine modification of eIF5A promotes islet beta cell inflammation and dysfunction in mice. J Clin Invest. 2010 Jun;120(6):2156–70. doi:10.1172/JCI38924

36. Calderon B, Suri A, Miller MJ, Unanue ER. Dendritic cells in islets of Langerhans constitutively present β cell-derived peptides bound to their class II MHC molecules. PNAS. 2008 Apr 22;105(16):6121–6. doi:10.1073/pnas.0801973105 PubMed PMID: 18427107.

37. Takasu N, Komiya I, Asawa T, Nagasawa Y, Yamada T. Streptozocin– and alloxan-induced H2O2 generation and DNA fragmentation in pancreatic islets. H2O2 as mediator for DNA fragmentation. Diabetes. 1991 Sep;40(9):1141–5. doi:10.2337/diab.40.9.1141 PubMed PMID: 1834504.

38. Szkudelski T. The mechanism of alloxan and streptozotocin action in B cells of the rat pancreas. Physiol Res. 2001;50(6):537–46. PubMed PMID: 11829314.

39. Bleich D, Chen S, Zipser B, Sun D, Funk CD, Nadler JL. Resistance to type 1 diabetes induction in 12-lipoxygenase knockout mice. J Clin Invest. 1999 May 15;103(10):1431–6. doi:10.1172/JCI5241 PubMed PMID: 10330425; PubMed Central PMCID: PMC408453.

40. Zhang XJ, Fu J, Cheng X, Shen H, Yang H, Wang K, et al. Integrated screening identifies GPR31 as a key driver and druggable target for metabolic dysfunction–associated steatohepatitis. J Clin Invest. 2025 Sep 2;135(17). doi:10.1172/JCI173193 PubMed PMID: 40892510.

41. Rui J, Deng S, Arazi A, Perdigoto AL, Liu Z, Herold KC. β Cells that Resist Immunological Attack Develop during Progression of Autoimmune Diabetes in NOD Mice. Cell Metabolism. 2017 Mar 7;25(3):727–38. doi:10.1016/j.cmet.2017.01.005 PubMed PMID: 28190773.

42. Colli ML, Hill JLE, Marroquí L, Chaffey J, Santos RSD, Leete P, et al. PDL1 is expressed in the islets of people with type 1 diabetes and is up-regulated by interferons-α and-γ via IRF1 induction. eBioMedicine. 2018 Oct 1;36:367–75. doi:10.1016/j.ebiom.2018.09.040 PubMed PMID: 30269996.

43. Yoshida H, Lareau CA, Ramirez RN, Rose SA, Maier B, Wroblewska A, et al. The cis-Regulatory Atlas of the Mouse Immune System. Cell. 2019 Feb 7;176(4):897–912.e20. doi:10.1016/j.cell.2018.12.036 PubMed PMID: 30686579; PubMed Central PMCID: PMC6785993.

44. Zakharov PN, Hu H, Wan X, Unanue ER. Single-cell RNA sequencing of murine islets shows high cellular complexity at all stages of autoimmune diabetes. Journal of Experimental Medicine. 2020 Apr 6;217(6):e20192362. doi:10.1084/jem.20192362

45. Szklarczyk D, Kirsch R, Koutrouli M, Nastou K, Mehryary F, Hachilif R, et al. The STRING database in 2023: protein-protein association networks and functional enrichment analyses for any sequenced genome of interest. Nucleic Acids Res. 2023 Jan 6;51(D1):D638–46. doi:10.1093/nar/gkac1000 PubMed PMID: 36370105; PubMed Central PMCID: PMC9825434.

46. Abedi M, Rai P, Zhou Y, Liu C, Johnson I, Chandra A, et al. Joint Profiling of Gene Expression and Chromatin Accessibility in Pancreatic Lymph Nodes and Spleens of Human Type 1 Diabetes. Sci Immunol. 2025 Nov 21;10(113):eadz0472. doi:10.1126/sciimmunol.adz0472 PubMed PMID: 41270190; PubMed Central PMCID: PMC12766627.

47. McDuffie M, Maybee NA, Keller SR, Stevens BK, Garmey JC, Morris MA, et al. Nonobese diabetic (NOD) mice congenic for a targeted deletion of 12/15-lipoxygenase are protected from autoimmune diabetes. Diabetes. 2008 Jan;57(1):199–208. doi:10.2337/db07-0830 PubMed PMID: 17940120; PubMed Central PMCID: PMC2993320.

48. Green-Mitchell SM, Tersey SA, Cole BK, Ma K, Kuhn NS, Cunningham TD, et al. Deletion of 12/15-lipoxygenase alters macrophage and islet function in NOD-Alox15(null) mice, leading to protection against type 1 diabetes development. PLoS ONE. 2013;8(2):e56763. doi:10.1371/journal.pone.0056763 PubMed PMID: 23437231; PubMed Central PMCID: PMC3578926.

49. Atouf F, Czernichow P, Scharfmann R. Expression of Neuronal Traits in Pancreatic Beta Cells. Journal of Biological Chemistry. 1997 Jan;272(3):1929–34. doi:10.1074/jbc.272.3.1929

50. Martens GA, Jiang L, Hellemans KH, Stangé G, Heimberg H, Nielsen FC, et al. Clusters of Conserved Beta Cell Marker Genes for Assessment of Beta Cell Phenotype. Tjwa M, editor. PLoS ONE. 2011 Sep 2;6(9):e24134. doi:10.1371/journal.pone.0024134

51. Van Arensbergen J, García-Hurtado J, Moran I, Maestro MA, Xu X, Van De Casteele M, et al. Derepression of Polycomb targets during pancreatic organogenesis allows insulin-producing beta-cells to adopt a neural gene activity program. Genome Res. 2010 Jun;20(6):722–32. doi:10.1101/gr.101709.109

52. Webster KL, Mirmira RG. Beta cell dedifferentiation in type 1 diabetes: sacrificing function for survival? Front Endocrinol. 2024 Jun 6;15. doi:10.3389/fendo.2024.1427723

53. Spurrell M, Tsang JS, Herold KC. A surviving beta cell subpopulation enriched in patients with T1D [Internet]. Immunology; 2026 [cited 2026 May 29]. Available from: http://biorxiv.org/lookup/doi/10.64898/2026.05.15.725449 doi:10.64898/2026.05.15.725449

54. Wang CJ, Chou FC, Chu CH, Wu JC, Lin SH, Chang DM, et al. Protective Role of Programmed Death 1 Ligand 1 (PD-L1)in Nonobese Diabetic Mice: The Paradox in Transgenic Models. Diabetes. 2008 Jul 1;57(7):1861–9. doi:10.2337/db07-1260

55. Enriquez JR, Zhou Z, Nelson JB, Huang F, Figatner KT, Chakraborty A, et al. Messenger RNA delivery to islet β cells using conjugated lipid nanoparticles. CR Med. 2026 Feb 20;0(0). doi:10.1016/j.xcrm.2026.102634

56. Michelucci A, Cordes T, Ghelfi J, Pailot A, Reiling N, Goldmann O, et al. Immune-responsive gene 1 protein links metabolism to immunity by catalyzing itaconic acid production. Proc Natl Acad Sci U S A. 2013 May 7;110(19):7820–5. doi:10.1073/pnas.1218599110 PubMed PMID: 23610393; PubMed Central PMCID: PMC3651434.

57. Bourner LA, Chung LA, Long H, McGettrick AF, Xiao J, Roth K, et al. Endogenously produced itaconate negatively regulates innate-driven cytokine production and drives global ubiquitination in human macrophages. Cell Rep. 2024 Aug 27;43(8):114570. doi:10.1016/j.celrep.2024.114570 PubMed PMID: 39093697.

58. Juan-Mateu J, Alvelos MI, Turatsinze JV, Villate O, Lizarraga-Mollinedo E, Grieco FA, et al. SRp55 Regulates a Splicing Network That Controls Human Pancreatic β-Cell Function and Survival. Diabetes. 2018 Mar 1;67(3):423–36. doi:10.2337/db17-0736

59. Carré A, Samassa F, Zhou Z, Perez-Hernandez J, Lekka C, Manganaro A, et al. Interferon-α promotes HLA-B-restricted presentation of conventional and alternative antigens in human pancreatic β-cells. Nat Commun. 2025 Jan 17;16(1):765. doi:10.1038/s41467-025-55908-9

60. Guyer P, Arribas-Layton D, Manganaro A, Speake C, Lord S, Eizirik DL, et al. Recognition of mRNA Splice Variant and Secretory Granule Epitopes by CD4+ T Cells in Type 1 Diabetes. Diabetes. 2023 Jan 1;72(1):85–96. doi:10.2337/db22-0191

61. Giancotti LA, Lauro F, Olayide I, Zhang J, Arnatt CK, Salvemini D. 12-(S)-Hydroxyeicosatetraenoic Acid and GPR31 Signaling in Spinal Cord in Neuropathic Pain. The Journal of Pharmacology and Experimental Therapeutics. 2024 Mar;388(3):765–73. doi:10.1124/jpet.123.001853

62. Anderson PJ, Xiao P, Zhong YN, Kaakati AN, Alfonso-DeSouza J, Patino A, et al. β-Arrestin condensates regulate G-protein-coupled receptor function. Nature. 2026 Jul;655(8122):525–34. doi:10.1038/s41586-026-10539-y

63. Herold KC, Delong T, Perdigoto AL, Biru N, Brusko TM, Walker LSK. The immunology of type 1 diabetes. Nature reviews Immunology. 2024 Feb 2;24(6):435. doi:10.1038/s41577-023-00985-4 PubMed PMID: 38308004.

64. Calderon B, Carrero JA, Unanue ER. The Central Role of Antigen Presentation in Islets of Langerhans in Autoimmune Diabetes. Curr Opin Immunol. 2014 Feb;26:32–40. doi:10.1016/j.coi.2013.10.011 PubMed PMID: 24556398; PubMed Central PMCID: PMC4118295.

65. Apaolaza PS, Balcacean D, Zapardiel-Gonzalo J, Nelson G, Lenchik N, Akhbari P, et al. Islet expression of type I interferon response sensors is associated with immune infiltration and viral infection in type 1 diabetes. Sci Adv. 2021 Feb 26;7(9):eabd6527. doi:10.1126/sciadv.abd6527

66. Ferreira RC, Guo H, Coulson RMR, Smyth DJ, Pekalski ML, Burren OS, et al. A type I interferon transcriptional signature precedes autoimmunity in children genetically at risk for type 1 diabetes. Diabetes. 2014 Jul;63(7):2538–50. doi:10.2337/db13-1777 PubMed PMID: 24561305; PubMed Central PMCID: PMC4066333.

67. Marro BS, Ware BC, Zak J, de la Torre JC, Rosen H, Oldstone MBA. Progression of type 1 diabetes from the prediabetic stage is controlled by interferon-α signaling. Proc Natl Acad Sci U S A. 2017 Apr 4;114(14):3708–13. doi:10.1073/pnas.1700878114 PubMed PMID: 28325871; PubMed Central PMCID: PMC5389329.

68. Coomans de Brachène A, Dos Santos RS, Marroqui L, Colli ML, Marselli L, Mirmira RG, et al. IFN-α induces a preferential long-lasting expression of MHC class I in human pancreatic beta cells. Diabetologia. 2018 Mar;61(3):636–40. doi:10.1007/s00125-017-4536-4 PubMed PMID: 29305625; PubMed Central PMCID: PMC6241216.

69. Marroqui L, Dos Santos RS, Op De Beeck A, Coomans De Brachène A, Marselli L, Marchetti P, et al. Interferon-α mediates human beta cell HLA class I overexpression, endoplasmic reticulum stress and apoptosis, three hallmarks of early human type 1 diabetes. Diabetologia. 2017 Apr;60(4):656–67. doi:10.1007/s00125-016-4201-3

70. Wagner LE, Melnyk O, Turner A, Duffett BE, Muralidharan C, Martinez-Irizarry MM, et al. IFN-α Induces Heterogenous ROS Production in Human β-Cells [Internet]. bioRxiv; 2025 [cited 2025 Mar 10]. p. 2025.02.19.639120. Available from: https://www.biorxiv.org/content/10.1101/2025.02.19.639120v1 doi:10.1101/2025.02.19.639120

71. Zinselmeyer BH, Vomund AN, Saunders BT, Johnson MW, Carrero JA, Unanue ER. The resident macrophages in murine pancreatic islets are constantly probing their local environment, capturing beta cell granules and blood particles. Diabetologia. 2018 Jun;61(6):1374–83. doi:10.1007/s00125-018-4592-4

72. Ferris ST, Carrero JA, Mohan JF, Calderon B, Murphy KM, Unanue ER. A Minor Subset of *Batf3*-Dependent Antigen-Presenting Cells in Islets of Langerhans Is Essential for the Development of Autoimmune Diabetes. Immunity. 2014 Oct 16;41(4):657–69. doi:10.1016/j.immuni.2014.09.012

73. Ferris ST, Liu T, Chen J, Ohara RA, Ou F, Wu R, et al. WDFY4 deficiency in NOD mice ameliorates autoimmune diabetes and insulitis. Proc Natl Acad Sci USA. 2023 Mar 28;120(13):e2219956120. doi:10.1073/pnas.2219956120

74. Steenbuck N, Damond N, Engler S, Kusmartseva I, Posgai AL, Drotar DM, et al. Imaging mass cytometry reveals functional and immunological changes during type 1 diabetes progression in human pancreata. Nat Metab. 2026 Jul 2;1–18. doi:10.1038/s42255-026-01559-z

75. Oguro-Igashira E, Murakami M, Mori R, Kuwahara R, Kihara T, Kohara M, et al. The pyruvate–GPR31 axis promotes transepithelial dendrite formation in human intestinal dendritic cells. Proceedings of the National Academy of Sciences. 2024 Oct 29;121(44):e2318767121. doi:10.1073/pnas.2318767121

76. Morita N, Umemoto E, Fujita S, Hayashi A, Kikuta J, Kimura I, et al. GPR31-dependent dendrite protrusion of intestinal CX3CR1+ cells by bacterial metabolites. Nature. 2019 Feb;566(7742):110–4. doi:10.1038/s41586-019-0884-1

77. Stull ND, Breite A, McCarthy R, Tersey SA, Mirmira RG. Mouse islet of Langerhans isolation using a combination of purified collagenase and neutral protease. J Vis Exp. 2012 Sep 7;(67):4137. doi:10.3791/4137 PubMed PMID: 22987198; PubMed Central PMCID: PMC3490250.

78. Ellett F, Pase L, Hayman JW, Andrianopoulos A, Lieschke GJ. mpeg1 promoter transgenes direct macrophage-lineage expression in zebrafish. Blood. 2011 Jan 27;117(4):e49–56. doi:10.1182/blood-2010-10-314120 PubMed PMID: 21084707; PubMed Central PMCID: PMC3056479.

79. Ray A, Dittel BN. Isolation of mouse peritoneal cavity cells. J Vis Exp. 2010 Jan 28;(35):1488. doi:10.3791/1488 PubMed PMID: 20110936; PubMed Central PMCID: PMC3152216.

80. Tersey SA, Levasseur EM, Syed F, Farb TB, Orr KS, Nelson JB, et al. Episodic β-cell death and dedifferentiation during diet-induced obesity and dysglycemia in male mice. FASEB J. 2018 May 29;fj201800150RR. doi:10.1096/fj.201800150RR PubMed PMID: 29812970.

81. Schindelin J, Arganda-Carreras I, Frise E, Kaynig V, Longair M, Pietzsch T, et al. Fiji: an open-source platform for biological-image analysis. Nat Methods. 2012 Jul;9(7):676–82. doi:10.1038/nmeth.2019

82. Stirling DR, Swain-Bowden MJ, Lucas AM, Carpenter AE, Cimini BA, Goodman A. CellProfiler 4: improvements in speed, utility and usability. BMC Bioinformatics. 2021 Sep 10;22(1):433. doi:10.1186/s12859-021-04344-9 PubMed PMID: 34507520; PubMed Central PMCID: PMC8431850.

83. Cabrera SM, Colvin SC, Tersey SA, Maier B, Nadler JL, Mirmira RG. Effects of combination therapy with dipeptidyl peptidase-IV and histone deacetylase inhibitors in the non-obese diabetic mouse model of type 1 diabetes. Clinical and Experimental Immunology. 2013 Apr 18;172(3):375–82. doi:10.1111/cei.12068

84. Muralidharan C, Huang F, Enriquez JR, Wang JE, Nelson JB, Nargis T, et al. Inhibition of the eukaryotic initiation factor-2-α kinase PERK decreases risk of autoimmune diabetes in mice. Journal of Clinical Investigation. 2024 Jun 18;134(16):e176136. doi:10.1172/JCI176136

85. Bolger AM, Lohse M, Usadel B. Trimmomatic: a flexible trimmer for Illumina sequence data. Bioinformatics. 2014 Aug 1;30(15):2114–20. doi:10.1093/bioinformatics/btu170

86. Martin M. Cutadapt removes adapter sequences from high-throughput sequencing reads. EMBnet j. 2011 May 2;17(1):10. doi:10.14806/ej.17.1.200

87. Dobin A, Davis CA, Schlesinger F, Drenkow J, Zaleski C, Jha S, et al. STAR: ultrafast universal RNA-seq aligner. Bioinformatics. 2013 Jan 1;29(1):15–21. doi:10.1093/bioinformatics/bts635

88. Anders S, Pyl PT, Huber W. HTSeq—a Python framework to work with high-throughput sequencing data. Bioinformatics. 2015 Jan 15;31(2):166–9. doi:10.1093/bioinformatics/btu638

89. Liao Y, Smyth GK, Shi W. featureCounts: an efficient general purpose program for assigning sequence reads to genomic features. Bioinformatics. 2014 Apr 1;30(7):923–30. doi:10.1093/bioinformatics/btt656

90. Love MI, Huber W, Anders S. Moderated estimation of fold change and dispersion for RNA-seq data with DESeq2. Genome Biol. 2014 Dec 5;15(12):550. doi:10.1186/s13059-014-0550-8

91. Zhang Y, Parmigiani G, Johnson WE. ComBat-seq: batch effect adjustment for RNA-seq count data. NAR Genomics and Bioinformatics. 2020 Sep 1;2(3):lqaa078. doi:10.1093/nargab/lqaa078

92. Yu G, Wang LG, Han Y, He QY. clusterProfiler: an R Package for Comparing Biological Themes Among Gene Clusters. OMICS: A Journal of Integrative Biology. 2012 May;16(5):284–7. doi:10.1089/omi.2011.0118

93. Liberzon A, Birger C, Thorvaldsdóttir H, Ghandi M, Mesirov JP, Tamayo P. The Molecular Signatures Database Hallmark Gene Set Collection. Cell Systems. 2015 Dec;1(6):417–25. doi:10.1016/j.cels.2015.12.004

94. The Gene Ontology Consortium, Aleksander SA, Balhoff J, Carbon S, Cherry JM, Drabkin HJ, et al. The Gene Ontology knowledgebase in 2023. Baryshnikova A, editor. GENETICS. 2023 May 4;224(1):iyad031. doi:10.1093/genetics/iyad031

95. Milacic M, Beavers D, Conley P, Gong C, Gillespie M, Griss J, et al. The Reactome Pathway Knowledgebase 2024. Nucleic Acids Res. 2024 Jan 5;52(D1):D672–8. doi:10.1093/nar/gkad1025

96. Heinz S, Benner C, Spann N, Bertolino E, Lin YC, Laslo P, et al. Simple combinations of lineage-determining transcription factors prime cis-regulatory elements required for macrophage and B cell identities. Mol Cell. 2010 May 28;38(4):576–89. doi:10.1016/j.molcel.2010.05.004 PubMed PMID: 20513432; PubMed Central PMCID: PMC2898526.

97. Tremblay BJM. universalmotif: An R package for biological motifanalysis. JOSS. 2024 Aug 19;9(100):7012. doi:10.21105/joss.07012

98. Monaco G, Lee B, Xu W, Mustafah S, Hwang YY, Carré C, et al. RNA-Seq Signatures Normalized by mRNA Abundance Allow Absolute Deconvolution of Human Immune Cell Types. Cell Rep. 2019 Feb 5;26(6):1627–1640.e7. doi:10.1016/j.celrep.2019.01.041 PubMed PMID: 30726743; PubMed Central PMCID: PMC6367568.

99. T.H. Vu H, Sun H, Sharp S, Kudtarkar P, Brusman L, Jurgens J, et al. Single cell expression map of pancreatic islets using data from HPAP, IIDP and Prodo v3.3 [Internet]. 2025 Jun 5. doi:10.5281/zenodo.15596314

100. Kursa MB, Rudnicki WR. Feature Selection with the **Boruta** Package. J Stat Soft. 2010;36(11). doi:10.18637/jss.v036.i11

101. Korotkevich G, Sukhov V, Budin N, Shpak B, Artyomov MN, Sergushichev A. Fast gene set enrichment analysis [Internet]. Bioinformatics; 2016 [cited 2025 Sep 24]. Available from: http://biorxiv.org/lookup/doi/10.1101/060012 doi:10.1101/060012

102. Patil AR, Schug J, Naji A, Kaestner KH, Faryabi RB, Vahedi G. Single-cell expression profiling of islets generated by the Human Pancreas Analysis Program. Nat Metab. 2023 May;5(5):713–5. doi:10.1038/s42255-023-00806-x PubMed PMID: 37188822; PubMed Central PMCID: PMC10731597.

103. Fasolino M, Schwartz GW, Patil AR, Mongia A, Golson ML, Wang YJ, et al. Single-cell multi-omics analysis of human pancreatic islets reveals novel cellular states in type 1 diabetes. Nat Metab. 2022 Feb;4(2):2. doi:10.1038/s42255-022-00531-x

104. Shapira SN, Naji A, Atkinson MA, Powers AC, Kaestner KH. Understanding islet dysfunction in type 2 diabetes through multidimensional pancreatic phenotyping: The Human Pancreas Analysis Program. Cell Metab. 2022 Dec 6;34(12):1906–13. doi:10.1016/j.cmet.2022.09.013 PubMed PMID: 36206763; PubMed Central PMCID: PMC9742126.

105. Kaestner KH, Powers AC, Naji A, HPAP Consortium, Atkinson MA. NIH Initiative to Improve Understanding of the Pancreas, Islet, and Autoimmunity in Type 1 Diabetes: The Human Pancreas Analysis Program (HPAP). Diabetes. 2019 Jul;68(7):1394–402. doi:10.2337/db19-0058 PubMed PMID: 31127054; PubMed Central PMCID: PMC6609987.

