## Supplemental Figures for "The G protein-coupled receptor GPR31 governs inflammatory signaling in the pancreatic islet microenvironment"

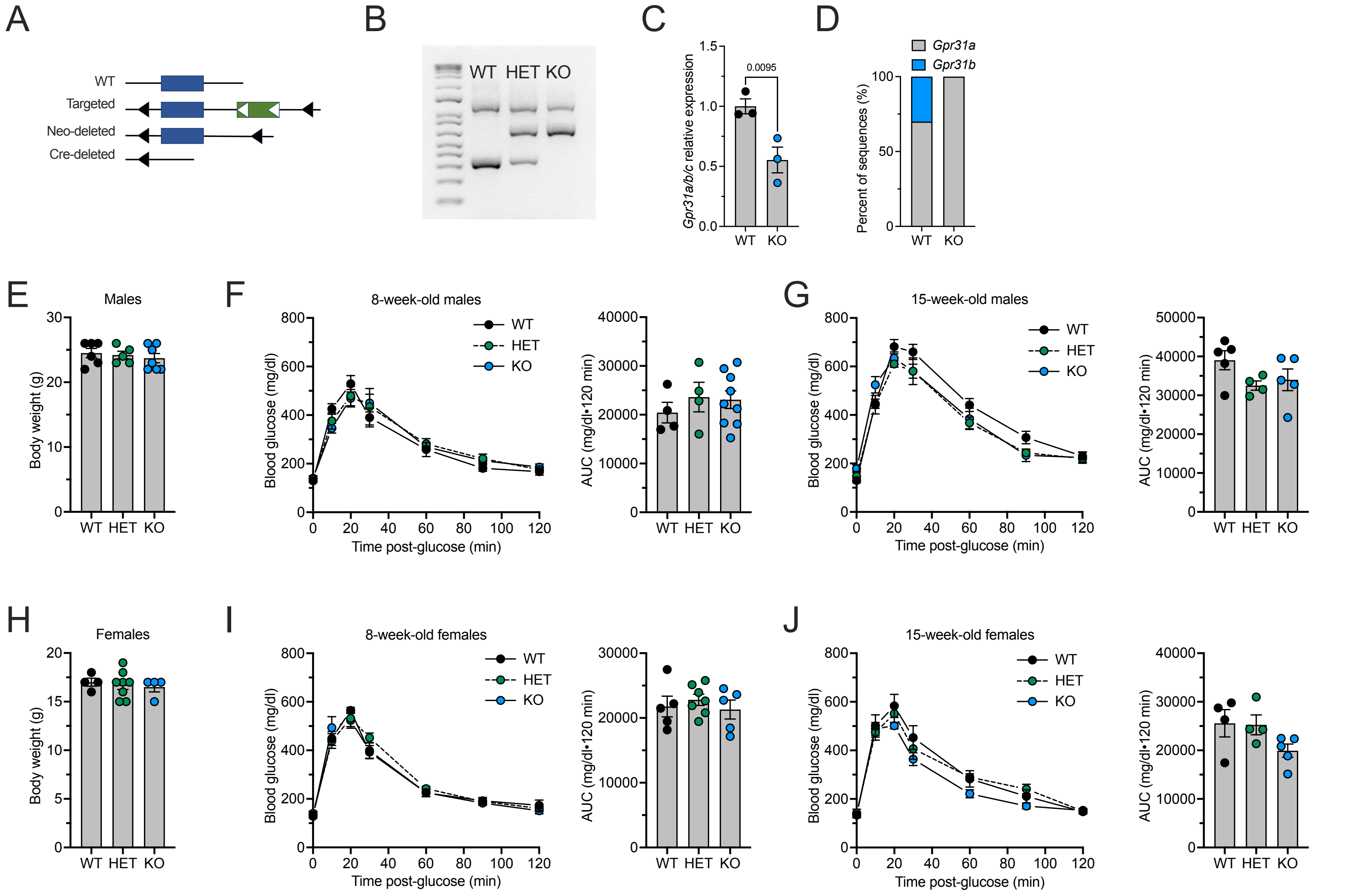

A
WT
Targeted
Neo-deleted
Cre-deleted
B
WT HET KO
C
Gpr31a/b/c relative expression
0.0095
WT KO
D
Gpr31a
Gpr31b
Percent of sequences (%)
WT KO
E
Males
Body weight (g)
WT HET KO
F
8-week-old males
Blood glucose (mg/dl)
Time post-glucose (min)
AUC (mg/dl•120 min)
WT HET KO
G
15-week-old males
Blood glucose (mg/dl)
Time post-glucose (min)
AUC (mg/dl•120 min)
WT HET KO
H
Females
Body weight (g)
WT HET KO
I
8-week-old females
Blood glucose (mg/dl)
Time post-glucose (min)
AUC (mg/dl•120 min)
WT HET KO
J
15-week-old females
Blood glucose (mg/dl)
Time post-glucose (min)
AUC (mg/dl•120 min)
WT HET KO

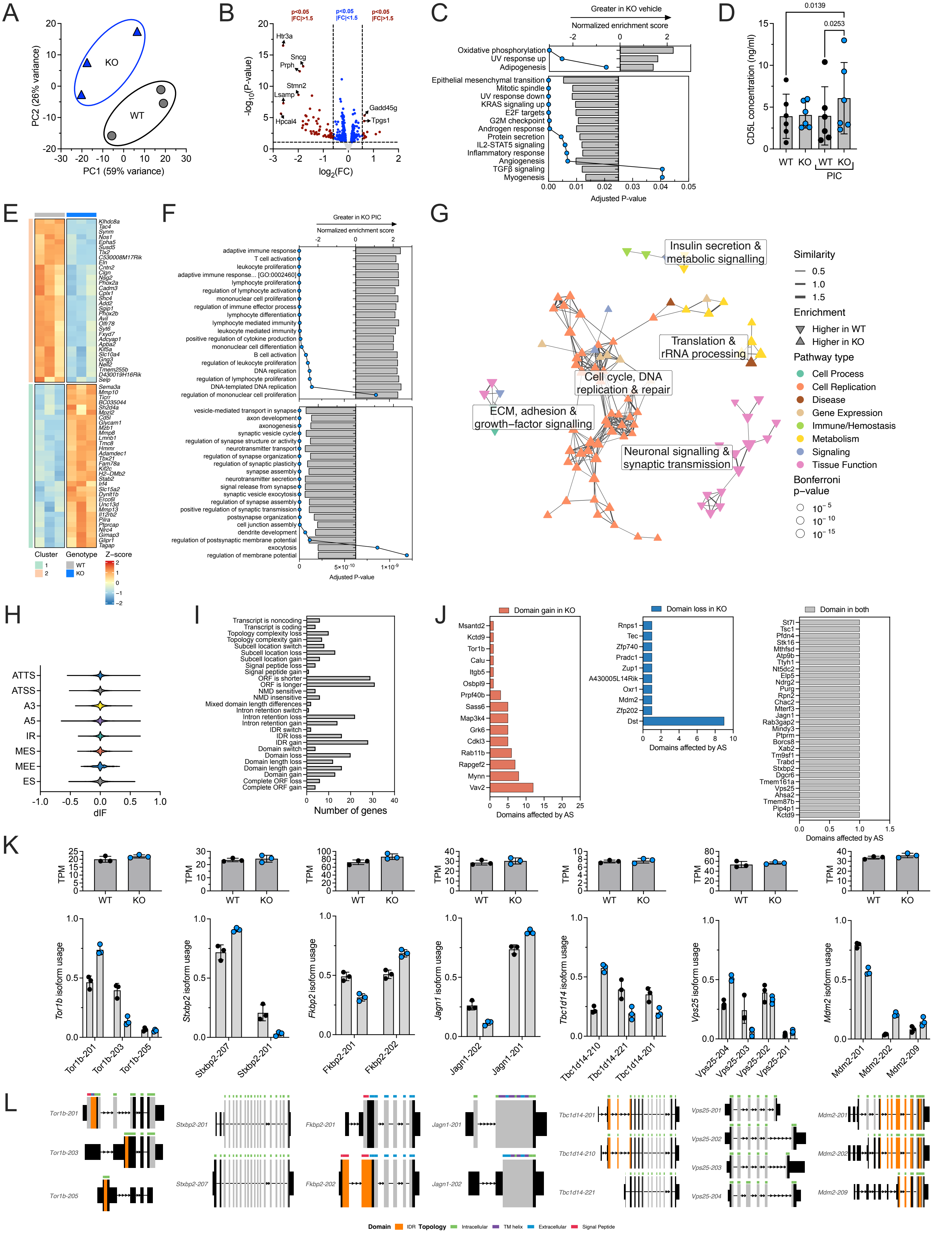
A
KO
WT
PC1 (59% variance)
PC2 (26% variance)
B
p<0.05 |FC|>1.5
p<0.05 |FC|<1.5
p<0.05 |FC|>1.5
Htr3a
Sncg
Prph
Stmn2
Lsamp
Hpcal4
Gadd45g
Tpgs1
-log10(P-value)
log2(FC)
C
Greater in KO vehicle
Normalized enrichment score
Oxidative phosphorylation
UV response up
Adipogenesis
Epithelial mesenchymal transition
Mitotic spindle
UV response down
KRAS signaling up
E2F targets
G2M checkpoint
Androgen response
Protein secretion
IL2-STAT5 signaling
Inflammatory response
Angiogenesis
TGFβ signaling
Myogenesis
Adjusted P-value
D
0.0139
0.0253
CD5L concentration (ng/ml)
WT KO WT KO
PIC
E
Klhdc8a
Tac4
Synm
Nos1
Epha5
Susd5
Tlx2
C530008M17Rik
Eln
Cntn2
Clgn
Nsg2
Phox2a
Cadm3
Cplx1
Shc4
Add2
Sgip1
Phox2b
Avil
Olfr78
Syt6
Fxyd7
Adcyap1
Apba2
Kif5a
Slc10a4
Gng3
Nell2
Tmem255b
D430019H16Rik
Selp
Sema3a
Mmp10
Ticrr
BC035044
Sh2d4a
Mpzl2
Cd5l
Glycam1
Mzb1
Mmp8
Lmnb1
Tmc8
Hmmr
Adamdec1
Tbx21
Fam78a
Kif2c
H2-DMb2
Stab2
Irf4
Slc15a2
Dynlt1b
Ercc6l
Unc13d
Mmp13
Il12rb2
Pilra
Ptprcap
Nlrc4
Gimap3
Glipr1
Tagap
Cluster
Genotype
Z-score
WT
KO
F
Greater in KO PIC
Normalized enrichment score
adaptive immune response
T cell activation
leukocyte proliferation
adaptive immune response... [GO:0002460]
lymphocyte proliferation
regulation of lymphocyte activation
mononuclear cell proliferation
regulation of immune effector process
lymphocyte differentiation
lymphocyte mediated immunity
leukocyte mediated immunity
positive regulation of cytokine production
mononuclear cell differentiation
B cell activation
regulation of leukocyte proliferation
DNA replication
regulation of lymphocyte proliferation
DNA-templated DNA replication
regulation of mononuclear cell proliferation
vesicle-mediated transport in synapse
axon development
axonogenesis
synaptic vesicle cycle
regulation of synapse structure or activity
neurotransmitter transport
regulation of synapse organization
regulation of synaptic plasticity
synapse assembly
neurotransmitter secretion
signal release from synapse
synaptic vesicle exocytosis
regulation of synapse assembly
positive regulation of synaptic transmission
postsynapse organization
cell junction assembly
dendrite development
regulation of postsynaptic membrane potential
exocytosis
regulation of membrane potential
Adjusted P-value
G
Insulin secretion & metabolic signalling
Translation & rRNA processing
Cell cycle, DNA replication & repair
ECM, adhesion & growth-factor signalling
Neuronal signalling & synaptic transmission
Similarity
Enrichment
Higher in WT
Higher in KO
Pathway type
Cell Process
Cell Replication
Disease
Gene Expression
Immune/Hemostasis
Metabolism
Signaling
Tissue Function
Bonferroni p-value
H
ATTS
ATSS
A3
A5
IR
MES
MEE
ES
dIF
I
Transcript is noncoding
Transcript is coding
Topology complexity loss
Topology complexity gain
Subcell location switch
Subcell location loss
Subcell location gain
Signal peptide loss
Signal peptide gain
ORF is shorter
ORF is longer
NMD sensitive
NMD insensitive
Mixed domain length differences
Intron retention switch
Intron retention loss
Intron retention gain
IDR switch
IDR loss
IDR gain
Domain switch
Domain loss
Domain length loss
Domain length gain
Domain gain
Complete ORF loss
Complete ORF gain
Number of genes
J
Domain gain in KO
Msantd2
Kctd9
Tor1b
Calu
Itgb5
Osbpl9
Prpf40b
Sass6
Map3k4
Grk6
Cdkl3
Rab11b
Rapgef2
Mynn
Vav2
Domains affected by AS
Domain loss in KO
Rnps1
Tec
Zfp740
Pradc1
Zup1
A430005L14Rik
Oxr1
Mdm2
Zfp202
Dst
Domains affected by AS
Domain in both
St7l
Tsc1
Pfdn4
Stk16
Mthfsd
Atp9b
Ttyh1
Nt5dc2
Elp5
Ndrg2
Purg
Rpn2
Chac2
Mterf3
Jagn1
Rab3gap2
Mindy3
Ptprm
Borcs8
Xab2
Tm9sf1
Trabd
Stxbp2
Dgcr6
Tmem161a
Vps25
Ahsa2
Tmem87b
Pip4p1
Kctd9
Domains affected by AS
K
TPM
WT KO
Tor1b isoform usage
Tor1b-201 Tor1b-203 Tor1b-205
Stxbp2 isoform usage
Stxbp2-207 Stxbp2-201
Fkbp2 isoform usage
Fkbp2-201 Fkbp2-202
Jagn1 isoform usage
Jagn1-202 Jagn1-201
Tbc1d14 isoform usage
Tbc1d14-210 Tbc1d14-221 Tbc1d14-201
Vps25 isoform usage
Vps25-204 Vps25-203 Vps25-202 Vps25-201
Mdm2 isoform usage
Mdm2-201 Mdm2-202 Mdm2-209
L
Tor1b-201
Tor1b-203
Tor1b-205
Stxbp2-201
Stxbp2-207
Fkbp2-201
Fkbp2-202
Jagn1-201
Jagn1-202
Tbc1d14-201
Tbc1d14-210
Tbc1d14-221
Vps25-201
Vps25-202
Vps25-203
Vps25-204
Mdm2-201
Mdm2-202
Mdm2-209
Domain IDR Topology Intracellular TM helix Extracellular Signal Peptide

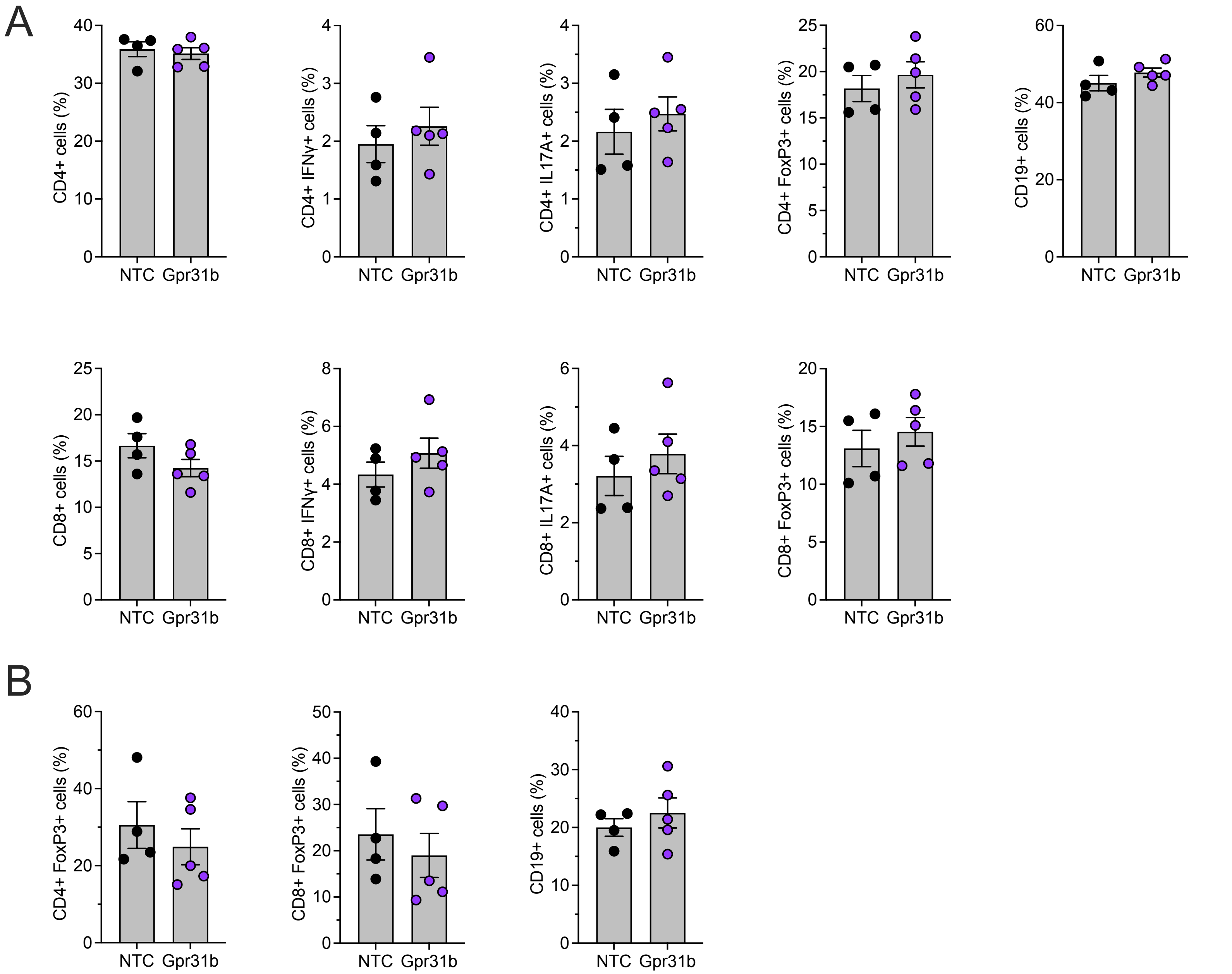

A
CD4+ cells (%)
CD4+ IFNγ+ cells (%)
CD4+ IL17A+ cells (%)
CD4+ FoxP3+ cells (%)
CD19+ cells (%)
CD8+ cells (%)
CD8+ IFNγ+ cells (%)
CD8+ IL17A+ cells (%)
CD8+ FoxP3+ cells (%)
B
CD4+ FoxP3+ cells (%)
CD8+ FoxP3+ cells (%)
CD19+ cells (%)
NTC
Gpr31b

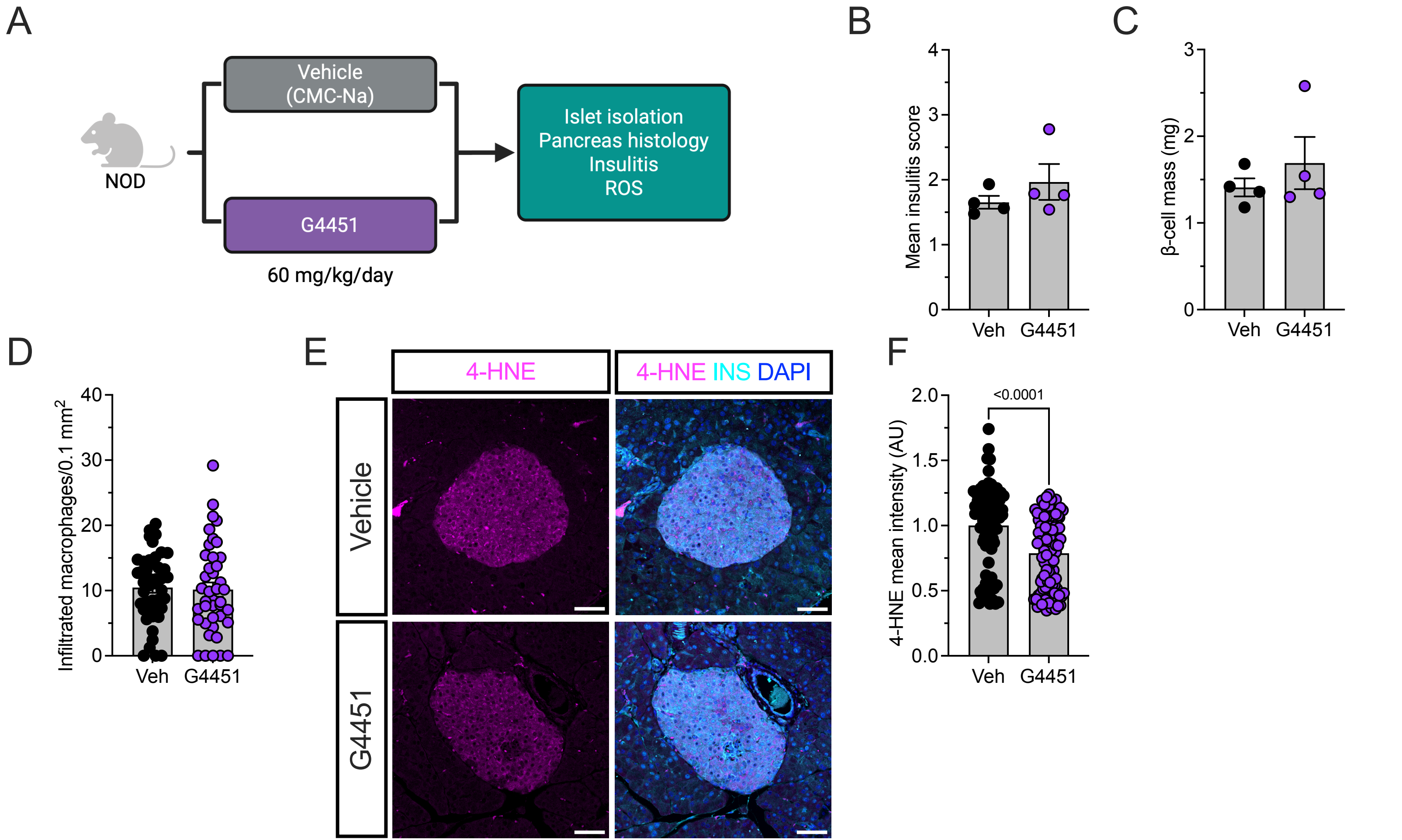

A
Vehicle
(CMC-Na)
NOD
G4451
60 mg/kg/day
Islet isolation
Pancreas histology
Insulitis
ROS
B
Mean insulitis score
Veh
G4451
C
β-cell mass (mg)
Veh
G4451
D
Infiltrated macrophages/0.1 mm2
Veh
G4451
E
4-HNE
4-HNE INS DAPI
Vehicle
G4451
F
4-HNE mean intensity (AU)
<0.0001
Veh
G4451

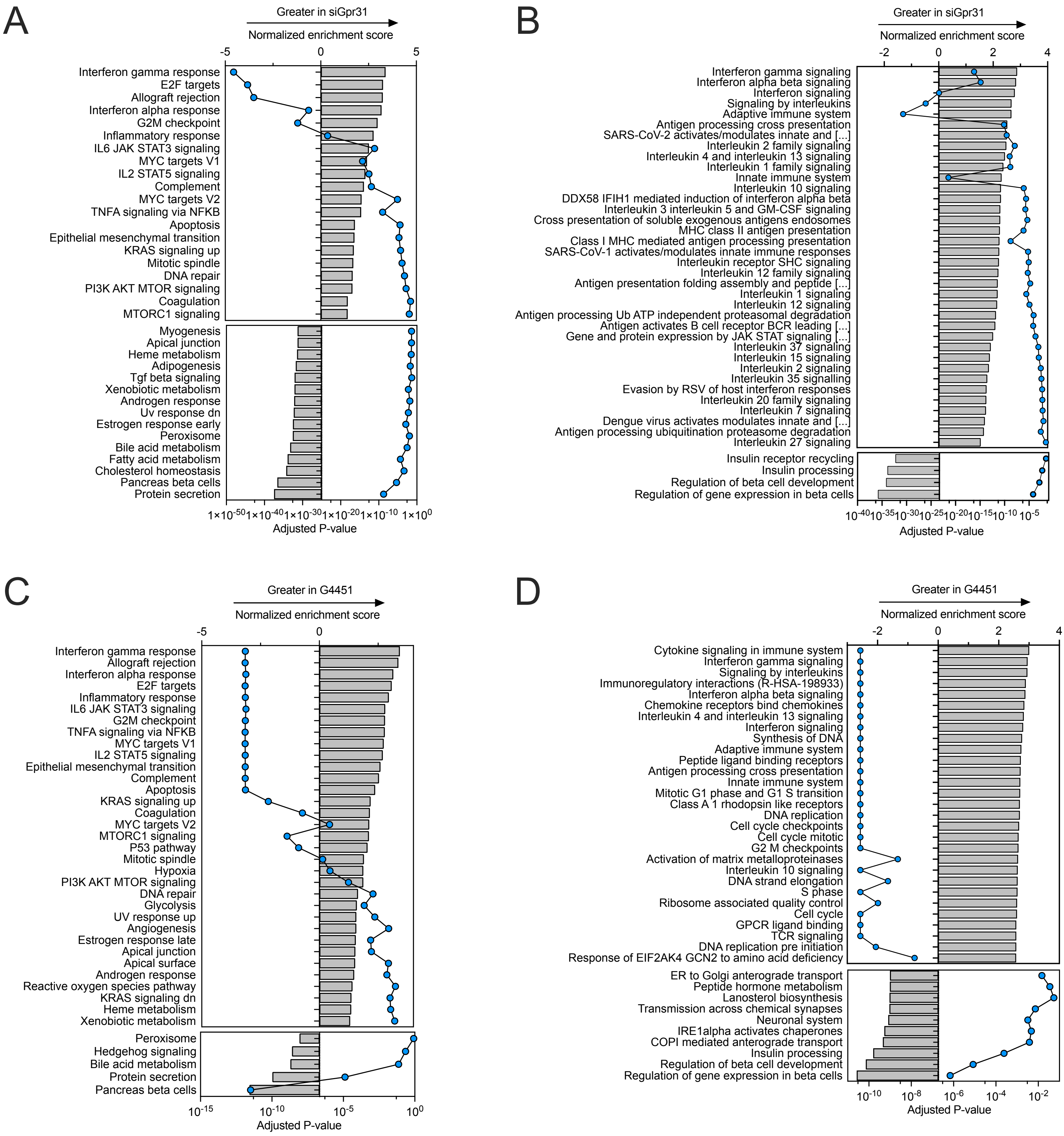
A
Greater in siGpr31
Normalized enrichment score
-5
0
5
Interferon gamma response
E2F targets
Allograft rejection
Interferon alpha response
G2M checkpoint
Inflammatory response
IL6 JAK STAT3 signaling
MYC targets V1
IL2 STAT5 signaling
Complement
MYC targets V2
TNFA signaling via NFKB
Apoptosis
Epithelial mesenchymal transition
KRAS signaling up
Mitotic spindle
DNA repair
PI3K AKT MTOR signaling
Coagulation
MTORC1 signaling
Myogenesis
Apical junction
Heme metabolism
Adipogenesis
Tgf beta signaling
Xenobiotic metabolism
Androgen response
Uv response dn
Estrogen response early
Peroxisome
Bile acid metabolism
Fatty acid metabolism
Cholesterol homeostasis
Pancreas beta cells
Protein secretion
1×10-50 1×10-40 1×10-30 1×10-20 1×10-10 1×100
Adjusted P-value
B
Greater in siGpr31
Normalized enrichment score
-2
0
2
4
Interferon gamma signaling
Interferon alpha beta signaling
Interferon signaling
Signaling by interleukins
Adaptive immune system
Antigen processing cross presentation
SARS-CoV-2 activates/modulates innate and [...]
Interleukin 2 family signaling
Interleukin 4 and interleukin 13 signaling
Interleukin 1 family signaling
Innate immune system
Interleukin 10 signaling
DDX58 IFIH1 mediated induction of interferon alpha beta
Interleukin 3 interleukin 5 and GM-CSF signaling
Cross presentation of soluble exogenous antigens endosomes
MHC class II antigen presentation
Class I MHC mediated antigen processing presentation
SARS-CoV-1 activates/modulates innate immune responses
Interleukin receptor SHC signaling
Interleukin 12 family signaling
Antigen presentation folding assembly and peptide [...]
Interleukin 1 signaling
Interleukin 12 signaling
Antigen processing Ub ATP independent proteasomal degradation
Antigen activates B cell receptor BCR leading [...]
Gene and protein expression by JAK STAT signaling [...]
Interleukin 37 signaling
Interleukin 15 signaling
Interleukin 2 signaling
Interleukin 35 signalling
Evasion by RSV of host interferon responses
Interleukin 20 family signaling
Interleukin 7 signaling
Dengue virus activates modulates innate and [...]
Antigen processing ubiquitination proteasome degradation
Interleukin 27 signaling
Insulin receptor recycling
Insulin processing
Regulation of beta cell development
Regulation of gene expression in beta cells
10-40 10-35 10-30 10-25 10-20 10-15 10-10 10-5
Adjusted P-value
C
Greater in G4451
Normalized enrichment score
-5
0
Interferon gamma response
Allograft rejection
Interferon alpha response
E2F targets
Inflammatory response
IL6 JAK STAT3 signaling
G2M checkpoint
TNFA signaling via NFKB
MYC targets V1
IL2 STAT5 signaling
Epithelial mesenchymal transition
Complement
Apoptosis
KRAS signaling up
Coagulation
MYC targets V2
MTORC1 signaling
P53 pathway
Mitotic spindle
Hypoxia
PI3K AKT MTOR signaling
DNA repair
Glycolysis
UV response up
Angiogenesis
Estrogen response late
Apical junction
Apical surface
Androgen response
Reactive oxygen species pathway
KRAS signaling dn
Heme metabolism
Xenobiotic metabolism
Peroxisome
Hedgehog signaling
Bile acid metabolism
Protein secretion
Pancreas beta cells
10-15 10-10 10-5 100
Adjusted P-value
D
Greater in G4451
Normalized enrichment score
-2
0
2
4
Cytokine signaling in immune system
Interferon gamma signaling
Signaling by interleukins
Immunoregulatory interactions (R-HSA-198933)
Interferon alpha beta signaling
Chemokine receptors bind chemokines
Interleukin 4 and interleukin 13 signaling
Interferon signaling
Synthesis of DNA
Adaptive immune system
Peptide ligand binding receptors
Antigen processing cross presentation
Innate immune system
Mitotic G1 phase and G1 S transition
Class A 1 rhodopsin like receptors
DNA replication
Cell cycle checkpoints
Cell cycle mitotic
G2 M checkpoints
Activation of matrix metalloproteinases
Interleukin 10 signaling
DNA strand elongation
S phase
Ribosome associated quality control
Cell cycle
GPCR ligand binding
TCR signaling
DNA replication pre initiation
Response of EIF2AK4 GCN2 to amino acid deficiency
ER to Golgi anterograde transport
Peptide hormone metabolism
Lanosterol biosynthesis
Transmission across chemical synapses
Neuronal system
IRE1alpha activates chaperones
COPI mediated anterograde transport
Insulin processing
Regulation of beta cell development
Regulation of gene expression in beta cells
10-10 10-8 10-6 10-4 10-2
Adjusted P-value

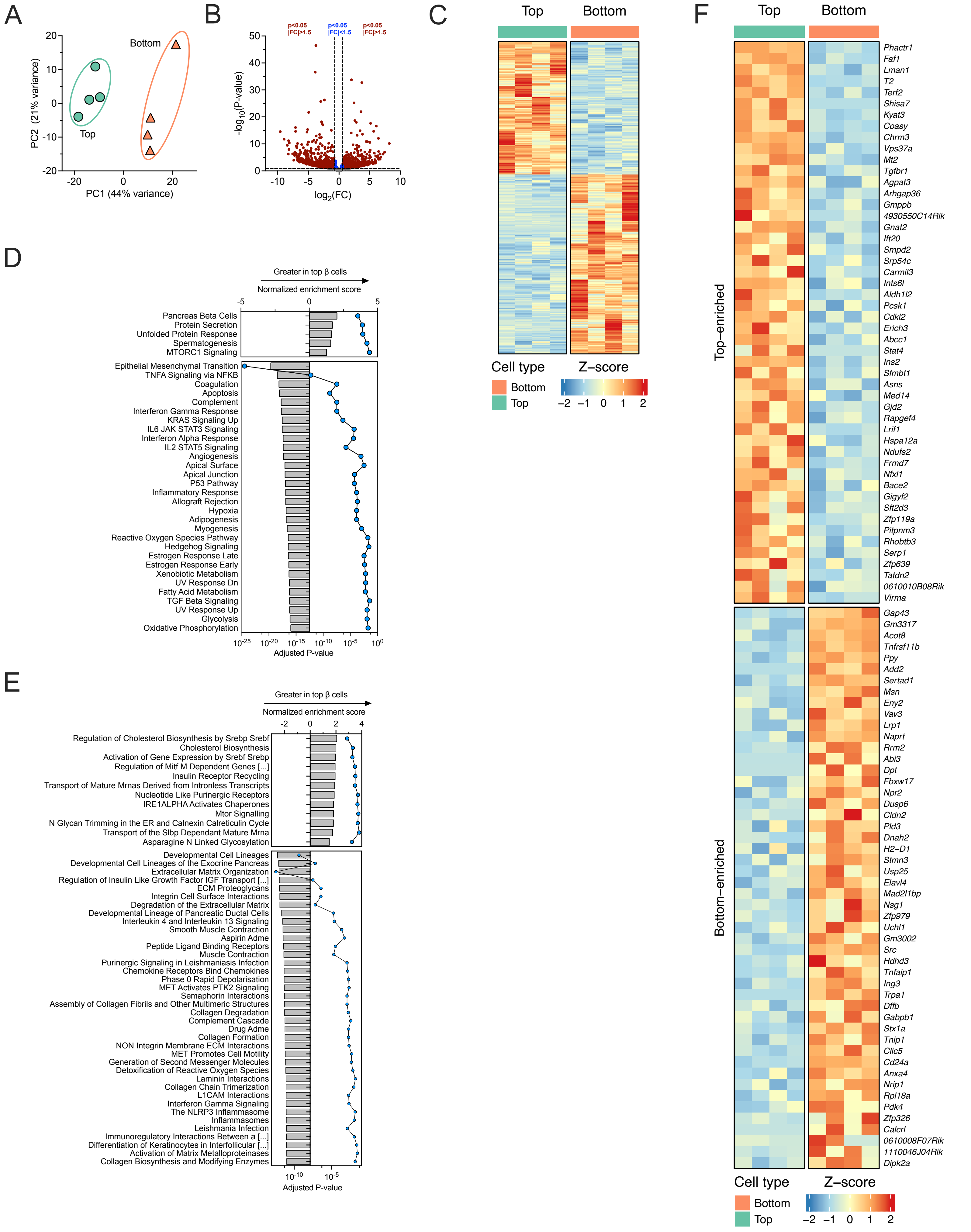
A
Bottom
Top
PC2 (21% variance)
PC1 (44% variance)
B
p<0.05 |FC|>1.5
p<0.05 |FC|<1.5
p<0.05 |FC|>1.5
-log10(P-value)
log2(FC)
C
Top
Bottom
Cell type
Bottom
Top
Z−score
D
Greater in top β cells
Normalized enrichment score
Pancreas Beta Cells
Protein Secretion
Unfolded Protein Response
Spermatogenesis
MTORC1 Signaling
Epithelial Mesenchymal Transition
TNFA Signaling via NFKB
Coagulation
Apoptosis
Complement
Interferon Gamma Response
KRAS Signaling Up
IL6 JAK STAT3 Signaling
Interferon Alpha Response
IL2 STAT5 Signaling
Angiogenesis
Apical Surface
Apical Junction
P53 Pathway
Inflammatory Response
Allograft Rejection
Hypoxia
Adipogenesis
Myogenesis
Reactive Oxygen Species Pathway
Hedgehog Signaling
Estrogen Response Late
Estrogen Response Early
Xenobiotic Metabolism
UV Response Dn
Fatty Acid Metabolism
TGF Beta Signaling
UV Response Up
Glycolysis
Oxidative Phosphorylation
Adjusted P-value
E
Greater in top β cells
Normalized enrichment score
Regulation of Cholesterol Biosynthesis by Srebp Srebf
Cholesterol Biosynthesis
Activation of Gene Expression by Srebf Srebp
Regulation of Mitf M Dependent Genes [...]
Insulin Receptor Recycling
Transport of Mature Mrnas Derived from Intronless Transcripts
Nucleotide Like Purinergic Receptors
IRE1ALPHA Activates Chaperones
Mtor Signalling
N Glycan Trimming in the ER and Calnexin Calreticulin Cycle
Transport of the Slbp Dependant Mature Mrna
Asparagine N Linked Glycosylation
Developmental Cell Lineages
Developmental Cell Lineages of the Exocrine Pancreas
Extracellular Matrix Organization
Regulation of Insulin Like Growth Factor IGF Transport [...]
ECM Proteoglycans
Integrin Cell Surface Interactions
Degradation of the Extracellular Matrix
Developmental Lineage of Pancreatic Ductal Cells
Interleukin 4 and Interleukin 13 Signaling
Smooth Muscle Contraction
Aspirin Adme
Peptide Ligand Binding Receptors
Muscle Contraction
Purinergic Signaling in Leishmaniasis Infection
Chemokine Receptors Bind Chemokines
Phase 0 Rapid Depolarisation
MET Activates PTK2 Signaling
Semaphorin Interactions
Assembly of Collagen Fibrils and Other Multimeric Structures
Collagen Degradation
Complement Cascade
Drug Adme
Collagen Formation
NON Integrin Membrane ECM Interactions
MET Promotes Cell Motility
Generation of Second Messenger Molecules
Detoxification of Reactive Oxygen Species
Laminin Interactions
Collagen Chain Trimerization
L1CAM Interactions
Interferon Gamma Signaling
The NLRP3 Inflammasome
Inflammasomes
Leishmania Infection
Immunoregulatory Interactions Between a [...]
Differentiation of Keratinocytes in Interfollicular [...]
Activation of Matrix Metalloproteinases
Collagen Biosynthesis and Modifying Enzymes
Adjusted P-value
F
Top
Bottom
Top−enriched
Phactr1
Faf1
Lman1
T2
Terf2
Shisa7
Kyat3
Coasy
Chrm3
Vps37a
Mt2
Tgfbr1
Agpat3
Arhgap36
Gmppb
4930550C14Rik
Gnat2
Ift20
Smpd2
Srp54c
Carmil3
Ints6l
Aldh1l2
Pcsk1
Cdkl2
Erich3
Abcc1
Stat4
Ins2
Sfmbt1
Asns
Med14
Gjd2
Rapgef4
Lrif1
Hspa12a
Ndufs2
Frmd7
Nfxl1
Bace2
Gigyf2
Sft2d3
Zfp119a
Pitpnm3
Rhobtb3
Serp1
Zfp639
Tatdn2
0610010B08Rik
Virma
Bottom−enriched
Gap43
Gm3317
Acot8
Tnfrsf11b
Ppy
Add2
Sertad1
Msn
Eny2
Vav3
Lrp1
Naprt
Rrm2
Abi3
Dpt
Fbxw17
Npr2
Dusp6
Cldn2
Pld3
Dnah2
H2−D1
Stmn3
Usp25
Elavl4
Mad2l1bp
Nsg1
Zfp979
Uchl1
Gm3002
Src
Hdhd3
Tnfaip1
Ing3
Trpa1
Dffb
Gabpb1
Stx1a
Tnip1
Clic5
Cd24a
Anxa4
Nrip1
Rpl18a
Pdk4
Zfp326
Calcrl
0610008F07Rik
1110046J04Rik
Dipk2a
Cell type
Bottom
Top
Z−score

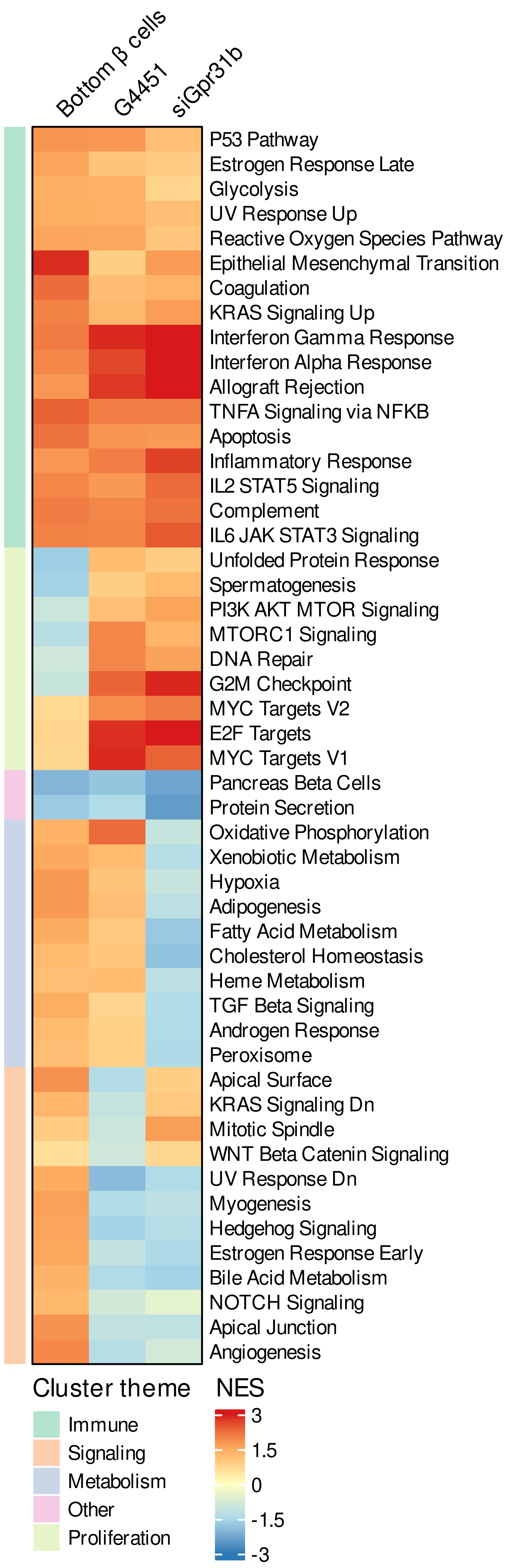
Bottom β cells
G4451
siGpr31b
P53 Pathway
Estrogen Response Late
Glycolysis
UV Response Up
Reactive Oxygen Species Pathway
Epithelial Mesenchymal Transition
Coagulation
KRAS Signaling Up
Interferon Gamma Response
Interferon Alpha Response
Allograft Rejection
TNFA Signaling via NFKB
Apoptosis
Inflammatory Response
IL2 STAT5 Signaling
Complement
IL6 JAK STAT3 Signaling
Unfolded Protein Response
Spermatogenesis
PI3K AKT MTOR Signaling
MTORC1 Signaling
DNA Repair
G2M Checkpoint
MYC Targets V2
E2F Targets
MYC Targets V1
Pancreas Beta Cells
Protein Secretion
Oxidative Phosphorylation
Xenobiotic Metabolism
Hypoxia
Adipogenesis
Fatty Acid Metabolism
Cholesterol Homeostasis
Heme Metabolism
TGF Beta Signaling
Androgen Response
Peroxisome
Apical Surface
KRAS Signaling Dn
Mitotic Spindle
WNT Beta Catenin Signaling
UV Response Dn
Myogenesis
Hedgehog Signaling
Estrogen Response Early
Bile Acid Metabolism
NOTCH Signaling
Apical Junction
Angiogenesis
Cluster theme
Immune
Signaling
Metabolism
Other
Proliferation
NES
3
1.5
0
-1.5
-3

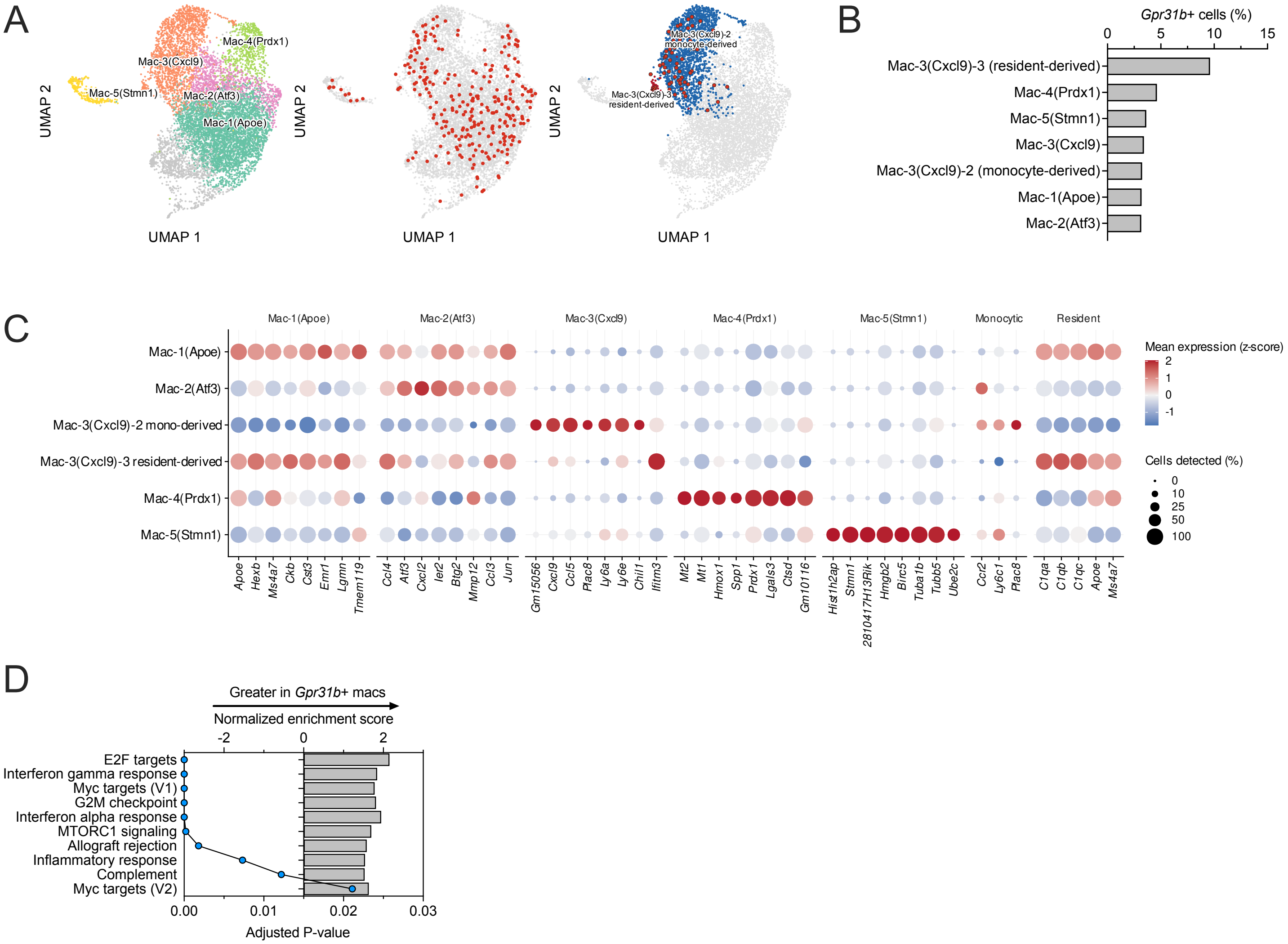
A
Mac-4(Prdx1)
Mac-3(Cxcl9)
Mac-5(Stmn1)
Mac-2(Atf3)
Mac-1(Apoe)
UMAP 2
UMAP 1
Mac-3(Cxcl9)-2 monocyte-derived
Mac-3(Cxcl9)-3 resident-derived
B
Gpr31b+ cells (%)
0 5 10 15
Mac-3(Cxcl9)-3 (resident-derived)
Mac-4(Prdx1)
Mac-5(Stmn1)
Mac-3(Cxcl9)
Mac-3(Cxcl9)-2 (monocyte-derived)
Mac-1(Apoe)
Mac-2(Atf3)
C
Mac-1(Apoe) Mac-2(Atf3) Mac-3(Cxcl9) Mac-4(Prdx1) Mac-5(Stmn1) Monocytic Resident
Mac-1(Apoe)
Mac-2(Atf3)
Mac-3(Cxcl9)-2 mono-derived
Mac-3(Cxcl9)-3 resident-derived
Mac-4(Prdx1)
Mac-5(Stmn1)
Apoe Hexb Ms4a7 Ckb Cst3 Emr1 Lgmn Tmem119
Ccl4 Atf3 Cxcl2 Ier2 Btg2 Mmp12 Ccl3 Jun
Gm15056 Cxcl9 Ccl5 Plac8 Ly6a Ly6e Chil1 Ifitm3
Mt2 Mt1 Hmox1 Spp1 Prdx1 Lgals3 Ctsd Gm10116
Hist1h2ap Stmn1 2810417H13Rik Hmgb2 Birc5 Tuba1b Tubb5 Ube2c
Ccr2 Ly6c1 Plac8
C1qa C1qb C1qc Apoe Ms4a7
Mean expression (z-score)
2 1 0 -1
Cells detected (%)
0 10 25 50 100
D
Greater in Gpr31b+ macs
Normalized enrichment score
-2 0 2
E2F targets
Interferon gamma response
Myc targets (V1)
G2M checkpoint
Interferon alpha response
MTORC1 signaling
Allograft rejection
Inflammatory response
Complement
Myc targets (V2)
0.00 0.01 0.02 0.03
Adjusted P-value

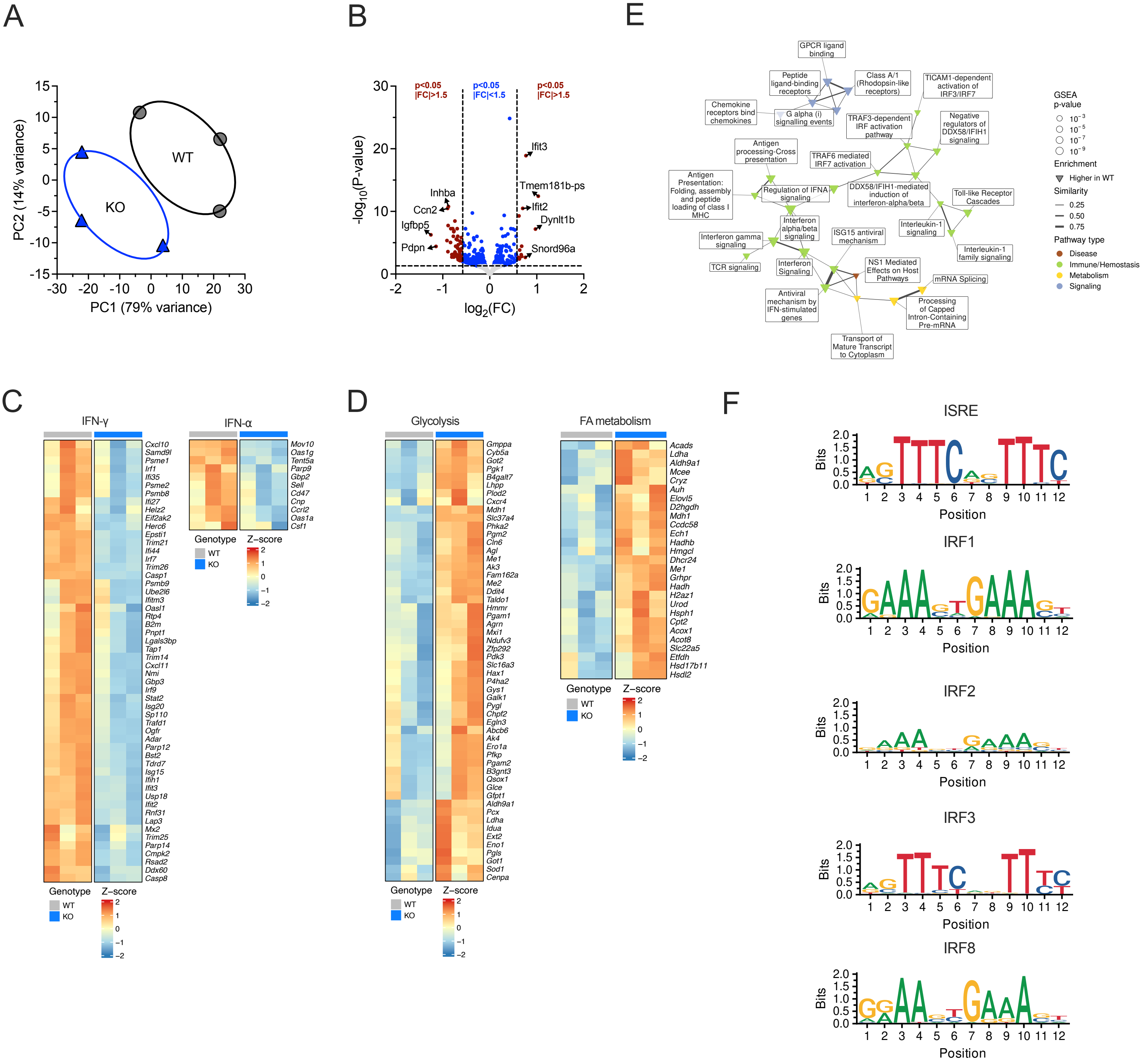
A
PC2 (14% variance)
PC1 (79% variance)
WT
KO
B
-log10(P-value)
log2(FC)
p<0.05
|FC|>1.5
p<0.05
|FC|<1.5
p<0.05
|FC|>1.5
Ifit3
Tmem181b-ps
Ifit2
Dynlt1b
Snord96a
Inhba
Ccn2
Igfbp5
Pdpn
E
GPCR ligand binding
Peptide ligand-binding receptors
Class A/1 (Rhodopsin-like receptors)
TICAM1-dependent activation of IRF3/IRF7
Chemokine receptors bind chemokines
G alpha (i) signalling events
TRAF3-dependent IRF activation pathway
Negative regulators of DDX58/IFIH1 signaling
Antigen processing-Cross presentation
TRAF6 mediated IRF7 activation
Antigen Presentation: Folding, assembly and peptide loading of class I MHC
Regulation of IFNA signaling
DDX58/IFIH1-mediated induction of interferon-alpha/beta
Toll-like Receptor Cascades
Interferon alpha/beta signaling
Interleukin-1 signaling
Interferon gamma signaling
ISG15 antiviral mechanism
Interleukin-1 family signaling
TCR signaling
Interferon Signaling
NS1 Mediated Effects on Host Pathways
mRNA Splicing
Antiviral mechanism by IFN-stimulated genes
Processing of Capped Intron-Containing Pre-mRNA
Transport of Mature Transcript to Cytoplasm
GSEA p-value
Enrichment
Higher in WT
Similarity
0.25
0.50
0.75
Pathway type
Disease
Immune/Hemostasis
Metabolism
Signaling
C
IFN-γ
Cxcl10 Samd9l Psme1 Irf1 Ifi35 Psme2 Psmb8 Ifi27 Helz2 Eif2ak2 Herc6 Epsti1 Trim21 Ifi44 Irf7 Trim26 Casp1 Psmb9 Ube2l6 Ifitm3 Oasl1 Rtp4 B2m Pnpt1 Lgals3bp Tap1 Trim14 Cxcl11 Nmi Gbp3 Irf9 Stat2 Isg20 Sp110 Trafd1 Ogfr Adar Parp12 Bst2 Tdrd7 Isg15 Ifih1 Ifit3 Usp18 Ifit2 Rnf31 Lap3 Mx2 Trim25 Parp14 Cmpk2 Rsad2 Ddx60 Casp8
IFN-α
Mov10 Oas1g Tent5a Parp9 Gbp2 Sell Cd47 Cnp Ccrl2 Oas1a Csf1
Genotype
WT
KO
Z-score
D
Glycolysis
Gmppa Cyb5a Got2 Pgk1 B4galt7 Lhpp Plod2 Cxcr4 Mdh1 Slc37a4 Phka2 Pgm2 Cln6 Agl Me1 Ak3 Fam162a Me2 Ddit4 Taldo1 Hmmr Pgam1 Agrn Mxi1 Ndufv3 Zfp292 Pdk3 Slc16a3 Hax1 P4ha2 Gys1 Galk1 Pygl Chpf2 Egln3 Abcb6 Ak4 Ero1a Pfkp Pgam2 B3gnt3 Qsox1 Glce Gfpt1 Aldh9a1 Pcx Ldha Idua Ext2 Eno1 Pgls Got1 Sod1 Cenpa
FA metabolism
Acads Ldha Aldh9a1 Mcee Cryz Auh Elovl5 D2hgdh Mdh1 Ccdc58 Ech1 Hadhb Hmgcl Dhcr24 Me1 Grhpr Hadh H2az1 Urod Hsph1 Cpt2 Acox1 Acot8 Slc22a5 Etfdh Hsd17b11 Hsdl2
F
ISRE
IRF1
IRF2
IRF3
IRF8
Bits
Position

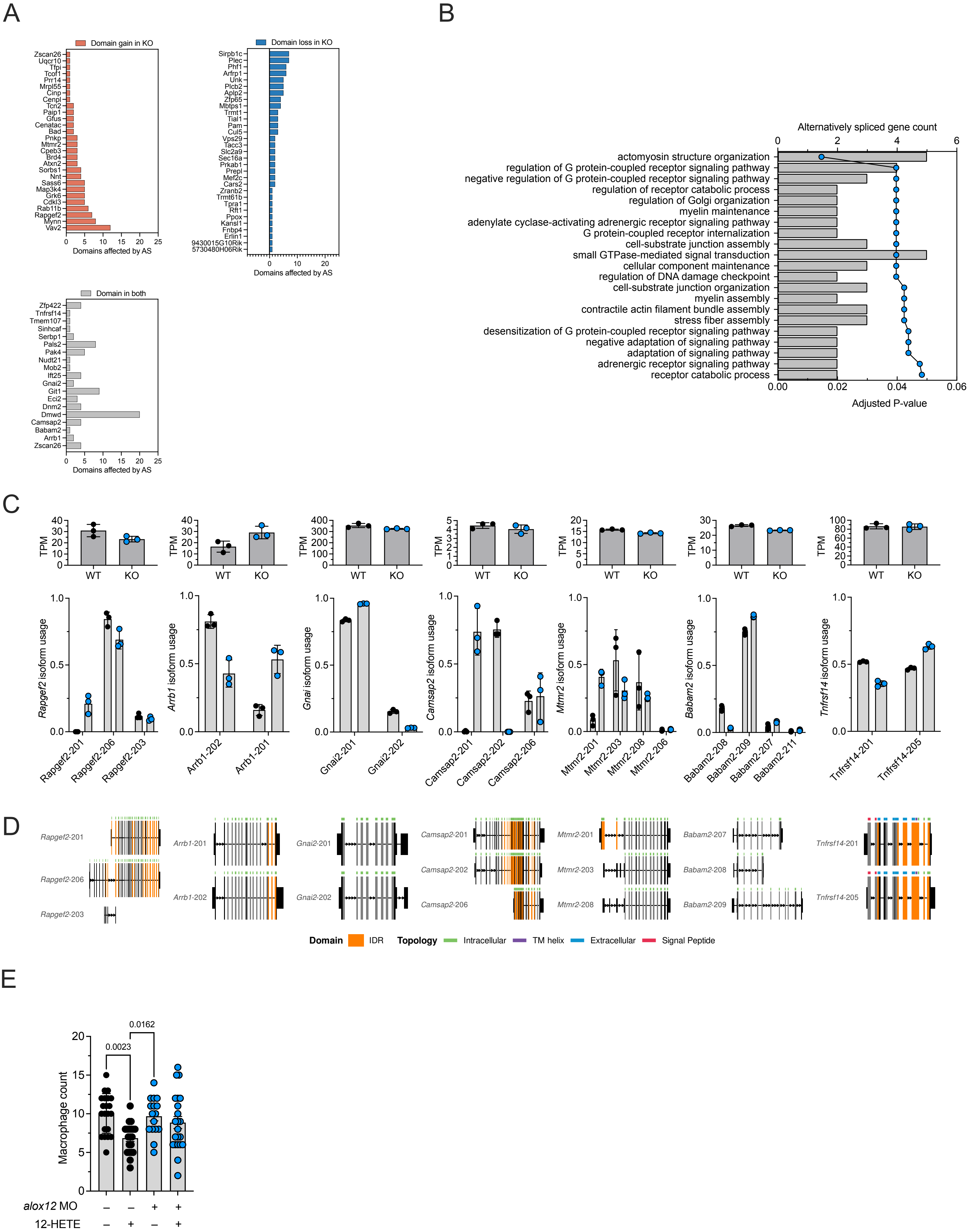
A
Domain gain in KO
Zscan26
Uqcr10
Tfpi
Tcof1
Prr14
Mrpl55
Cinp
Cenpl
Tcn2
Paip1
Gfus
Cenatac
Bad
Pnkp
Mtmr2
Cpeb3
Brd4
Atxn2
Sorbs1
Nnt
Sass6
Map3k4
Grk6
Cdkl3
Rab11b
Rapgef2
Mynn
Vav2
Domains affected by AS
Domain loss in KO
Sirpb1c
Plec
Phf1
Arfrp1
Unk
Plcb2
Aplp2
Zfp65
Mbtps1
Trmt1
Tial1
Pam
Cul5
Vps29
Tacc3
Slc2a9
Sec16a
Prkab1
Prepl
Mef2c
Cars2
Zranb2
Trmt61b
Tpra1
Rft1
Ppox
Kansl1
Fnbp4
Erlin1
9430015G10Rik
5730480H06Rik
Domains affected by AS
Domain in both
Zfp422
Tnfrsf14
Tmem107
Sinhcaf
Serbp1
Pals2
Pak4
Nudt21
Mob2
Ift25
Gnai2
Git1
Eci2
Dnm2
Dmwd
Camsap2
Babam2
Arrb1
Zscan26
Domains affected by AS
B
Alternatively spliced gene count
actomyosin structure organization
regulation of G protein-coupled receptor signaling pathway
negative regulation of G protein-coupled receptor signaling pathway
regulation of receptor catabolic process
regulation of Golgi organization
myelin maintenance
adenylate cyclase-activating adrenergic receptor signaling pathway
G protein-coupled receptor internalization
cell-substrate junction assembly
small GTPase-mediated signal transduction
cellular component maintenance
regulation of DNA damage checkpoint
cell-substrate junction organization
myelin assembly
contractile actin filament bundle assembly
stress fiber assembly
desensitization of G protein-coupled receptor signaling pathway
negative adaptation of signaling pathway
adaptation of signaling pathway
adrenergic receptor signaling pathway
receptor catabolic process
Adjusted P-value
C
TPM
WT
KO
Rapgef2 isoform usage
Rapgef2-201
Rapgef2-206
Rapgef2-203
Arrb1 isoform usage
Arrb1-202
Arrb1-201
Gnai isoform usage
Gnai2-201
Gnai2-202
Camsap2 isoform usage
Camsap2-201
Camsap2-202
Camsap2-206
Mtmr2 isoform usage
Mtmr2-201
Mtmr2-203
Mtmr2-208
Mtmr2-206
Babam2 isoform usage
Babam2-208
Babam2-209
Babam2-207
Babam2-211
Tnfrsf14 isoform usage
Tnfrsf14-201
Tnfrsf14-205
D
Rapgef2-201
Rapgef2-206
Rapgef2-203
Arrb1-201
Arrb1-202
Gnai2-201
Gnai2-202
Camsap2-201
Camsap2-202
Camsap2-206
Mtmr2-201
Mtmr2-203
Mtmr2-208
Babam2-207
Babam2-208
Babam2-209
Tnfrsf14-201
Tnfrsf14-205
Domain IDR Topology Intracellular TM helix Extracellular Signal Peptide
E
0.0023
0.0162
Macrophage count
alox12 MO
12-HETE

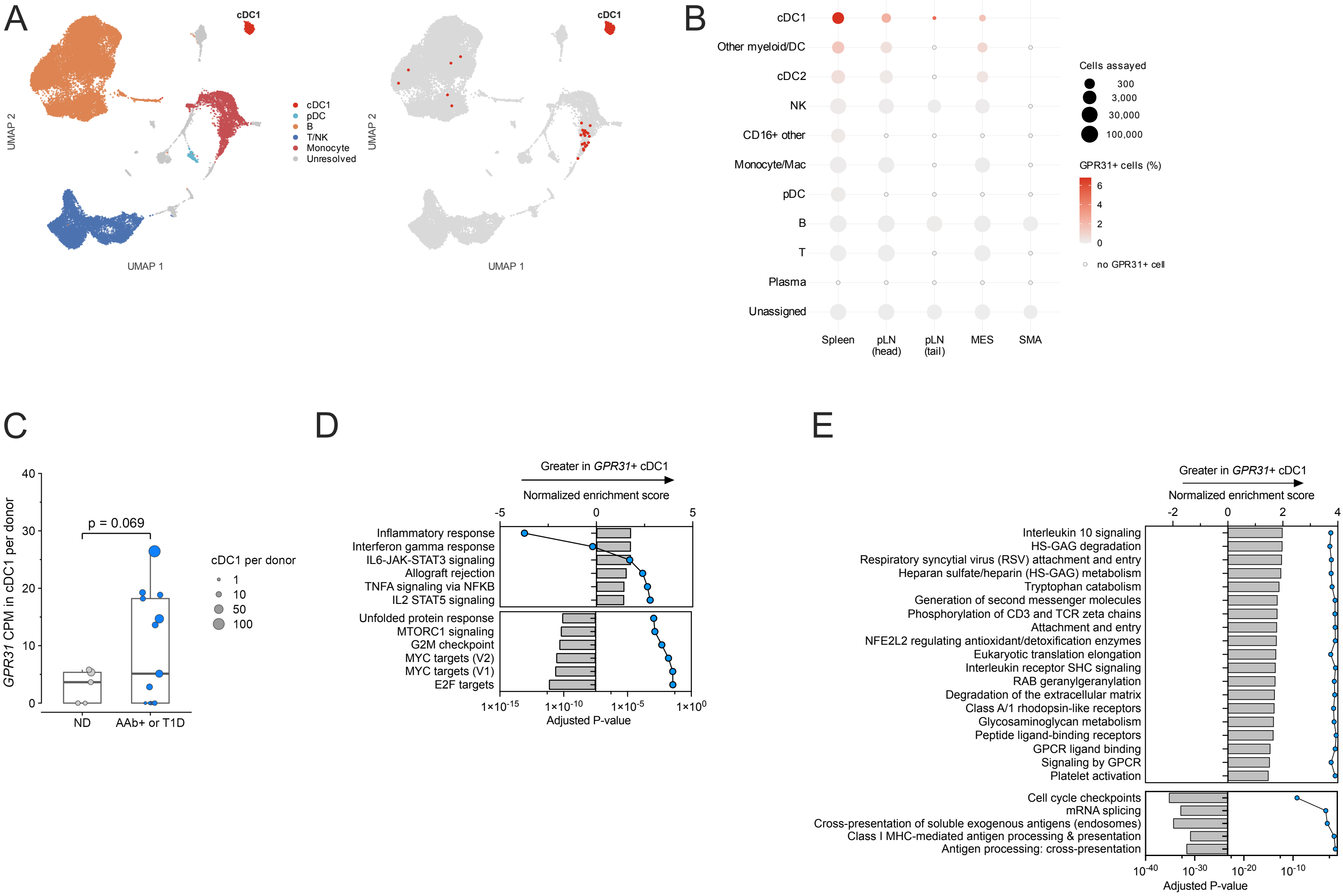

A
cDC1
cDC1
pDC
B
T/NK
Monocyte
Unresolved
UMAP 2
UMAP 1
B
cDC1
Other myeloid/DC
cDC2
NK
CD16+ other
Monocyte/Mac
pDC
B
T
Plasma
Unassigned
Spleen
pLN (head)
pLN (tail)
MES
SMA
Cells assayed
300
3,000
30,000
100,000
GPR31+ cells (%)
6
4
2
0
no GPR31+ cell
C
GPR31 CPM in cDC1 per donor
p = 0.069
ND
AAb+ or T1D
cDC1 per donor
1
10
50
100
D
Greater in GPR31+ cDC1
Normalized enrichment score
Inflammatory response
Interferon gamma response
IL6-JAK-STAT3 signaling
Allograft rejection
TNFA signaling via NFKB
IL2 STAT5 signaling
Unfolded protein response
MTORC1 signaling
G2M checkpoint
MYC targets (V2)
MYC targets (V1)
E2F targets
Adjusted P-value
E
Greater in GPR31+ cDC1
Normalized enrichment score
Interleukin 10 signaling
HS-GAG degradation
Respiratory syncytial virus (RSV) attachment and entry
Heparan sulfate/heparin (HS-GAG) metabolism
Tryptophan catabolism
Generation of second messenger molecules
Phosphorylation of CD3 and TCR zeta chains
Attachment and entry
NFE2L2 regulating antioxidant/detoxification enzymes
Eukaryotic translation elongation
Interleukin receptor SHC signaling
RAB geranylgeranylation
Degradation of the extracellular matrix
Class A/1 rhodopsin-like receptors
Glycosaminoglycan metabolism
Peptide ligand-binding receptors
GPCR ligand binding
Signaling by GPCR
Platelet activation
Cell cycle checkpoints
mRNA splicing
Cross-presentation of soluble exogenous antigens (endosomes)
Class I MHC-mediated antigen processing & presentation
Antigen processing: cross-presentation
Adjusted P-value
